# Deep neural network automated segmentation of cellular structures in volume electron microscopy

**DOI:** 10.1101/2022.08.02.502534

**Authors:** Benjamin Gallusser, Giorgio Maltese, Giuseppe Di Caprio, Tegy John Vadakkan, Anwesha Sanyal, Elliott Somerville, Mihir Sahasrabudhe, Justin O’Connor, Martin Weigert, Tom Kirchhausen

## Abstract

Recent advances in automated segmentation using deep neural network models allow identification of intracellular structures. This study describes a new pipeline to train a convolutional neural network for rapid and efficient detection of structures of wide range in size and complexity.

**Abstract:** Three-dimensional electron microscopy is an important imaging modality in contemporary cell biology. Identification of intracellular structures is laborious and time-consuming, however, and impairs effective use of a potentially powerful tool. Resolving this bottleneck is therefore a critical next step in frontier biomedical imaging. We describe ***A****utomated **S**egmentation of intracellular substructures in **E**lectron **M**icroscopy* (ASEM), a new pipeline to train a convolutional neural network to detect structures of wide range in size and complexity. We obtain for each structure a dedicated model based on a small number of sparsely annotated ground truth annotations from only one or two cells. To improve model generalization to different imaging conditions, we developed a rapid, computationally effective strategy to refine an already trained model by including a few additional annotations. We show the successful automated identification of mitochondria, Golgi apparatus, endoplasmic reticulum, nuclear pore complexes, caveolae, clathrin coated pits and coated vesicles in cells imaged by focused ion beam scanning electron microscopy with quasi-isotropic resolution. ASEM enabled us to uncover a wide range of membrane-nuclear pore diameters within a single cell and to derive morphological metrics from clathrin coated pits and vesicles at all stages of maturation consistent with the classical constant-growth assembly model.

## Introduction

Three-dimensional, high-resolution imaging provides a snapshot of the internal organization of a cell at a defined time point and in a defined physiological state. Focused ion beam scanning electron microscopy (FIB-SEM) yields nearly isotropic, nanometer-level resolution, three-dimensional images by sequential imaging of the surface layer of a sample, which is then etched away with an ion beam to reveal the layer beneath (Knott et al., 2008; Xu et al., 2017). FIB-SEM technology continues to develop, and it can be a particularly valuable contemporary tool for imaging the complete volume of a cell, but segmentation of the three-dimensional data sets and subsequent analysis of the results are substantial hurdles, as the images are far too large to interpret by inspection (Heinrich et al., 2021).

The widespread success of machine learning in bioimage informatics has recently inspired the application of deep-learning approaches to automated segmentation. Examples using deep convolutional networks for data with anisotropic resolution include DeepEM3D (Zeng et al., 2017) and CDeep-3M (Haberl et al., 2018), for segmentation of mitochondria and Golgi apparatus with extensive post-processing (Mekuč et al., 2020, 2022), as well as cell-organelle segmentation in quasi-isotropic FIB-SEM data of beta cells (Müller et al., 2020). A pipeline created by the COSEM project (Heinrich et al., 2021) enables automated whole-cell simultaneous segmentation of up to 35 organelles from relatively sparse but very precise 3D ground truth annotations from FIB-SEM data of cells prepared by high pressure freezing and freeze substitution (HPFS), obtained at 3-5 nm voxel size with approximately isotropic resolution. The most common strategy used by the COSEM project involved training with ground truth annotations from multiple classes of objects at the same time, typically at a high computational cost (500,000 or more training iterations) (Heinrich et al., 2021).

The current approaches all suffer from a demand for substantial computational resources, and they generally require a large set of precise manual annotations. Both requirements limit their practical applicability. We describe here the development and use of a new deep learning pipeline called ***A****utomated **S**egmentation of intracellular substructures in **E**lectron **M**icroscopy* (ASEM), which can detect structures of a wide range in size and complexity using deep neural networks trained on a limited number of loosely marked i.e., not necessarily pixel-precise, ground truth annotations. ASEM includes a semi-automated graph-cut procedure we developed to assist in the tedious task of ground truth preparation and a computationally efficient transfer-learning approach with a fine-tuning protocol that can be used without the need for high-end specialized CPU/GPU workstations.

We illustrate here the utility of ASEM by describing the results of its application to data from several types of cells, including FIB-SEM datasets made publicly available by the COSEM Project (Heinrich et al., 2021). We note that while cellular samples have traditionally been processed by chemical fixation (CF) and staining at room temperature, HPFS at cryogenic temperatures (as was the case for the COSEM Project data) yields a substantial increase in the preservation of many complex cellular features. We applied ASEM to three-dimensional FIB-SEM images of cells prepared by either CF or HPFS. We validated our approach by segmenting mitochondria, endoplasmic reticulum (ER) and Golgi apparatus, as these organelles had been studied previously in similar efforts (Mekuč et al., 2020, 2022)(Heinrich et al., 2021; Liu et al., 2020), and then used ASEM to recognize much smaller structures, nuclear pores and clathrin coated pits and vesicles. For nuclear pores in interphase, we can segment nearly all the pores in the nuclear membrane. We can therefore directly analyze the range of membrane-pore diameters, even for a single cell in a particular physiological state. For clathrin coated pits, we show that a relatively restricted training set leads to accurate segmentation of coated pits at all stages of their maturation as well as coated vesicles, the final step after fission from the originating membrane, and we can derive morphological metrics consistent with the classical constant-growth assembly model (Ehrlich et al., 2004; Kirchhausen, 1993, 2009; Willy et al., 2021).

All datasets, code and models are open-source https://open.quiltdata.com/b/asem-project, so that other users working with images acquired with the same or somewhat different imaging conditions can generate their own predictive models and benefit from our pre-trained models, either directly or by adapting them by fine-tuning, without the need for specialized CPU/GPU workstations.

## Results

### FIB-SEM imaging of cells

We obtained three-dimensional focused ion beam scanning electron microscopy (FIB-SEM) data sets for different types of adherent mammalian cells grown in culture (Table S1). The samples we imaged were prepared either by conventional chemical fixation and staining with osmium and uranyl acetate at room temperature (CF) or by fixation and similar staining at very low temperature using high pressure freezing and freeze substitution (HPFS), a protocol that substantially increases sample preservation (Hoffman et al., 2020; Studer et al., 2008; Xu et al., 2021). To image a volume of a cell, we used a block-face crossbeam FIB-SEM with nominal isotropic resolution of 5 or 10 nm per voxel; each image stack, obtained during 1-2 days of continuous FIB-SEM operation, was about 15-20 GB in size and contained ∼ 2000 registered sequential TIFF files and spanned a volume of roughly 2000^3 voxels corresponding to large parts of each cell. These volume datasets were used to train the deep learning pipeline for automated segmentation of intracellular structures and to explore the effects of different fixation and staining procedures on the outcome of the segmentation tasks.

We also tested the performance of our deep learning models with a small number of FIB-SEM images from HPFS preparations of complete cells (Xu et al., 2021), obtained from the publicly available OpenOrganelle initiative (Heinrich et al., 2021) (Xu et al., 2021) (Table S1). They were acquired by the COSEM team at Janelia Research Campus at a nominal resolution of 4 x 4 x 3-5 nm per voxel with a custom-modified FIB-SEM as part of their concurrent efforts to develop methodology for automated organelle segmentation aided by deep learning.

As described below, using the specific models generated with our deep learning pipeline (Fig. 1), we could reliably identify intracellular structures ranging in size and complexity from mitochondria, endoplasmic reticulum, Golgi apparatus to nuclear pores, clathrin coated pits, coated vesicles, and caveolae.

**Figure 1.**
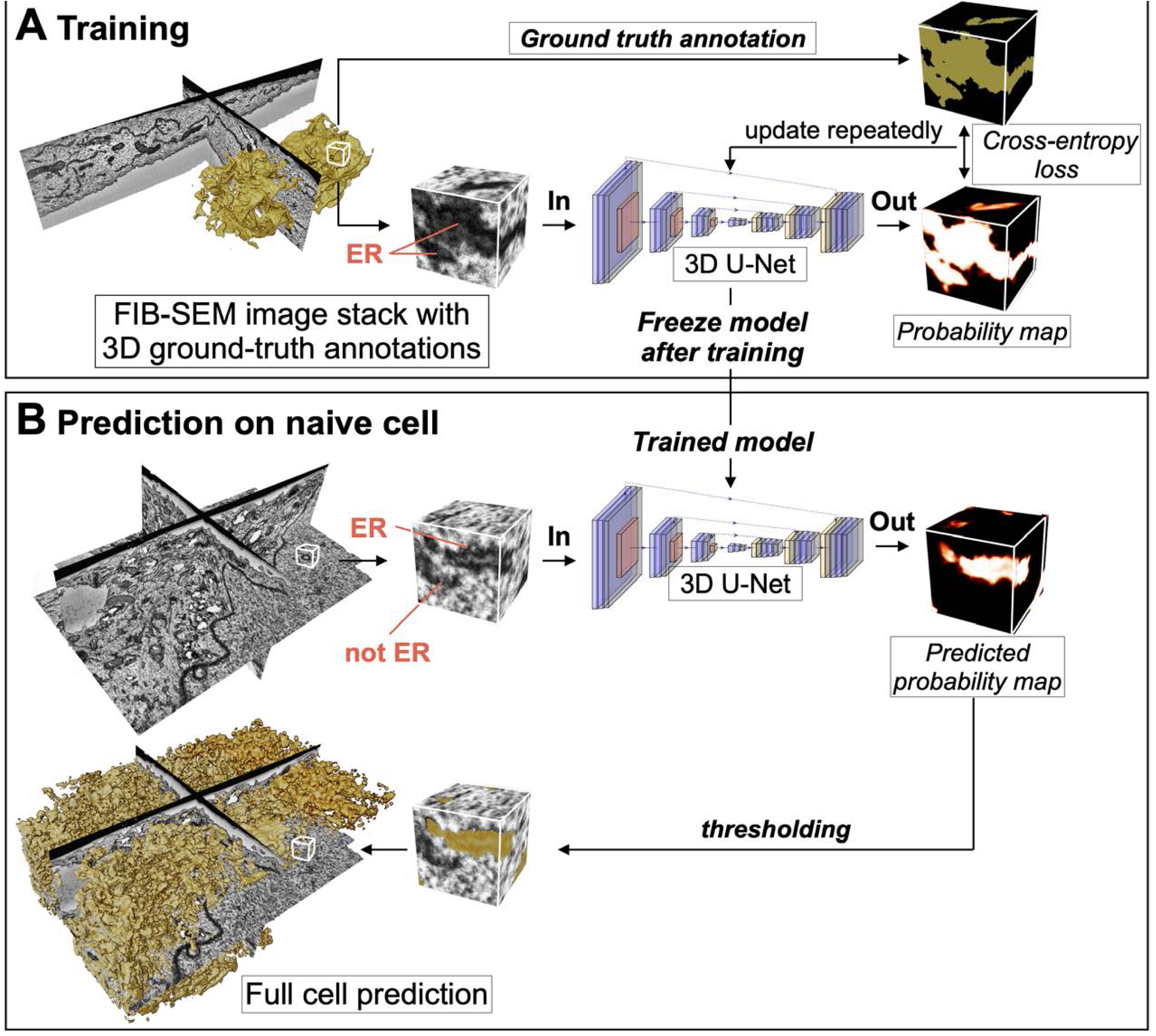
Pipelines used for training and deep-learning neural network prediction. Schematic representation of the deep-learning approach for recognizing intracellular structures in FIB-SEM volume images using a 3D U-net encoder-decoder neural network. **(A)** For training, three-dimensional stacks containing FIB-SEM data, augmented as described in Methods, are provided as input images to the 3D U-Net; in this example, the stack includes a limited number of three-dimensional ground truth annotations for the ER in the form of binary masks (yellow). The ER predicted by the 3D U-net model is a probability 3D map, whose accuracy is evaluated as a prediction error by measuring the cross-entropy loss. The model parameters are iteratively updated during training until convergence of the probability error is achieved. **(B)** For prediction, small three-dimensional stacks with data not used for training covering the complete FIB-SEM volume image of a naïve cell (or from the remaining regions of the cell used for training) are provided as input to the 3D U-net model trained in **(A)**. In this example, the predicted ER is a three-dimensional probability map centered on the three-dimensional stack. This arrangement standardizes the three-dimensional context for all predicted voxels within the FIB-SEM image.

**Figure 2.**
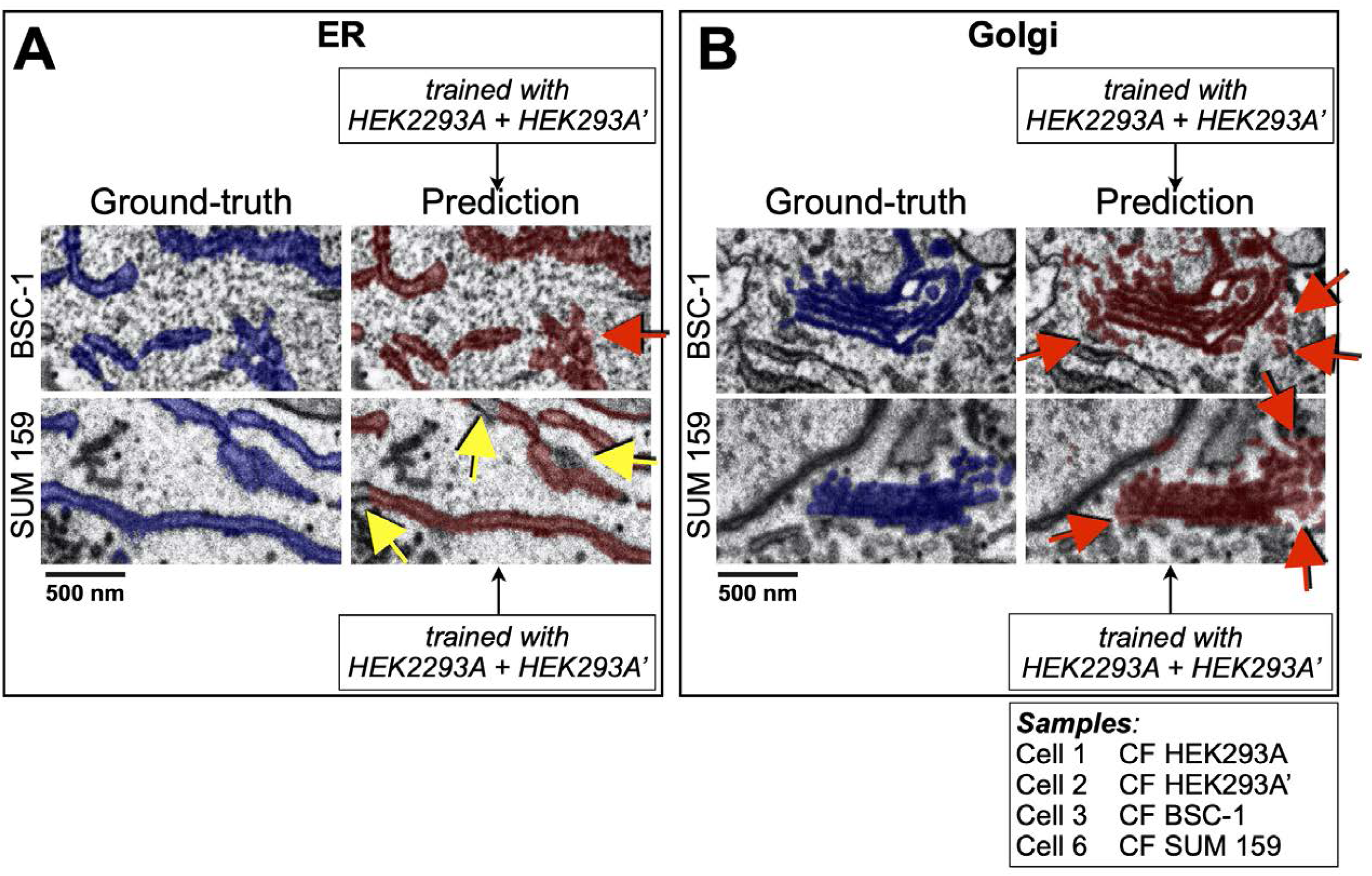
Performance of the deep-learning network to predict in naïve cells. Visual comparisons between predictions (crimson) by 3D U-net models trained using combined data from two HEK293A cells to recognize **(A)** ER or **(B)** Golgi apparatus and corresponding ground truth annotations (purple) in the naïve BSC-1 and SUM 159 cells not used for training (Table S1). The representative images from single plane views from FIB-SEM volume data are from cells prepared by CF isotropically acquired at 5 nm resolution; red and yellow arrows highlight small regions containing voxels of false positive and false negative assignments. Scale bar, 500 nm (See Videos 1 and 2).

### Ground truth annotation

The first step in most common machine-learning segmentation procedure is to create pixelwise “ground truth” annotations -- to be used for training a specific segmentation model. In the present work, we used a modest number of manual annotated segmentations of the intracellular structure of interest (see Methods for details). These segmentations came from arbitrarily chosen, diverse, and not necessarily overlapping regions from one or more cells (Table S2). As the contrast and textural appearance of raw FIB-SEM images can vary substantially due to sample preparation and imaging conditions, we make heavy use of augmentations, a technique commonly used to train deep neural networks (Shorten and Khoshgoftaar, 2019). We applied a series of randomized transformations to each of the manual segmentations to generate a set of the required size (see Methods and Table S3). The various applications described here validate this approach, which greatly reduces the manual annotation effort and makes broad application feasible.

We obtained ground truth annotations for mitochondria and Golgi apparatus, portions of endoplasmic reticulum (ER), ten endocytic clathrin coated pits at the plasma membrane, and eight nuclear pores on the nuclear envelope (Table S4). We annotated mitochondria using the carving module in Ilastik (Berg et al., 2019) and if required, edited the annotation manually using VAST (Berger et al., 2018), a volume annotation and segmentation tool for manual and semi-automatic labeling of large 3D image stacks (see example in Fig. S1 and Video 1). We annotated the more complex Golgi apparatus and ER with a dedicated graph cut-assisted, semi-automated annotation tool we developed and describe in Methods that accelerated the annotation time by 5 to 10-fold; when needed, we corrected the annotation locally with VAST (see example in Fig. S2). We generated manually, also with VAST, the ground truth annotations for nuclear pores (Fig. 6A) clathrin and coated structures (Fig. 7A). In all cases, we applied the data augmentation procedure to the curated manual annotations.

### Deep learning segmentation pipeline

Our general training strategy (schematically represented in Fig. 1A) relied on a 3D convolutional neural network (CNN) architecture based on a 3D U-Net (Çiçek et al., 2016) (Fig. S3A); this approach has been used previously for segmenting intracellular structures in electron microscopy data (Guay et al., 2021; Heinrich et al., 2021; Wei et al., 2020). For each organelle class, we used a single, dedicated deep neural network, trained on augmented ground truth annotations generated from a small number of annotations contained within subvolumes (∼ 2-80 µm^3^) of the FIB-SEM data (Table S4). We used binary cross entropy as a loss function and trained each model for roughly 100k iterations on a single GPU (∼23h), after which the training/validation loss converged to a stable value (Fig. S3B-D). To avoid overfitting, we validated the evolution of the model periodically during the training session, by monitoring the loss between the model prediction and the subset of ground truth annotations in validation blocks of the FIB-SEM image not used for training. The final model yielded a predicted map that assigned to each voxel a probability of belonging to the structure (Fig. 1B), from which we derived a final binary map by setting a threshold value of 0.5. These models, unique for a given organelle or structure, were then used to find the specific cellular structure of interest in the FIB-SEM images of regions excluded from training or of ‘naïve’ cells that had not been used for training at all.

As previously noted by others, we also observed that the image contrast and texture of FIB-SEM data can vary substantially between different acquisitions, depending not only on cell type and mode of sample preparation (CF, HPFS), but unexpectedly also between adjacent cells of the same type in the same Epon block (Fig. S4). We found empirically that while the neural network could be trained to segment organelles successfully from samples prepared by the same mode of preparation, a model trained with ground truth annotations from HPFS cells failed when applied to CF treated cells, and vice versa. Although routinely implemented in our pipeline, contrast normalization by contrast-limited adaptive histogram equalization (CLAHE) (Pizer et al., 1987; Zuiderveld, 1994) of FIB-SEM data sets from different cells failed to improve the predictions (Table S5). Novel use of local shape descriptors as an auxiliary learning task (Sheridan et al., 2021) calculated from the ground truth annotations and representing high-level morphological notions such as object size and distance to object boundary also did not improve model prediction. As described below in detail, we resolved this problem by combining ground truth annotations from both data sets for training.

### Automated segmentation of organelles

We first applied ASEM to perform automated segmentation of FIB-SEM images from cells prepared by CF of nominal 5 nm isotropic resolution and relatively high contrast (Fig. 2 and Video 2); the summary shown in Table S6 illustrates the predictive performance obtained for models specific for mitochondria, ER and Golgi apparatus. For mitochondria, we selected from Cell 1 a training block of about 462 x 10^6^ voxels (1200 x 700 x 550 voxels) and used semi-automated annotation to generate ground truth annotations for the mitochondria contained within this volume, representing roughly 8% of all voxels (Table S6); Model performance was assessed every 1,000 iterations during the training phase by calculating the cross-entropy loss between the current prediction and the mitochondria ground truth within a validation block (not used during training). Additional smaller validation blocks (Table S4) containing mitochondria ground truth from naïve Cells 2, 3 and 6 were used to avoid overfitting during the training phase and to validate the model performance by measuring the validation losses. Validation losses rapidly converged within 20,000 - 40,000 training iterations, resulting in a relatively high F1 score (0.91) for Cell 1 and lower values for the data from naïve Cells 2, 3 and 6 (0.47, 0.66, 0.81, cf. Table S6). Similar results were obtained when training with ground truth annotations from Cell 2 instead of Cell 1 (Table S6); the validation losses also converged within 20,000-40,000 training iterations with good F1 scores for Cell 2 (0.87) and naïve Cells 1 and 3 (0.89, 0.74) and a slightly lower score for Cell 6 (0.7) with no further improvement with additional training iterations.

To find additional ways to enhance the generalization ability of the model, we modified the training pipeline to combine the ground truth annotations from Cells 1 and 2. We first tested the performance of the mitochondria model using the validation blocks in naïve Cells 3 and 6. In this case, the new model had a significantly improved performance (Table S6), reflected by even higher F1 scores for naïve Cells 3 and 6 (0.75, 0.88), but only after 95,000 - 115,000 iterations (Fig. S5). A similar improvement in model performance was observed for ER predictions when we first combined ground truth annotations of Cells 1 and 2. The new ER model, then used to predict ER in Cells 1, 2 and naïve Cells 3 and 6, led to generally improved F1 scores of 0.95, 0.90, 0.92 and 0.77, respectively (Table S6). Consistent with F1 scores smaller than the optimal value of 1, we observed by visual inspection a small number of false negative (yellow arrows) or false positive (red arrows) assignments as highlighted in Fig. 2A (see also Movie 2). Combining ground truth annotations from Cells 19 and 20 during training to predict the more complex Golgi apparatus in naïve Cells 3 or 6 marginally outperformed the models trained with either Cell 1 or Cell 2 (Table S6), also illustrated with one example of visual inspection of ground truth annotations and predictions showing instances of false positive assignments (red arrows, Fig. 2B). Thus, the predictive performance of a model could often be improved by using a model obtained by jointly training with ground truth annotations from two cells instead of training with data from one cell or the other.

We also tested the performance of ASEM using FIB-SEM images and ground truth annotations acquired by the OpenOrganelle initiative (Xu et al., 2021) (Table S1). These cells were prepared by HPFS and imaged with higher isotropic resolution (4 x 4 x 3-5 nm) but lower contrast). We examined the ability of our training pipeline to segment these data sets and focused on mitochondria and ER but not Golgi due to a lack of a sufficient number of ground truth annotations for Golgi objects in the available OpenOrganelle datasets (Table S7). We generated independent models for mitochondria and ER, by training with corresponding combined ground truth annotations from Hela Cells 19 and 20, followed by model performance verification using unseen ground truth annotations from the same Hela cells or from different types of naïve cells not used for training (Cell 21 Jurkat-1 and Cell 22 Macrophage-2, Table S7). Our pipeline performed well after ∼ 100K training cycles and managed to segment mitochondria in unseen data from each of the two Hela cells used for training (F1 scores of 0.99, Table S7) and from unseen data from each of the naïve Cell 21 Jurkat-1 or Cell 22 Macrophage-2 (F1 scores of 0.94 and 0.93; Table S7). Automated segmentation of the ER was less efficient, requiring ∼ 200K training cycles to reach the highest model performance (F1 scores of 0.91, 0.80, 0.48 and 0.81, respectively; Table S7). These first results indicate that our training strategy can create predictive models for successful identification of mitochondria, ER and Golgi apparatus in cells prepared by CF and of mitochondria and Golgi in samples prepared by HPFS.

To test the strength of combining ground truth annotations from two cells to train and then predict on a naïve cell, we explored the tolerance of the training pipeline to modest variations in image resolution. The results are shown for the representative FIB-SEM images in Figs. 3 and 4A and Video 3 obtained for a naïve Cell 15 SVG-A prepared by HPFS acquired at an isotropic 5 x 5 x 5 nm (Table S4); visual inspection of the images show successful predicted segmentations for mitochondria, ER and Golgi apparatus using models obtained by combined training with ground truth annotations from Hela cells 19 and 20 also prepared by HPFS and whose FIB-SEM images were acquired with mixed resolutions of 4 x 4 x 5.2 and 4 x 4 x 3.2 nm, respectively (Table S8).

**Figure 3.**
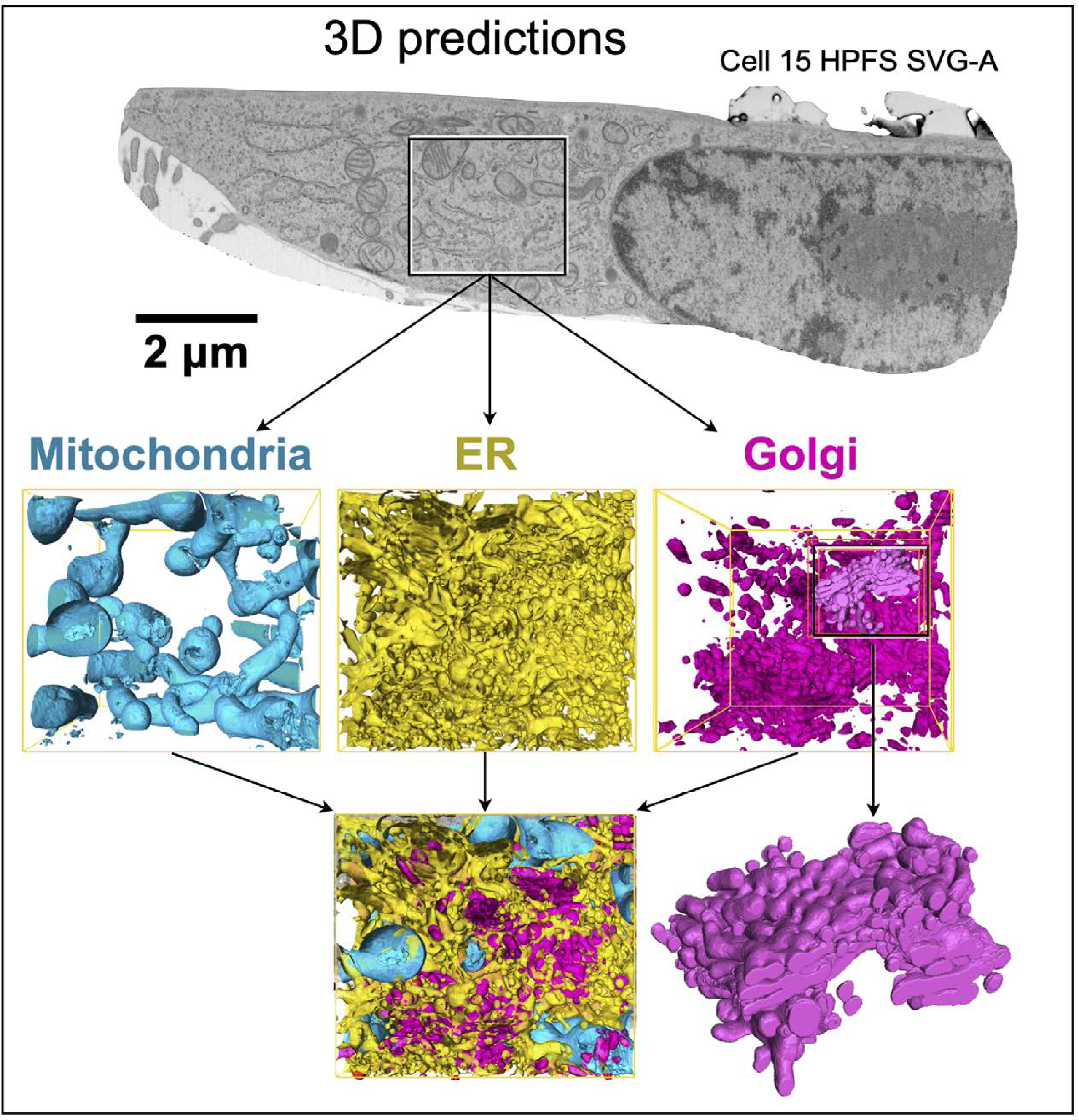
Network predictions of mitochondria, ER and Golgi apparatus. Single plane view from a FIB-SEM volume image from naïve cell 15 (SVG-A) not used for training prepared by HPFS and visualized during interphase at 5 nm isotropic resolution. The small region contains representative model predictions for mitochondria (cyan), ER (yellow) and Golgi apparatus (magenta) obtained from three 3D U-net models, each trained with organelle-specific ground truth annotations, without fine-tuning, from interphase cells 19 (Hela-2) and 20 (Hela-3) prepared by HPFS. Scale bar, 2 um.

**Figure 4.**
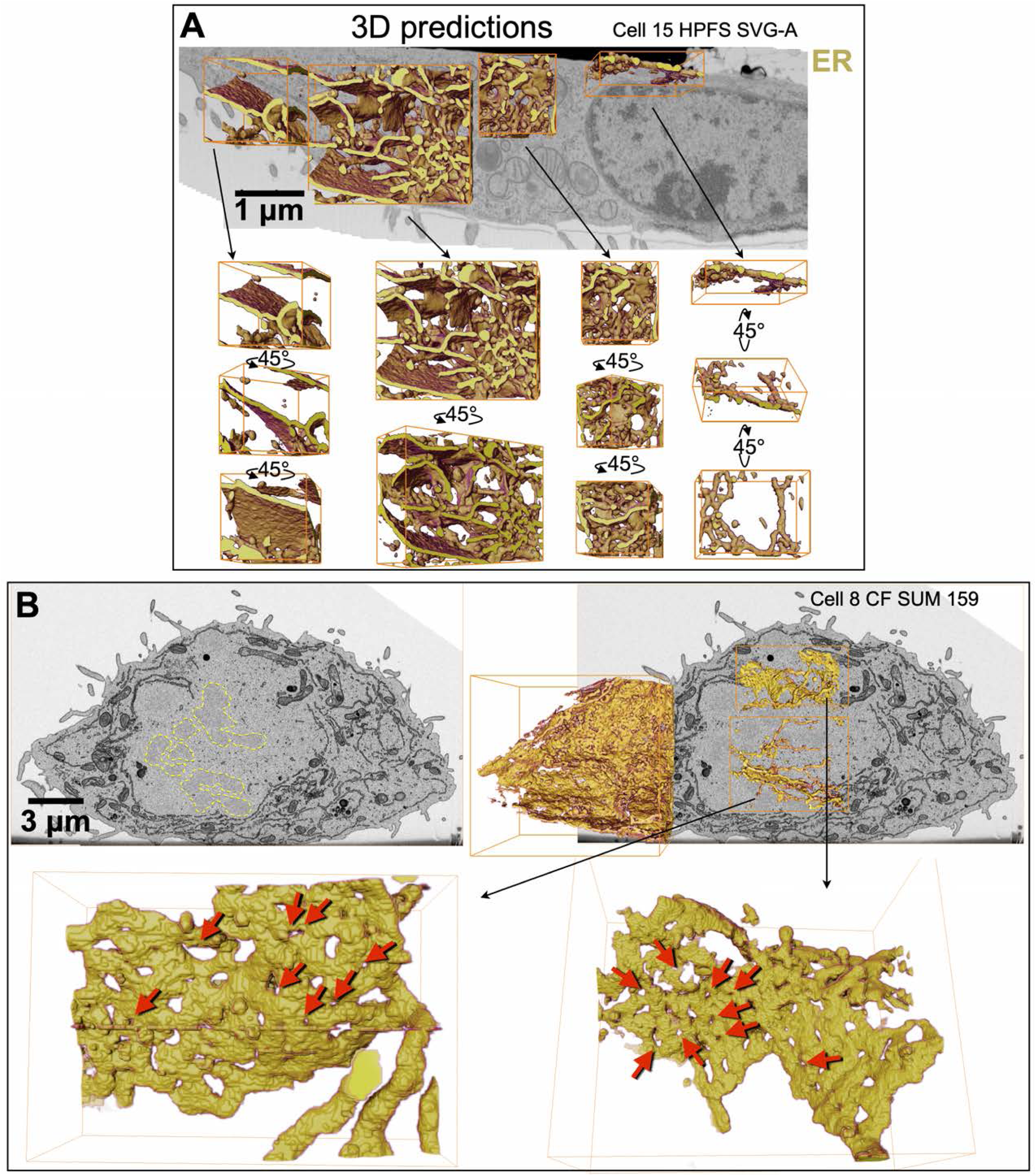
Predictive ER model resolves the structural complexity of the ER network during different stages of the cell cycle. **(A)** Representative examples of ER predictions in naïve cell 15 (SVG-A) processed during interphase as described in Fig. 3 showing the characteristic network of ER sheets connected at branch points to ER tubules. ER tubules were more abundant towards the periphery of this cell, ER sheets were more abundant closer to the nucleus. For clarity, manual VAST editing was used to eliminate pixels due to false positive predictions associated with the nuclear envelope. Scale bar, 1 um. **(B)** Representative examples of ER predictions from a mitotic naïve cell 8 (SUM 159) prepared by CF and imaged isotropically at 10 nm; the ER model was trained with ER ground truth annotations from interphase cells 1 and 2 (HEK293A) prepared by CF visualized isotropically at 5 nm resolution and downsampled to 10 nm. It shows successful recognition of an extensive network of fenestrated ER sheets (red arrow heads) connected to ER tubules, characteristic of mitotic cells. Ground truth annotations used to train the interphase ER model did not contain ER fenestrations, as they are barely present during stage of the cell cycle. Darker regions corresponding to chromosomes are outlined with yellow dotted lines. Scale bar, 3 um (see Video 3).

Because models generated with mitochondria or ER ground truth annotations from cells prepared by CF were unable to predict well on cells prepared by HPFS and vice versa, we explored the possibility of combining training data from both sample preparation protocols, using the same training datasets from HEK293A Cells 1 and 2, prepared by CF, and Cells 19 and 20 Hela prepared by HPFS. The mitochondria and the ER models performed nearly as well as specialized single-organelle models on almost all validation data sets regardless of sample preparation protocol used (Fig. 5 A and Table S7).

**Figure 5.**
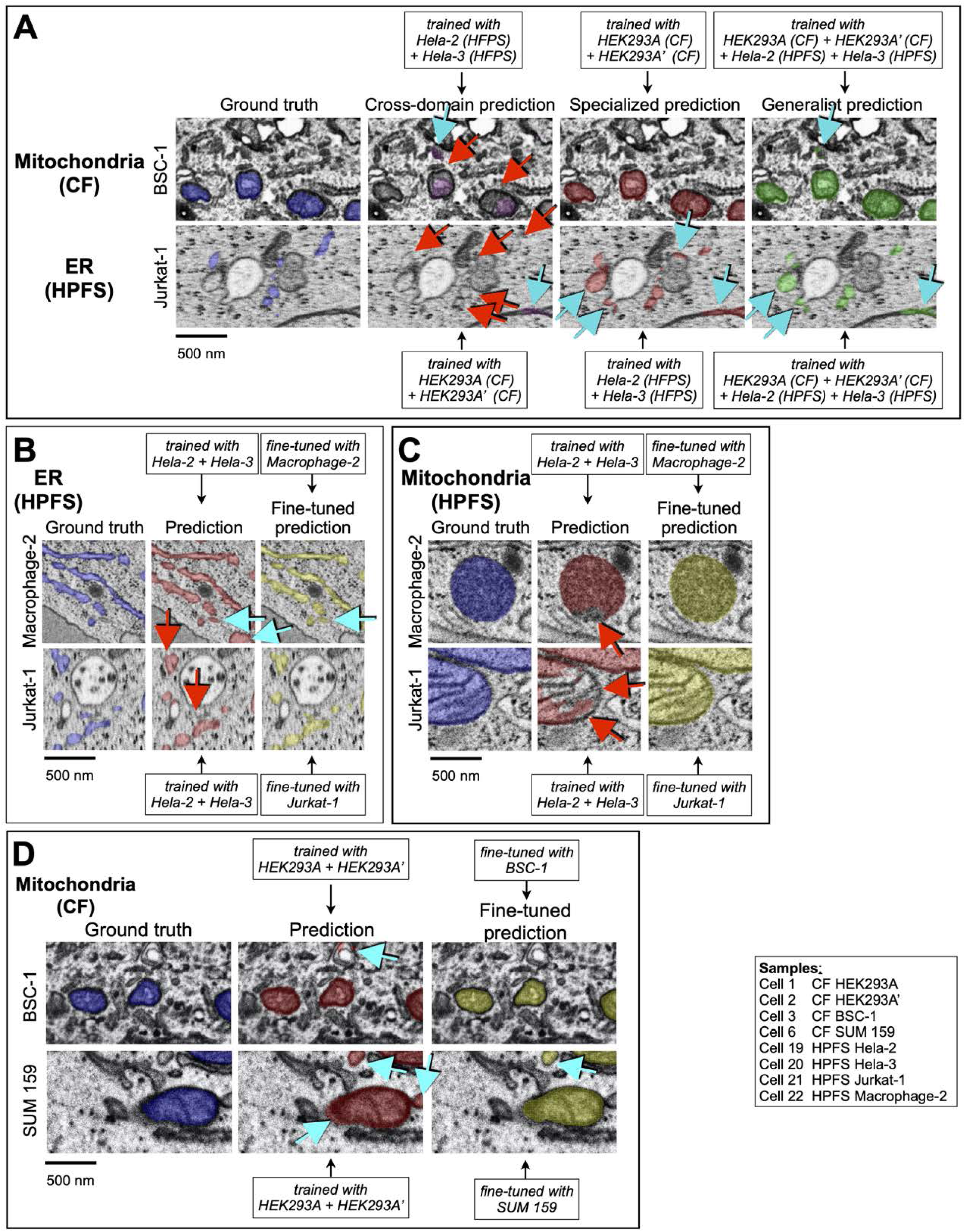
Effects of extensive combination of data sets and fine-tuning during training. **(A-D)** Examples to highlight the effect on the predictive performance of **(A, C, D)** mitochondria and **(A, B)** ER and models trained with data from cells prepared by CF or HPFS, with substantial differences in general appearance and contrast. The images show several comparisons between ground truth annotations and predictions from models trained as described in the insets with data obtained from cells prepared by different sample preparation protocols. Details of the cell and training protocols are in Tables S1, S2 and S8. Voxels corresponding to false positive (cyan arrows) and false negative (red arrows) predictions are indicated. Scale bar, 500 nm. **(A)** Predictions from *cross-domain* models, for which the training data and predictions were done using cells prepared with different sample preparation protocols, were less accurate than those obtained from the *specialized* models, for which training, and predictions were done using cells prepared with the same sample preparation protocol. Predictions from the *generalist* models, obtained by training using ground truth annotations from cells prepared by CF and HPFS, performed as well as the predictions from the specialized models. **(B-D)** Effect on the predictive performance of the models by fine tuning during training.

We also evaluated the performance of ASEM to predict mitochondria, ER and Golgi apparatus imaged with FIB-SEM data at 5 nm isotropic resolution but processed at the lower resolution of 10 nm. This test was done by using data sets from Cells 1 and 2 isotropically downsampled to 10 nm to train new models for mitochondria, ER and Golgi apparatus, then used to predict in validation data from Cells 1, 2, 3 and 6 isotropically downsampled to 10 nm (Table S9). These results showed that while the mitochondria and ER models performed similarly at both resolutions, the performance for the Golgi apparatus model notably decreased (Table S9), presumably explained by the relatively larger spatial complexity of the Golgi apparatus.

### Fine-tuning

To improve the predictive performance with images from naïve cells, we explored the effect of fine-tuning a pre-existing model on its segmenting accuracy, a simple implementation of transfer learning (Weiss et al., 2016). As described in Methods, we started with an already trained model and resumed model training for a low number of iterations (15,000) using only the new ground truth annotations from the naïve cell; the new ground truth annotations, although resembling those used for the first training, would typically have slightly different characteristics.

The following examples illustrate the range of results obtained upon implementation of fine-tuning using HPFS FIB-SEM data. The ER model, first obtained after ∼ 180,000 training cycles using ground truth annotations from Hela Cells 19 and 20, was then fine-tuned for additional 12,000 or 6,000 training cycles with small amounts of ground truth data from either naïve Cell 21 Jurkat-1 or Cell 22 Macrophage-2; both fine-tuning cases showed a significant improvement in the F1 precision scores, from 0.48 to 0.69 and from 0.81 to 0.90, without affecting recall (Fig. S5, Table S10); in other words, the model learned to correctly classify ER while at the same time reducing the number of false positives by rejecting structures that appeared similar but did not belong to the same semantic class (Fig. 5B). The next two cases of fine tuning illustrate little or no improvement in predictive model performance for mitochondria in cells prepared by HPFS or CF (Fig. 5C, S5 and Table S10). The model obtained after 95,000 training cycles using HPFS FIB-SEM data from Hela Cells 19 and 20 showed similar F1 scores (0.93) for naïve Cell 21 Jurkat-1 or Cell 22 Macrophage-2 before or after fine-tuning for 7,000 cycles. Similarly, a mitochondria model obtained after 95,000 training iterations using CF FIB-SEM data from Cells 1 and 2 and then fine-tuned for additional 6,000 fine-tuning training steps using ground truth annotations from Cells 3 or 6 showed either a significant increase (from 0.75 to 0.88) or no increase at all (0.88) in F1 scores, respectively (Fig 5D and Table S10). The fine-tuning strategy could not adjust mitochondria, ER or Golgi apparatus models generated using cells prepared by HPFS to predict in naïve CF treated cells, and vice versa. Likewise, fine-tuning had minimal or no effect for situations in which the pretrained model produced a prediction of naïve cells with a high F1 score, such as with mitochondria with an F1 score of around 0.9. We conclude that fine-tuning can be beneficial for segmenting relatively large membrane-bound organelles particularly in cases where the pre-trained model behaved poorly in naïve cells, but it could not resolve situations in which the staining characteristics of the samples were extremely different, even though they had been prepared by the same staining procedures.

### Automated segmentation of nuclear pores

To test whether our pipeline can automatically identify and segment small intracellular structures, we trained the neural network with ground truth annotations from nuclear pores, structures embedded in the double-membrane nuclear envelope with membrane pore openings of ∼100-120 nm in diameter. We used FIB-SEM data with nominal 5 nm isotropic resolution from interphase SVG-A and Hela cells imaged using HPFS to ensure minimal perturbations in the structural organization of the nuclear pores and their surrounding inner and outer nuclear membranes.

We used VAST to generate ground truth annotations for ten nuclear pores from Cell 13a – SVG-A (5 x 5 x 5 nm isotropic resolution) (Table S4). The segmentations, representing the inner and outer nuclear membrane envelope contours immediately adjacent to nuclear pores, also included 5 additional pixels (∼ 25 nm) of inner and outer nuclear membrane extending away from the nuclear pore opening (Fig. 6A). Training was performed with the augmented data generated from only eight nuclear pores (with two additional objects for validation), resulting in a nuclear pore model that performed well after 100,000 training cycles (F1=0.52, Precision=0.35, Recall=0.99, Table S8). In all cases, the high recall score was consistent with a perfect correspondence to all the voxels that defined the ground truth annotations. The relatively low F1 and precision scores reflected ‘fatter’ predictions due to voxels assigned to positions immediately adjacent to the ‘single row’ of voxels overlapping the nuclear pores in the ground truth annotations. Visual inspection confirmed accurate identification of all nuclear pores in naïve SVG-A cells 15 (Video 4) and 17 (5 x 5 x 5 nm isotropic resolution) and Cell 19 Hela (4 x 4 x 5.2 nm) not used for training (Fig. 6B). Because of the high predictive accuracy attained with this simple nuclear pore model (Video 4), it was not necessary to improve the model using our more extended training pipelines.

**Figure 6.**
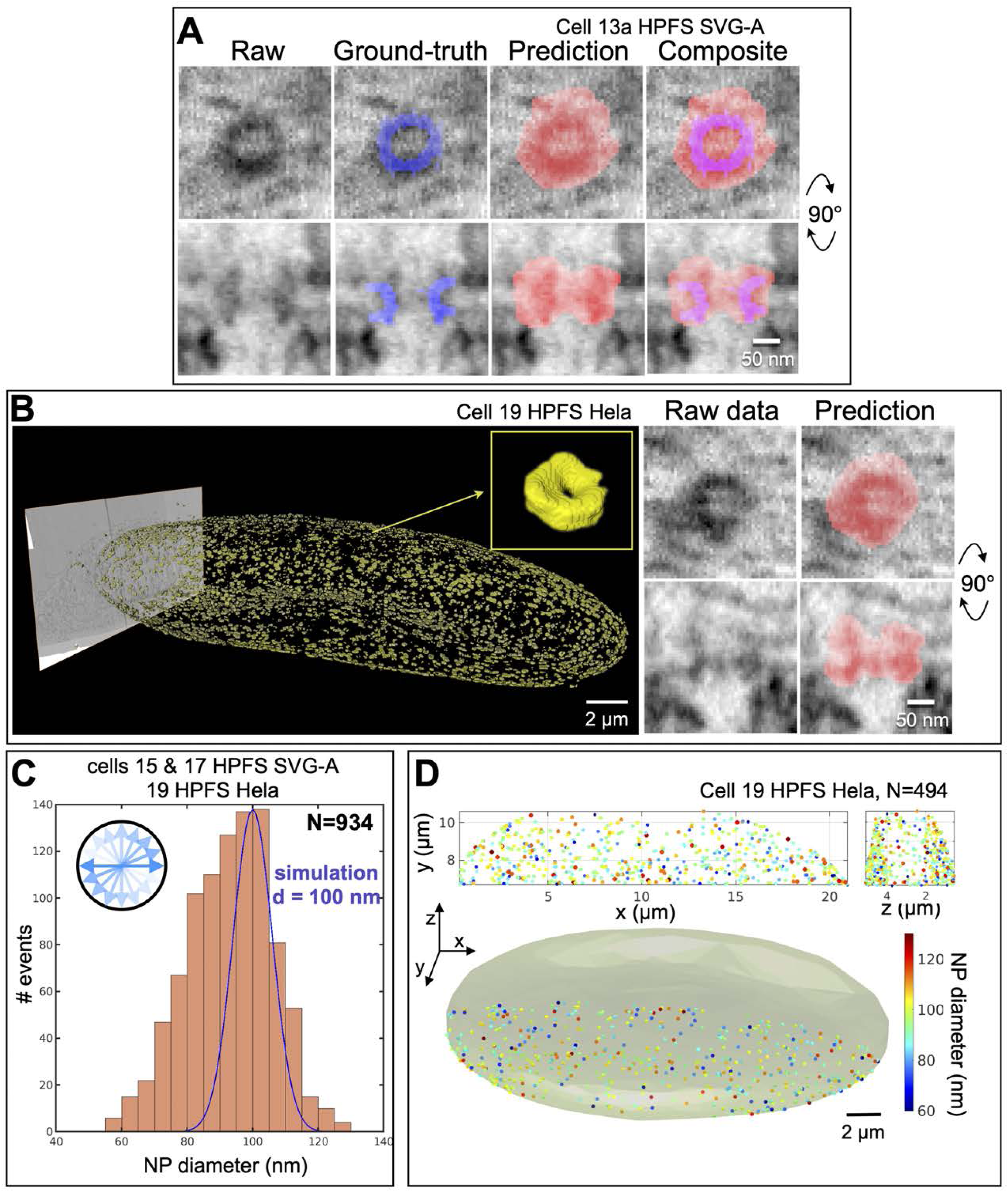
Identification of nuclear pores and variations in their membrane pore diameter. A nuclear pore model was generated by training without fine tuning used ground truth annotations of nuclear pores from cell 13a SVG-A prepared by HPFS and imaged at 5 nm isotropic resolution. **(A)** Orthogonal views of a representative nuclear pore not used for training show ground truth annotations and model prediction. Scale bar, 50 nm. **(A) (B)** Nuclear pore predictions for all the pores on the nuclear envelope of naïve cell 19 (Hela-2) prepared by HPFS and visualized during interphase at 4 x 4 x 5.3 nm isotropic resolution (left panel); the inset highlights the characteristic doughnut shape of the nuclear pore. Scale bar, 2 um. Representative orthogonal views (right panels) of a nuclear pore and model prediction. Scale bar, 50 nm. **(B)** Histogram of nuclear membrane pore diameters of nuclear pores measured in naïve cells 15, 17 and 19 (N=934) identified by the nuclear pore model. Each membrane pore diameter determined in the raw image represents the average value from 18 measurements spaced 10° apart (see inset and Methods). The Gaussian fit (blue) shows the expected size distribution if the data had come from membrane pores of a single diameter centered on the experimentally determined median (most abundant species); the bar width (6 nm) corresponds to the expected error of the measurements (see Methods). **(C)** Three-dimensional distribution of nuclear pores on the nuclear envelope of cell 19, color coded as a function of membrane pore diameter.

Based on ensemble cryo EM data from thousands of nuclear pores that provide a unique atomic model per data set (Schuller et al., 2021), combined with more selective images of single nuclear pores obtained using cryo tomography of yeast cells in different physiological states (Zimmerli et al., 2021), it is now believed that the diameter of the nuclear pore varies in response to the physiological state of the cell. It is not known, however, to what extent this size variability occurs within a single cell in a unique physiological state. Taking advantage of our automated segmentation pipeline that makes it practical to analyze hundreds of single nuclear pores, we explored the extent by which their membrane pore diameters varied within a single cell. Inspection of the nuclear membrane surrounding the pores viewed along the axis normal to the nuclear envelope confirmed the radial symmetry of the pore (Fig. 6B) with a relatively broad and continuous variation in membrane pore diameter, ranging from 60 to 130 nm (median 92 nm, with 75-108 nm 10-90 percentile range: n= 934; 305, 135, and 494 pores from SVG-A Cells 15 and 17, and Hela Cell 19, respectively) (Fig. 6C); these values were obtained by measuring in the raw images the distances between the peak signals at opposite ends of the nuclear membrane pore (see Methods and Fig. S6 A-D). The membrane pore sizes did not follow a normal distribution, but instead had a slight asymmetry contributed by smaller species. They were also distinct from the Gaussian fit (blue, Fig. 6C) corresponding to the expected size distribution if the data would have originated from a single pore size centered on the most abundant species (d = 100 nm). We found no evidence to suggest presence of spatial correlation between pore diameter and different regions of the nuclear envelope within the cell, for example away from the cover slip or normal to this surface, nor did we find evidence of local clustering of pores with a favored size (Fig. 6D and S6E).

### Automated segmentation of clathrin coated pits, coated vesicles and caveolae

As a further test of ASEM with relatively small structures, we chose clathrin coated pits, 30-100 nm membrane invaginations in the plasma membrane and the trans Golgi network (TGN) involved in selective cargo traffic (Kirchhausen, 2000). We trained the model with ground truth annotations from 15 endocytic plasma membrane coated pits of different sizes and shapes thus representing different stages of clathrin coat assembly. While the resolution of the FIB-SEM was insufficient to discern the familiar spikes or the hexagonal and pentagonal facets of a clathrin coat as seen in samples imaged by TEM, the presence of strong membrane staining, which we attribute to clathrin and associated proteins (Fig. 7A), made these invaginations recognizably distinct from caveolae, which are smaller (50 – 100 nm) flask-shaped invaginations and lack enhanced membrane staining (Fig. 7B). None of the cells had recognizable regions of strongly stained, flat membrane, often found on the coverslip-attached surface of cells in culture and in other specialized locations (Akisaka et al., 2008; Grove et al., 2014; Heuser, 1980; Maupin and Pollard, 1983; Saffarian et al., 2009; Signoret et al., 2005). We used VAST to generate the coated-pit ground truth annotations, which were simply a collection of single traces loosely overlapping the endocytic membrane invagination (Fig 7A, blue).

**Figure 7.**
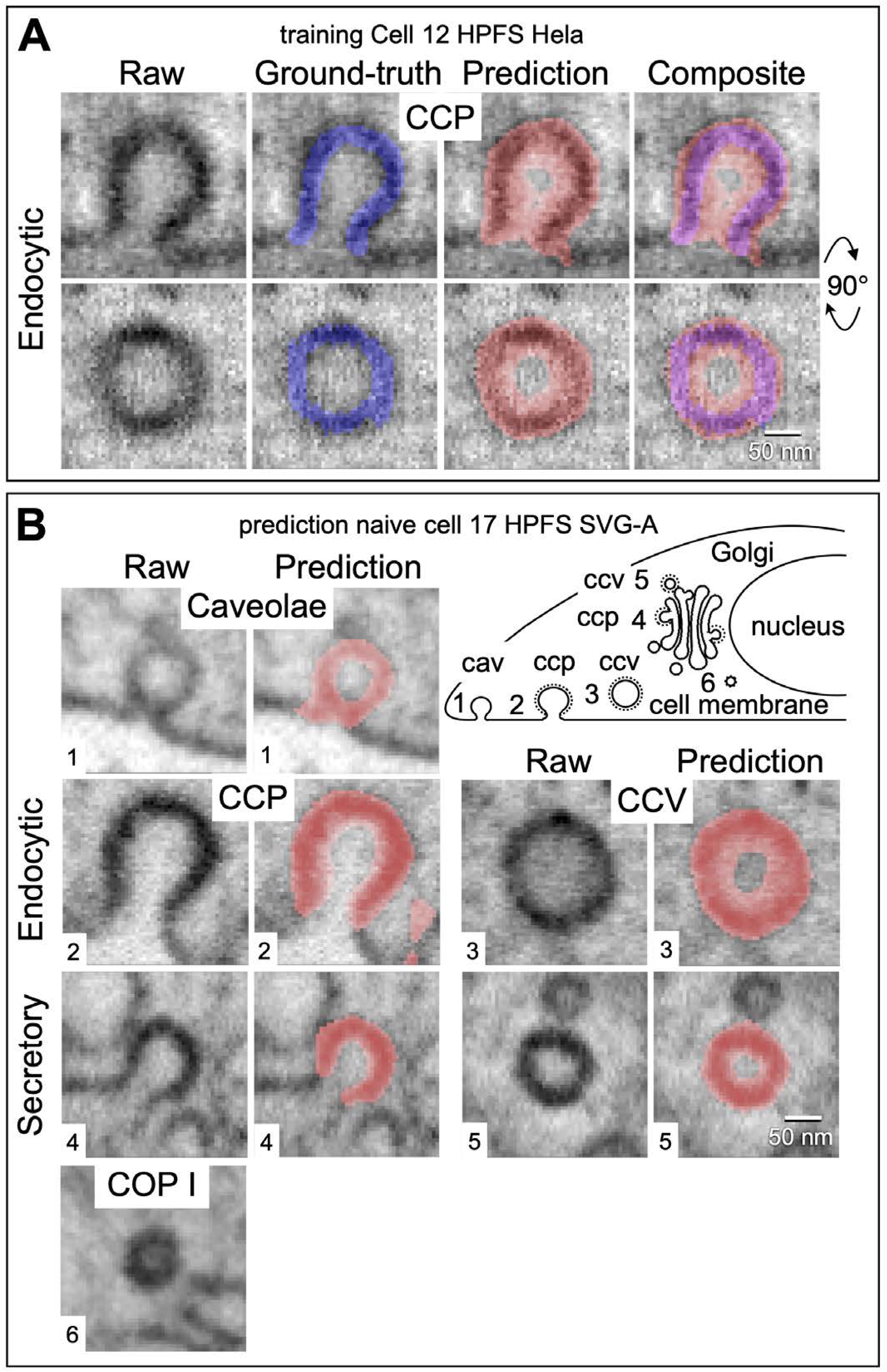
Identification of clathrin coated pits, coated vesicles and caveolae. A coated pit model was generated by training with ground truth annotations from Cell 12 (Hela-2) prepared by HPFS and imaged at ∼ 5 nm isotropic resolution. **(A)** Orthogonal views of a representative endocytic clathrin coated pit (CCP) not used for training showing ground truth annotations and model prediction. Scale bar, 50 nm. **(B)** Orthogonal views of a caveola, an endocytic clathrin coated pit (CCP) and a clathrin coated vesicle (CCV) at the plasma membrane, and a coated pit (CCP) and vesicle (CCV) associated with membranes from the secretory pathway. Each panel shows the ground truth annotation and the model prediction. An example of a COPI vesicle not predicted by the coated pit model is also shown. Views are from naïve cell 17 SVGA prepared by HPFS and imaged with ∼ 5 nm isotropic resolution.

The coated pit model obtained after 80,000-100,000 training iterations combined 15 ground truth annotations we made, six from Cell 12 Hela and nine from Cell 13 Hela imaged by the COSEM project. Visual inspection of the predictions generated by this relatively simple training in parts of Hela cells 12 and 13 cells that had not been used for training showed accurate recognition of all endocytic coated pits (representative example in Fig. 7B); we obtained similar results from naïve SVG-A cells 15 (Video 4) and 17, and Hela cell 19. The model also identified all coated pits in the TGN (Fig. 7B). It incorrectly identified caveolae as coated pits (Fig 7B), but we could detect no other incorrect predictions anywhere in the cell volume. Sharply invaginating curvature of the stained membrane outline thus appears to be an important component of the pattern the model learned to recognize.

We used our additional annotated ground truth annotations from Hela Cells 12 and 13 that had not been included in the training set to calculate F1, recall and precision scores (Table S8). In all cases, the high recall score (0.99) demonstrated the almost perfect reconstruction of all voxels belonging to the ground truth annotations. The relatively low F1 and precision scores (∼ 0.65 and 0.51) were due to incorrect voxel predictions immediately adjacent to the ‘single row’ of true voxel assignments overlapping the invaginated membrane in the ground truth annotations (Fig. 7A, B).

The model also recognized vesicles near the plasma membrane and the TGN that an expert human observer would have interpreted from their staining to be clathrin coated vesicles, even though training of the model did not include ground truth annotations representing them (Fig. 7B). We confirmed that the model recognized *all* the presumptive coated vesicles in a cell, by visual inspection across the full volumes of Hela cells 12 and 13, as well as of three cells that did not contribute at all to the training set, SVG-A cells 15 and 17 and Hela cell 19. Training on endocytic coated pits thus also allowed recognition of endocytic coated vesicles and TGN coated pits. In contrast, the model did not recognize vesicles associated with the Golgi apparatus or the ER an interpreted by their staining as COPI or COPII.

We took advantage of the large, combined set of three-dimensional image data from coated pits and vesicles to analyze assembly stages using the metrics depicted in Fig. S7). We determined the depths and widths at ½ depth for each of the membrane invaginations in SVG-A cell 17 (Fig. 8A). Caveolae, recognized by the absence of an enhanced membrane signal, were relatively small, with narrow distributions of depths and widths centered on 61 and 81 nm (Fig. 8A). Endocytic coated pits, identified by their enhanced membrane signals, were generally larger than caveolae and had wider distributions of depths and widths, which clustered into two groups. Coated pits with open necks (>40 nm) had shallow, ∼ 50 nm invaginations; those with narrower necks (∼ 10 to 40 nm) had deep, ∼ 100-130 nm invaginations (Fig. 8B, left and central panel, and 8C, right panel). Endocytic coated pits and vesicles were also larger than the corresponding secretory structures emanating from internal membranes associated with the TGN (Fig. 8B, left panel).

**Figure 8.**
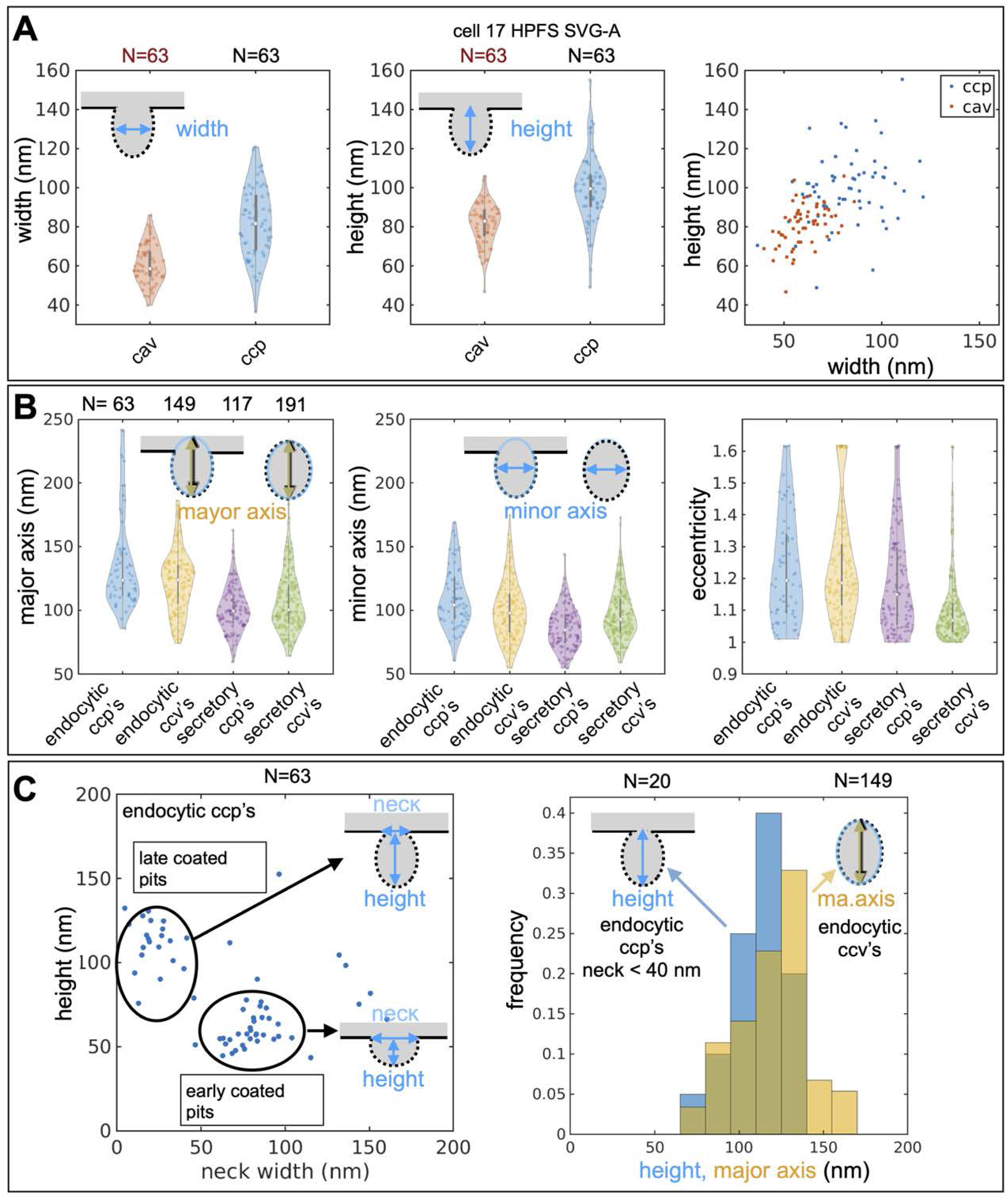
Dimensions of clathrin coated pits, coated vesicles and caveolae. **(A)** Violin plots of width and height for caveolae (CAV) and endocytic clathrin coated pits (CCP) in the raw images of the structures identified by the coated pit model in cell 17 (see also Figure S8, and Clathrin coated pits and vesicles, Methods). **(B)** Violin plots of major and minor axis and eccentricity of the fitted ellipse of all pits and vesicles in the raw images of the structures identified by the coated pit model in cell 17 (see also Figure S8, and Clathrin coated pits and vesicles, Methods). **(C)** The left-hand panel shows the distribution of height versus neck width for endocytic clathrin coated pits in cell 17, identified by the coated pit model. The plot shows two clusters, which correspond to early and late coated pits, respectively, as illustrated by the schematics (see also Figure S8, and “Clathrin coated pits and vesicles” in Methods). The right-hand panel shows histograms for height and major axis of the fitted ellipse for late endocytic coated pits and coated vesicles, respectively.

The eccentricity of the assembling pit, defined as the ratio of major and minor axes of the ellipsoid that fit best to a given membrane profile, showed a relatively narrow and overlapping distribution (Fig. 8B, right panel), ranging from 1 (symmetric) and 1.6 (less symmetric) for endocytic pits and vesicles. Most of the pits and vesicles associated with internal membranes in SVG-A Cell 17 (Fig. 8B, right panel) had eccentricities close to 1; in those cases, the major axis of most pits was orthogonal to the plane from which the pits invaginated. Similar results were obtained for SVG-A cell 15 and Hela cells 12 and 13 (Fig. S8, A-C). These results are consistent with a budding mechanism in which stepwise growth of the clathrin coat drives invagination of the membrane, ultimately creating a constriction, as the curved clathrin lattice approaches closure, that is narrow enough for dynamin to pinch off the nascent vesicle (Kirchhausen et al., 2014).

## Discussion

The automated 3-D image segmentation pipeline embodied in ASEM overcomes three critical hurdles for making FIB-SEM more practical and more broadly useful than currently available procedures. (1) Our graph-cut based annotation approach facilitates and simplifies the manual stages of ground annotation for convoluted structures like the Golgi apparatus, by minimizing the number of hand-curated annotations. Between 8 and 16 annotated ground truth annotations encompassing the complete volumes of smaller objects (nuclear pores, clathrin coated pits) or partial volumes of larger ones (mitochondria, ER, Golgi apparatus) were generally enough when augmented as described. We used rough annotations that could be off by 1-2 voxels rather than voxel-precise labeling to delineate the outline of the intracellular structures for which we were training. While this strategy was effective for our training pipeline, it was much less time-consuming than the precise delineation efforts used by COSEM (See Methods). We could then readily correct any erroneous voxel detections, either by manual intervention or by automatic post-processing. (2) For the applications described here, ASEM requires far less computational effort than does COSEM or other approaches, largely because we restrict the training to a single type of structure and thus create a separate model for each type. Consequently, we found that about ∼100,000-150,000 training iterations were sufficient for accurate prediction, whereas COSEM required five times as many. (3) We can substantially improve the success rate in a completely naïve cell by using a model trained on ground truth annotations from another cell and re-training by a simplified transfer-learning approach with a very small number of ground truth annotations from the new cell, thereby adapting the model to a cell with slightly different imaging characteristics at the cost of modest additional segmentation and computational effort. In the examples here, just 5,000-10,000 training iterations were enough to increase prediction accuracy throughout the rest of that cell.

To test the robustness and flexibility of ASEM, we used the model trained with ER ground truth annotations from cells in interphase for identifying and segmenting ER in an early anaphase mitotic cell. The model, which had correctly identified and segmented the complete ER in a naïve interphase cell imaged at ∼ 5 nm isotropic resolution, also accurately identified and segmented the ER in the mitotic cell imaged at 10 nm isotropic resolution with a model trained with ground truths from the same cells in interphase but computationally downsampled to 10 nm (Fig. 4B and Video 4). The result is non-trivial, because relatively extended, fenestrated, double-membrane sheets, with small interconnecting tubules, dominate the morphology of the mitotic ER, while tubules of varying lengths, connecting much smaller sheets, are the principal structures in the interphase ER. Segmenting the mitotic ER within the imaged cell volume (∼2^9^ voxels, voxel size 10 x 10 x 10 nm) required less than an hour; it would have taken a human annotator several months. Previous analyses were limited to small cell volumes precisely because of this constraint. We further showed that automatic segmentation of the Golgi apparatus with ASEM confirmed the results described by the COSEM Project team (Heinrich et al., 2021). The Golgi is not a stack of closely packed, uniform cisternae, as often diagrammed in textbooks. Rather, each member of the stack is a complex, perforated structure of variable shape, surrounded by many small vesicles.

We used automatic segmentation of mitochondria, ER and Golgi apparatus primarily for comparison with published results from other methods, to validate the features of ASEM designed to accelerate and simplify the entire pipeline. We turned to smaller intracellular structures as tests of new and potentially more challenging applications. Nuclear pores are more homogenous than larger organelles, and thus in principle easier to recognize, and while any one pore has much less distinct substructure than does a Golgi stack or a mitochondrion, we have found, the diameter of the membrane pore varies even across the nucleus of single cells at a fixed time point, despite the likely invariance of much of the nuclear pore complex protein assembly. Clathrin coated pits are both small (on the scale of the ER and Golgi apparatus) and variable, in size and as well as in assembly stage. In both cases, by training ASEM with a large set of ground truth annotations generated by data augmentation from a very small number of hand-annotated objects, we could automatically identify essentially all the objects in the cell, despite the variable diameter of the nuclear membrane pore and the variable size and stage of completion of a clathrin coated pit. Moreover, a model trained on plasma-membrane coated pits identified coated pits in the Golgi and free clathrin coated vesicles in the cytosol.

The osmium-uranyl staining in current FIB-SEM sample preparation, for both CF and HPFS, preferentially marks lipid headgroups, proteins and nucleic acids. Although with the training set used here, the model did not distinguish between clathrin coated pits and caveolae, the eye clearly picks up the much heavier staining of the former (Fig. 7). The model correctly retrieved clathrin coated vesicles, as well as coated pits in the TGN, and distinguished them from COPI and COPII vesicles which carry cargo between the Golgi apparatus and the ER, perhaps because they are smaller structures, of substantially sharper curvature than the clathrin coated structures the model had learned to recognize. How well the model will find protein-dominated structures – e.g., virus assembly intermediates – remains to be determined.

Imaging the entire volume of a single cell at ∼5 nm resolution can answer questions that are much harder to tackle by methods such as cryo-tomography that access at somewhat higher resolution only a small slice of a cell. One example is our finding, that nuclear pores vary in size across the nuclear membrane and hence that the variability identified by cryo-tomography is present at an arbitrary time point in a single cell. The deep-learning protocols we have developed, and the readily implemented and freely accessible analysis tools we provide, form an experimental pipeline that will run entirely on commercially available workstations. We suggest that EM volume imaging will prove to be a powerful complement to fluorescence volume imaging afforded by lattice light-sheet microscopy (Chen et al., 2014; Gao et al., 2019; Liu et al., 2018).

## Material and methods

### Chemical Fixation, dehydration and embedding

Cells plated on glass coverslips were processed for chemical fixation (CF) by incubation for 30 minutes at room temperature with 0.2% glutaraldehyde (Electron Microscopy Science, Cat.16220) and 2.5% paraformaldehyde (PFA, Electron Microscopy Science, Cat. 15700) dissolved in 0.1M PIPES buffer (pH 7.4, Sigma-Aldrich, Cat. P6757), followed by a rinse with 0.1M PIPES buffer. A 2% OsO4 aqueous solution (Electron Microscopy Sciences) dissolved in 0.1M PIPES, pH 7.4 was used to stain the cells for 1 hour at RT, followed by incubation for another 1 hour at RT in a solution containing 2.5% potassium ferrocyanide (Sigma-Aldrich) in 0.1 M PIPES, pH 7.4. The cells were rinsed three times at five-minute intervals with deionized ultrapure water followed by a 30-minute incubation at RT with a filtered (Whatman, 0.2 μm) freshly prepared solution of 1% thiocarbohydrazide (Electron Microscopy Sciences) made by dissolving it at 60°C for 15 minutes. The cells were again rinsed three times at 5-minute intervals, followed by another incubation with 2% OsO4 aqueous solution for 1 hour at RT. The cells were again rinsed three times at five-minute intervals with ultrapure water, followed by two rinses with 0.05 M maleate buffer, pH 5.15 (Sigma-Aldrich), and finally incubated with 1% uranyl acetate (Electron Microscopy Sciences) dissolved in 0.05 M maleate buffer pH 5.15 for 12 hours at 4°C.

A resin mixture containing methylhexahydrophthalic anhydride (J&K Scientific) and cycloaliphatic epoxide (ERL 4221, Electron Microscopy Sciences) at a weight ratio of 1.27:1, mixed with the catalyzing agent (Hishicolin PX-4ET, Nippon Chemical Industrial) at a 1:100 ratio by volume, was prepared in a water bath sonicator at RT for 15 minutes.

During the same period, the glass coverslips with the attached CF samples were placed face up on wet ice and then rinsed twice for five minutes with ultrapure water, followed by dehydration using a graded series of ethanol solutions (30, 50, 70, 90%) each step lasting three minutes, then three washes in 100% absolute ethanol for 10 minutes ending with three washes with anhydrous acetone (Sigma-Aldrich) for 10 minutes at RT.

The glass cover slips with the attached, dehydrated CF samples immersed in anhydrous acetone were placed in a wide-mouth glass jar, mixed with the resin at a 1:1 volumetric ratio, and gently rocked on a plate rocker for 12 hours at room temperature. The resin mixture was then removed by aspiration and replaced with 10 ml of freshly prepared resin mixture and further incubated with gentle rocking for another two hours; this step was repeated thrice, each time with freshly prepared resin. Finally, the glass cover slips with the attached cells were placed on top of cut off caps from 1.5 mL Eppendorf tubes containing freshly prepared resin was oriented with the cells towards the cap, and the resin allowed to polymerize for 12 hours at 100°C. Upon resin hardening, the caps were immersed in boiling water for 5 minutes and then quickly transferred into liquid nitrogen leading to separation of the glass cover slip from the resin and retention of the cells in the polymerized resin.

### High pressure freezing, freeze-substitution and embedding

Cells were plated on 6 x 0.1 mm sapphire disks in MEM (Corning™ 10009CV) supplemented with 10% Fetal Bovine Serum (Atlanta Biologicals S11150). Two sapphire discs (Technotrade international 616-100), one or both containing attached cells facing inwards, were separated by a 100 μm stainless steel spacer (Technotrade international 1257-100) and processed for high pressure freezing on a Leica EM ICE high pressure freezer (Leica microsystems). Following high pressure freezing, the sapphire discs were placed under liquid nitrogen and transferred into the top of cryotubes placed in liquid nitrogen and containing frozen 2% OsO4, 0.1% uranyl acetate and 3% water in acetone; freeze substitution (FS) was carried using an EM AFS2 automatic freeze substitution device (Leica Microsystems) according to a pre-programmed FS schedule (-140°C to -90°C for 2h, -90°C to -90°C for 24h, -90°C to 0°C for 12h and 0°C to 22°C for 1h). Samples were then removed from the AFS2 device, rinsed 3 times in anhydrous acetone, 3 times in propylene oxide (Electron Microscopy Sciences), 3 times in 50% resin (24g Embed 812, 9g DDSA, 15g NMA, 1.2g BDMA; Electron Microscopy Sciences 14121) dissolved in propylene oxide, and finally transferred into embedding molds (EMS 70900) containing 100% resin; the resin was then allowed to polymerize for 48 hours at 65°C. The sapphire disc was then separated from the resin block by sequential immersion in liquid nitrogen and boiling water.

### FIB-SEM imaging

The polymerized resin blocks were cut from the molds and glued, with the free face facing away, onto the top of aluminum pin mount stubs (Ted Pella), using conductive silver epoxy adhesive (EPO-TEK H20S, Electron Microscopy Sciences). The free face was then coated with carbon (20 nm thickness) generated from a high purity carbon cord source (Electron Microscopy Sciences) using a Quorum Q150R ES sputter coater (Quorum Technologies) and the resin block loaded on the microscope specimen stage of a Zeiss crossbeam 540 microscope for FIB-SEM imaging. After eucentric correction, the stage was titled to 54° with a working distance of 5 mm for coincidence of ion and electron beams. A cell of interest was located on the free face of the resin block by SEM, after which a thin layer of platinum was deposited using the gas injection system. A coarse trench was then milled adjacent to the cell using the 30 kV/30 nA gallium ion beam. This block face was polished with a 30 kV/7 nA gallium beam before starting the interlaced sequence of FIB milling with a 30 kV/3 nA gallium beam and SEM imaging with a 1.5 kV/400 pA electron beam advanced in 5 nm steps. The X/Y pixel size was 5 nm to create isotropic voxels. For samples prepared by HPFS, we added registration marks on top of the platinum layer generated with a 1.5 kV/50 pA gallium beam, followed by contrast enhancement of the marks by irradiation with a 1.5 kV/5 nA electron beam and final deposition of second platinum layer. FIB-SEM images were collected using the Inlens detector with a pixel dwell time of 10-15 us. The FIB-SEM images were aligned post-acquisition with the Fiji plugin Register Virtual Stack Slices https://imagej.net/plugins/register-virtual-stack-slices, using the translation (Feature extraction model and Registration model) and shrinkage constraint options (Schroeder et al., 2021).

FIB-SEM data at 10 nm were acquired using a backscatter electron detector (EsB) with a grid voltage set to 808 V to filter out scattered secondary electrons, with a dwell time of 3 μs, line averaging of 8, and pixel size of 10 x 10 nm (X/Y). FIB milling was performed with the 30 kV/30 nA gallium ion beam in 10 nm steps to create isotropic 10 x 10 x 10 nm (XYZ) voxels. The sequential FIB-SEM images were registered using the Fiji plugin *StackReg* with Rigid Body transformation.

### Ground truth annotation

All our ground truth annotations were binary masks located at least 47 voxels away from the boundaries of the 3D FIB-SEM image. This ensured that training of the neural network was done with sufficient semantic context within the image resulting in improved model predictions.

Ground truth annotations for mitochondria, Golgi apparatus and endoplasmic reticulum were generated by using the carving module of Ilastik (Berg et al., 2019) or our graph cuts-based semi-automated annotation tool, and when needed, by further manual editing using VAST (Berger et al., 2018) to remove voxels that did not belong to the structure of interest or to add voxels for regions that had not been included in the original binary mask.

Ground truth annotations for the relatively complex three-dimensional substructure of the Golgi apparatus included membrane boundaries and the lumen for the characteristic 3-6 closely stacked fully enclosed membrane lamellae, the fenestrated and somewhat swollen trans-Golgi network stack, and the variable number of small vesicles clustered next to the Golgi apparatus. They were created with our semi-automated graph-cut annotation tool.

Ground truth annotations for endocytic clathrin coated pits and for caveolae were manually generated in consecutive planes using VAST, by drawing along the contours following the plasma membrane invaginations characteristic of these structures.

Ground truth annotations for nuclear pores were manually generated plane by plane using VAST, by drawing along the contours of the nuclear outer and inner membrane adjacent to the nuclear pore.

### Graph-cut annotation tool

We developed a new semi-automated tool to aid an expert annotator in marking *sparse* and *coarse* labels in a sub-volume, one plane at a time, so that the annotator can define high-level seeds required to generate ground truth annotations for a chosen organelle and separate it from the background (see Figure S2). Once the seeds are marked, the tool, at the push of a button, uses mathematical techniques (detailed below) to combine them with the grayscale values in the volume to infer the structure of the organelle, thus resulting in a semi-automated segmentation. The tool allows the annotator to visualize the volume plane-by-plane and define seed voxels in a plane as either part of the organelle or of background. Scrolling through, the annotator can seed a few planes at arbitrary intervals over the entire stack. It also allows views along all three axes so that the annotator can look down the *z*-axis to mark the *xy*-plane, and similarly for the *y*- and *x*-axes.

The technique of graph cuts-based segmentation was adopted to generate segmentations from these seeds. The seeds were defined on manually extracted volumes from cells. Even though the seeds were coarse, the annotator took care not to mis-label any voxel. The maxflow algorithm (Boykov et al., 2001) was then employed to segment organelles based on these annotations. This tool was written in Python and adopted to suit the annotation needs associated with 3D FIB-SEM data. It is publicly available at https://github.com/kirchhausenlab/gc_segment, accompanied by detailed usage instructions and best practices.

The next step of the semi-automated segmentation is a reduction of problem complexity. To get a segmentation based on the seeding, each voxel must be assigned either an organelle label, or a background label. At this point, we note that the volumes with which we work are characteristically large—a volume of 1 cubic micrometer contains 8 million voxels at a resolution of 5nm × 5nm × 5nm. We also observe that as organelles are contiguous objects in the volume, a group of nearby voxels have a high chance of belonging to the same organelle. We hence merge nearby voxels to form *supervoxels* using the SLIC algorithm (Achanta et al., 2012), so that we can work with ∼10^3^ supervoxels instead of ∼10^6^ voxels - a reduction of three orders of magnitude. Under this strategy, supervoxels were formed by grouping adjacent voxels together based on a similarity criterion, which for our problem setting was chosen as agreement in grayscale values.

A post-processing step has been included in this tool that can modify the resulting segmented foreground voxels by fitting to the foreground voxels a univariate Gaussian mixture model based on its grayscale values. This optional post-processing can help remove outlier voxels with minimal additional overhead of computation time.

The aim of the graph cuts-based strategy is therefore to define and optimize an objective energy function over the space of labels ℒ^V^ = {0,1}^V^, where the labels 0 and 1 represent organelle and background respectively, and *V* is the number of supervoxels in the volume. Each point in this space determines whether a supervoxel is considered organelle or background, and hence represents a segmentation of the volume. The objective energy function for our problem was formulated based on early work on graph-cut segmentation in computer vision (Boykov and Kolmogorov, 2004).

To describe our energy function, we introduce the following notation. The subscripts *u* and *v* denote supervoxels; the subscripts *p* and *q* denote voxels. Let G = (G_1_, G_2_,…, G_p_,…, G_S_), *G*_p_ ∈ {0,1}, *S* ≪ *N* be the labels for *S* seeded voxels for a volume consisting of *N* voxels, and let A = (*A*_1_, *A*_2_,…, *A*_u_,…, *A*_V_), *A*_u_ ∈ {0,1} be the vector corresponding to the unknown segmentation of the *V* supervoxels in the volume. The vector *A* represents the *deduced labelling* (or *segmentation*) for the volume. The final segmentation is the vector that results in the *least* value of the objective energy function. The energy function is

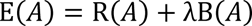

where

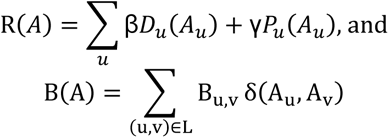

Here, δ(*A*_*u*_, *A*_*v*_) is set equal to 1 if *A*_u_ ≠ *A*_v_ and 0 otherwise. *N* denotes the set of supervoxels adjacent to each other and hence deemed *neighbors*. Adjacency implies a shared boundary between the two supervoxels. β, γ, and λ are weights given to the individual terms. *P*_u_ and *D*_u_represent *unary* terms of the energy function, as they depend only on one supervoxel, while *B*_u,v_represents *pair-wise* terms.

The unary terms are defined as follows. The terms *P*_u_ are determined by the aggregate grayscale values of supervoxels and their agreement with the seeded foreground and background voxels (the vector *G*), as in earlier work (Boykov and Kolmogorov, 2004; Boykov et al., 2001), and the terms *D*_u_ are determined by the distance of the voxels from the nearest seeding of organelle and background. The pair-wise terms are also defined according to earlier work, based on the distance between two supervoxels, defined as the distance between the arithmetic center of two supervoxels. As the defined energy function is sub-modular, it can be optimized using graph cuts (Kolmogorov and Zabin, 2004). An efficient algorithm to optimize such functions, called *maxflow* (Boykov and Kolmogorov, 2004), was used to find the optimum vector A. There are hyper-parameters in our formulation, namely β, γ, and λ. These were empirically defined as β = 1, γ = 1, and λ = 10. It should be noted that these values can be tuned according to the user’s needs and observations.

### Data preprocessing for deep learning

Cell image stacks underwent the following steps before they were ready for training:

1. Conversion from TIFF format to the block-wise storage format ZARR. The size of a FIB-SEM dataset corresponding to a stack of registered TIFF files (approximately 2000 planes) was about 20GB. These TIFF stacks were converted into a ZARR 3-D compressed array [https://zenodo.org/record/7115955] to increase the efficiency for further pre-processing steps and, most importantly, for the neural network training.
2. Cropping of the dataset to exclude empty regions outside the cell and to speed up all further preprocessing steps.
3. Block-wise adjustment of brightness and contrast with 3D contrast-limited adaptive histogram equalization (CLAHE, (Zuiderveld, 1994)) using *scikit-image.exposure.equalize_adapthist* with kernel size 128 and clip limit 0.02 (see figure S4).
4. Application of morphological operations to automatically clean up ground truth annotations based on biological assumptions, implemented with the python libraries *scikit-image.morphology* (Walt et al., 2014) and *scipy.ndimage* (Virtanen et al., 2020). For mitochondria and Golgi apparatus, small objects were removed (less than 27 voxels); for endoplasmic reticulum, holes were removed (less than 20,000 voxels, corresponding to 0.0025 µm^3^); no clean-up was required for nuclear pores or clathrin coated pits.
5. Automatic creation of a coarse voxel-wise mask to mark voxels outside of the cell, using a combination of operations from the python libraries *scikit-image.morphology* and *scipy.ndimage*. The parameters and combination of operations were adapted visually to each dataset. Operations included intensity thresholding, binary opening and closing, filling small holes, and removing small objects.
6. Optional: Correction for systematic biases in annotations. We observed that our semi-automatic annotations carry biases that can be corrected automatically. For mitochondria and Golgi apparatus, most of the annotations did not include the membrane, which we wanted to consider as part of the organelle. Note that this correction depended on the characteristics of a specific dataset (e.g., contrast of membranes): mitochondria annotations were dilated by one voxel (5 nm), Golgi apparatus annotations were dilated by three voxels (15 nm); ER, nuclear pores and clathrin coated pits annotations were not dilated.
7. Defining a metric exclusion zone. Although step (6) allowed us to add most of the organelles’ membrane to the annotation, the ground truth was often not voxel accurate at the organelle boundaries. A neural network model trained with such data cannot produce voxel-accurate predictions at the organelle boundaries, leading to misleading evaluation scores (e. g., F1, see Fig. S3E). Following previous works (Haberl et al., 2020; Lucchi et al., 2012), we avoided this issue by defining an exclusion zone around our semi-automatic imprecise annotations, created by dilating and eroding the annotations and taking the logical difference between the two outcomes. The size of both dilation and erosion depends on the specific structure, as follows — four voxels for mitochondria, two voxels for Golgi apparatuses, one voxel each for endoplasmic reticulum, and three voxels dilation plus one voxel erosion for clathrin coated pits and nuclear pores.

All operations required only local context, meaning that they could be applied block-wise, and the computation could be parallelized to multiple CPU cores. To avoid artefacts at the block boundary, we provided sufficient spatial context to each block with the python library DAISY https://github.com/funkelab/daisy, which was used for multi-process computation on all cores of a CPU. These computations were performed on Intel Xeon workstation processors with 20-40 physical cores. Detailed instructions on the use of the preprocessing pipeline are provided at https://github.com/kirchhausenlab/incasem#Prepare-your-own-ground-truth-annotations-for-fine-tuning-or-training.

### Deep learning

#### Model architecture

A 3D U-Net (Çiçek et al., 2016) based on the architecture used in Funke et al. (2018) was defined, with three downsampling layers with a factor of two, and two convolutional layers on each downsampling level. Refer to Figure S3A for details. In total, the network had ∼6 million parameters. It was implemented in PyTorch (Paszke et al., 2019).

#### Training: Overview of pipeline

The pipeline to feed blocks to the neural network was based on (Buhmann et al., 2021) and implemented using GUNPOWDER [https://github.com/funkey/gunpowder], a library that facilitates machine learning on large multi-dimensional arrays.

We trained one model per organelle; that is, for model training data, *foreground* refers to voxels corresponding to only one type of organelle. For each iteration during the training phase, a block of 204 x 204 x 204 voxels was randomly sampled from the electron microscopy dataset, together with the corresponding block of ground truth. The blocks were augmented by voxel-wise transformations, e.g., random intensity shifts, and geometric transformations, e.g., random rotations and deformations. The blocks were processed through the network, which returned as an output block a 3D probability map of 110 x 110 x 110 voxels, centered with respect to the larger input block. The input blocks contained an additional 47 voxels per side to provide the context required by our convolutional neural network architecture. The output probability map was compared with the ground truth using cross-entropy loss, which was minimized by iteratively updating the model parameters by using the Adam optimizer (Kingma and Ba, 2014).

#### Training: Data sampling

Our dataset annotations were highly imbalanced. Because our structures of interest were small, only a few voxels formed the so-called foreground (FG), with a large portion of the dataset consisting of arbitrary background (BG=1-FG) (e.g., cytosol, nucleus, other organelles). Imbalanced datasets are known to be problematic for convergence of neural network training, and we confirmed this empirically while working with our datasets. As a rule of thumb, it is desirable to sample blocks having a foreground to background ratio roughly equivalent to the global ratio of the two. To make use of all available training data, while keeping the number of unbalanced training blocks as low as possible, we implemented the following scheme (using operations available in GUNPOWDER):

1. Reject blocks that contain more than 25% out-of-sample voxels.
2. Calculate the FG/BG ratio for each incoming block.
3. Reject a block with probability 0.9 if fewer than 5% of the voxels in it (ratio in step (2)) are labeled as foreground.

#### Training: Data augmentation

It was impractical to process and store tens of thousands of augmented FIB-SEM blocks, required to train the 3D neural network model. Instead, during each training cycle, we augmented the number of ground truth annotations by randomly applying the transformations listed in Table S8 to the training block.

#### Training: Pipeline details

After data augmentation, we shifted the scale of the data in the input block (204 x 204 x 204 voxels), such that the input intensities were in [-1, 1]. Each block accepted by the neural network was then propagated through the network leading to outputs of spatial dimensions 110 x 110 x 110 voxels, centered with respect to the larger input block. The neural network assigned complementary FG and BG probability to each voxel. The probability map was then compared to the ground truth annotations with the binary cross-entropy loss. We balanced the loss contribution of foreground and background voxels inversely proportional to their occurrence, clipped at a value of 1:100. The training loss was backpropagated, and the network parameters were updated using the Adam optimizer (Kingma and Ba, 2014) with 0.00003 learning rate and 0.00003 weight decay. The network parameters were saved at the end of every 1000 training iterations.

#### Training: Computational requirements

Eight CPU cores were used in parallel for data fetching and augmentation, while a single GPU (Nvidia A100 on a DGX-A100 system) was used for training. Typically, a training iteration lasted 1 sec, and 100,000-150,000 iterations (28 – 42 h, including periodic validation tests) were sufficient to train our 3D neural network model. A training session could also be done with a standard GPU workstation equipped with a Nvidia GPU with 12 GB GPU memory and 500 GB CPU memory. Presumably workstations with 64 GB CPU memory can also be used since our training pipeline processes out-of-memory data sets.

#### Validation: Procedure

To avoid overfitting, we assessed the model’s performances during the training phase on both training dataset and validation dataset, where the latter dataset was not used to update the model parameters. Every 1,000 iterations we saved the training model, froze its weights, and calculated the loss on a small set of ground truth blocks. These were hold-out blocks from the cells that contained the training data, or blocks originating from naïve cells not represented in the training data. By comparing training and validation losses (see plots in Fig. S3 B-C) we usually identified three different regimes: under-fit, fit and over-fit. When the model under-fitted, both training and validation loss decreased with the training iteration. In the fit regime (Fig. S3 B-C, in grey), typically from more than 20,000 training iterations, the validation loss was approximately constant, while the training loss slightly reduced. In the over-fit regime, the training loss continued to drop, but the validation loss started to rise. We considered the model saved at the training iteration in the middle of the fit regime to be the one that could best generalize better, i.e., make optimal predictions on previously un-seen data. This is a standard procedure in Machine Learning, known as “early stopping”.

#### Validation: Performance metrics

The performance of the models obtained by the 3D U-net neural networks was determined by comparing the predicted binary segmentation with respect to ground-truth using the following three metrics: (1) precision (percentage of voxels predicted as intracellular structure that is the substructure), (2) recall (percentage of substructure voxels correctly predicted as substructure) and (3) F1 index score (harmonic mean of precision and recall, see Fig. S3E). These metrics were also used by other state-of-the-art supervised learning methods, such as COSEM, allowing for a quantitative comparison.

Using the true positives TP, false positives FP and false negatives FN, we define precision, recall and F1 as:

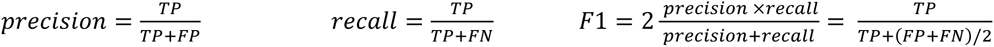

#### Prediction: Data preprocessing

As for training, the segmentation of new FIB-SEM datasets required certain preprocessing steps (refer to *Data preprocessing for deep learning* for details):

1. Conversion from TIFF format to block-wise storage format ZARR.
2. Crop the dataset to exclude empty regions outside the cell.
3. Create an approximate voxel-wise mask to mark voxels outside of the cell.
4. Image data normalization with CLAHE (see Figure S4).

#### Prediction: Pipeline

As first step, the trained model at the iteration determined by early stopping (see *Validation: Procedure*, above) was loaded with frozen weights. The dataset to segment was scanned block by block and fed into the model, without performing data augmentation. Since the architecture of the 3D U-Net neural network is fully convolutional and since each predicted voxel has access to sufficient context, we could produce the predictions block-wise independently and in parallel.

The model performed a forward pass, producing as output a 3D voxel-wise probability map, saved to disk in ZARR format. A threshold of 0.5 was applied to the predicted probability map to extract a binary segmentation map (organelle/background). Although our pipeline allowed the user to set this threshold to an arbitrary value from zero to one, we set it to the default value of 0.5 in every experiment to avoid introducing any post-processing bias. The segmentation map was later visualized in neuroglancer [https://github.com/google/neuroglancer], superimposed on the electron microscopy image. Finally, we converted the predicted segmentations back to TIFF format and reverted the initial dataset cropping to obtain a segmentation that was globally aligned with the originally acquired image stack from the microscope.

#### Prediction: Computational requirements

A prediction was performed on a single GPU (Nvidia A100 on a DGX-A100 system), backed by multiple CPU cores, employed to parallelly load and pre-process the data. When using five CPU cores, we achieve a prediction throughput of approximately 1.6M voxels per second, roughly corresponding to the size of 115^3^ voxels for one block of actual prediction as depicted in our Unet architecture in Fig. S3. Hence the prediction of one cell image stack, acquired at 5 nm isotropic resolution, typically took between 30 minutes and 90 minutes, depending on its volume. A similar prediction could also be done with a standard GPU workstation equipped with a Nvidia GPU with 12 GB GPU memory and 500 GB CPU memory. Presumably workstations with 32 GB CPU memory can also be used since our training pipeline processes out-of-memory data sets.

#### Fine-tuning

To fine-tune a trained model on a naïve cell, we performed the following steps:

1. One block of ground-truth block (minimum 204 x 204 x204 voxels) within the cell to fine-tune was annotated.
2. A model previously trained on the same organelle, whose weights were frozen at the first train iteration of the fit region, was loaded for continued training.
3. A training of 15’000 iterations was launched, using as training data only the new prepared ground-truth block. All fine-tuning training hyperparameters were set identical to the original training.
4. The model was saved every 1’000 iterations. The best model iteration was picked based on the original validation dataset from the fine-tuning target cell.

Details to perform fine-tuning training using our pipeline are provided at https://github.com/kirchhausenlab/incasem#Fine-tuning.

### Brief example of fine-tuning

We started by preparing a small ground-truth block of the cell to fine-tune. Because the volume of the additional ground truth was much smaller than the volume of the ground truth fed to the pre-trained model (from ∼1/69 to ∼1/3, Table S10), the additional annotation effort was not very demanding. In the case of CF datasets, for the fine-tuning of mitochondria in Cell 3 BSC-1 and Cell 6 SUM 159, we loaded the segmentation map performed by the pre-trained model (model 1847) and refined it by manual editing with VAST. Conversely, to fine-tune HPFS OpenOrganelle datasets in the prediction of mitochondria and ER, we picked one additional COSEM ground-truth crop, previously excluded from the training and validation data. We loaded the pre-trained model at the first training iteration within the fit regime (i.e., the iteration after which the validation loss was stable, Figure S3 B-C); for example, in Figure S5A, CF Mitochondria model, we loaded the model at the training iteration 95,000. We resumed training of the model using only the additional ground truth block and data augmentation. Typically, after a few thousand training iterations (about ∼1/15 of the training iterations needed to produce the pre-trained models), the fine-tuned model learned to segment the fine-tune dataset more precisely, with an increased overall F1 score (Table S10). We noticed that fine-tuning was beneficial mainly when the initially trained model behaved poorly, such as in the case of HPFS ER, Cell 21 Jurkat-1, for which F1 increased by 0.21.

To understand how fine-tune training helped to improve segmentation, we compared the segmentation performed by the pre-trained model versus the one produced by the fine-tuned model (Figure 5 D). In the case of CF, mitochondria, only a few portions of mitochondria were occasionally missed by the pre-trained model, but they were eventually retrieved by the fine-tuned models. In HPFS ER, fine-tuning reduced the number of false positives, resulting in a neater segmentation map, suitable for further biological studies.

To quantify the impact of fine-tune training, we calculated precision and recall in addition to F1 (Figure S5). Typically, fine-tuning enhanced precision, without affecting recall: the fine-tuned model learned to classify cell components that looked like the organelle under study, but did not actually belong to the same semantic class, reducing the number of false positives.

As baseline, we compared the fine-tuned model with one having randomly initialized weights and trained using the same (small) ground truth volume. We found that for most datasets and organelles (ER Cell 21 Jurkat-1, ER Cell 22 Macrophage-2, mitochondria Cell 21 Jurkat-1, mitochondria Cell 3 BSC-1, mitochondria Cell 6 SUM 159), the models trained using randomly initialized weight reached substantially lower F1 scores, even when trained much longer, with up to 200,000 training iterations. Only in one case (mitochondria Macrophage-2), did the model achieved approximately the same F1 score, but only after training for 160,000 iterations, 20 times more than the number required by the corresponding fine-tuned model.

We concluded that fine-tune training is a useful tool to apply whenever the segmentation of a naïve cell falls short. By taking advantage of pre-trained models and preparing a small ground truth volume, one can train more accurate models, at a small fraction of the ground truth annotation and computational costs usually required.

### Estimate of time effort required to annotate ground truths

We evaluated our annotation times for all cells and organelle type (Table S4). The total annotation time for the given volumes were 120 h for 176 um^3^ (mitochondria), 100 h for 166 um^3^ (Golgi apparatus), and 120 h for 92 um^3^ (ER). On average, this equates to roughly 0.8 h per cubic micron per organelle for ASEM. In contrast, the COSEM paper (Heinrich et al., 2021) states that annotating a block of 1 cubic microns with 35 organelle classes took one annotator two weeks (∼ 100 h), which equates to 2.8 h per cubic micron per organelle, thus being ∼ 3.5 times slower than ASEM.

### Analysis of nuclear pores

#### Automated orientation of nuclear pores on the nuclear envelope

We determined the nuclear pore membrane diameters from top-down FIB-SEM views towards the nuclear envelope of each of the nuclear pores. This orientation process was automated with the following steps. First, we generated a 3D binary mask corresponding to the predicted probability with a threshold of 0.5 for each one of the nuclear pores identified by the 3D U-net nuclear pore model (Fig. S6A). Second, we determined for each mask its volume, principal axis, and centroid coordinates. A filtering step was included to eliminate masks with small volume or short axis due to incompleteness of the predicted mask. Third, we used median filtering of the 3D point cloud data to remove outliers, thus creating a virtual ‘low resolution’ 3D surface of the nuclear envelope by alpha-shape triangulation of the centroids (Akkiraju et al.) (Fig. S6B, top). Fourth, we obtained a vector normal to each triangle within the triangulation (Fig. S6B, bottom). Finally, we used the angular information of this vector to reorient the coordinates of the raw image of the nuclear pore closest to the triangulation to position the nuclear membrane on a view normal to the observer (Fig. S6C).

#### Determination of nuclear pore membrane diameter

The diameter of the membrane pore, defined by the contact between the nuclear membrane and the pore opening, was determined from the distance separating the two peak signals measured in the FIB-SEM image along a line transecting the middle of the nuclear envelope immediately surrounding the nuclear pore (d1 and d2 in Fig. S6D). The nuclear pore membrane diameter was expressed as the median of eighteen radial measurements 10° apart. This calculation increased the precision of the measurement by taking advantage of the known radial symmetry of the nuclear pore and the surrounding nuclear envelope on the axis normal to the nuclear envelope; the standard deviation for each pore diameter measurement (i.e., experimentally determined uncertainty) was 6 nm.

### Analysis of clathrin coated pits and coated vesicles

The model trained on endocytic clathrin coated pits in cell 12 and cell 13 was used to predict clathrin coated structures in cells 12, 13, 15, and 17. The predictions were gated at a probability of 0.5 and the corresponding masks then used to locate the clathrin coated structures. These structures were classified as endocytic clathrin coated pits and coated vesicles if located at the plasma membrane or within 400 nm, respectively; ‘secretory’ coated pits and coated vesicles denote the remaining similar structures located in the cell interior. Each prediction was confirmed by visually inspection of the corresponding image along the three orthogonal directions.

Measurements of neck, height, and width from the pits (Fig. S7A) and major and minor axis of the ellipses best fitting the pits and coated vesicles (Fig. S7A,B) were determined by the following sequential steps: (1) select the view displaying the largest outline by inspection of 9 consecutive planes along each of the three orthogonal views centered on the centroid of the pit or vesicles; (2) manually measure the neck height and widths of the pits; (3) establish the outline of the pit or vesicle in the section chosen in the first step; the darker pixels where the pit or vesicle was present were selected manually (Fig. S7C, white square in the left panel), segmented into a binary mask with an Otsu intensity threshold (Otsu, 1979), and skeletonized (Fig. S7C, middle panel); (4) establish the ellipse best fitting the skeletonized outline of the pit or vesicle (Fig. S7C, right panel); (5) obtain major and minor axis of the ellipse.

### Statistical analysis

The normality of the nuclear pore size distribution was examined using the Shapiro-Wilk test (Fig. 6C). The comparison of size distributions between nuclear pore diameters from values determined experimentally and from simulated values based on the experimental median value with an uncertainty of 6 nm using the nonparametric Kolmogorov-Smirnov test showed they were statistically different (p<0.0001).

### Data availability

The datasets of raw and normalized FIBSEM cells images, ground truth annotations, probability maps predicted by the models and corresponding segmentation masks are publicly available at the AWS ASEM bucket [https://open.quiltdata.com/b/asem-project].

### Code availability

The software and step-by-step instructions to use it are publicly available at https://github.com/kirchhausenlab/gc_segment (Graph-cut annotation tool) and at https://github.com/kirchhausenlab/incasem (Deep-learning pipeline).

Trained neural network models are available at https://open.quiltdata.com/b/asem-project with usage instructions at https://github.com/kirchhausenlab/incasem.

## Figure and Video legends

**Video 1. Ground truth annotations for mitochondria, ER and Golgi apparatus**

Passing through a FIB-SEM volume with contrast equalized using CLAHE. Image is from Cell 1 HEK293A prepared by CF and imaged at ∼ 5 nm isotropic resolution. The video shows ground truth annotations for mitochondria (cyan), ER (red) and Golgi apparatus (green). The annotations were generated for all mitochondria and Golgi apparatus within the FIB-SEM volume, and all ER within the highlighted 8 µm x 3 µm x 3 µm block (orange box).

**Video 2. Predictions of mitochondria**

Passing through the FIB-SEM volume with contrast was equalized using CLAHE. The video shows image and predictions from naïve Cell 1 HEK293A (not used for training using the model trained with ground truth annotations for mitochondria from Cell 2 HEK293A. Both cells were prepared by CF and imaged at ∼ 5 nm isotropic resolution. The model identified all mitochondria; comparison of the ground truth annotations and predictions shows correct voxel assignments (true positives, yellow), missed assignments (false negatives, cyan), incorrect assignments (false positives, magenta). The small fraction of false positive assignments predicted by the model are associated with unidentified tubular and spherical structures of small size.

(Chou et al., 2021)

**Video 3. Prediction of mitochondria, ER, Golgi apparatus, nuclear pores, and clathrin coated pits and vesicles**

Passing through the raw FIB-SEM volume from naïve Cell 15 SVG-A prepared by HPFS and imaged at ∼ 5 nm isotropic resolution. The video shows predictions as surface renderings for mitochondria (cyan), ER (yellow), Golgi apparatus (magenta). For simplicity, only predictions in a block of 3 µm x 3 µm x 3 µm (block size: 664 x 586 x 572 voxels) are shown. A small number of false positive pixels generated by the Golgi model and located within a 323 x 271 x 230 voxel block were removed using VAST. One identified Golgi apparatus is highlighted (light pink). Predictions for all nuclear pores (yellow) and clathrin coated pits and vesicles within the imaged volume are also shown. Visual inspection confirmed that the models trained with ground truth annotations from Cell 19 HeLa and Cell 20 HeLa prepared by HPFS and imaged at ∼ 5 nm isotropic resolution correctly predicted all the intracellular structures in Cell 15 SVG-A.

**Video 4. Prediction of mitotic ER**

Passing through the FIB-SEM volume with contrast equalized. Image is from naïve prometaphase Cell 8 SUM 159 imaged at ∼ 10 nm isotropic resolution. The video shows ER predictions (yellow) generated with ground truth annotations from interphase Cell 1 HEK293A and Cell 2 HEK293A imaged at ∼ 5 nm isotropic resolution. All cells were prepared by CF. Visual inspection confirmed that the model correctly predicted all the ER, including the fenestrations characteristic of the mitotic ER sheets; fenestrations were not included in the ground truths used for training, as they are mostly absent in the ER of interphase cells (Chou et al., 2021).

## Supplemental material

Supplemental material consists of Figs. S1-S10 and Tables S1-S10.

## Acknowledgments

We thank Jan Funke for initial guidance on the use of deep learning and for maintaining together with members of his laboratory the gunpowder and funlib libraries; Jose Inacio Da Costa Filho for testing our public available codes; Arlo Sheridan for helpful discussions; Teresa Rodriguez and Henrique Girão for providing cells 1 and 2; Giovanni de Nola and Teresa Rodriguez for preparing Cells 1-4 for FIB-SEM imaging; Rasmus Herlo and Max Page for preparing Cell 5 for FIB-SEM imaging; Justin Houser for preparing Cell 6a-9 for FIB-SEM imaging; Gleb Shtengel and C. Shan Xu for preparing Cells10-13 for FIB-SEM imaging; and S.C. Harrison for extensive editorial help; and members of the Kirchhausen laboratory for help and encouragement.

The research was supported by a National Institute of General Medical Sciences Maximizing Investigators’ Research Award GM130386 and a generous grant from SANA to T.K. B.G. and G. M. were supported in part by discretionary funds available to T.K. Acquisition of the FIB-SEM microscope was supported by a generous grant from Biogen to T. K. Acquisition of the computing hardware including the DGX’s GPU-based computers, CPU clusters, fast access memory, archival servers and workstations that made possible this study were supported by generous grants from the Massachusetts Life Sciences Center to T.K. and an equipment supplement to the National Institute of General Medical Sciences Maximizing Investigators’ Research Award GM130386. Construction of the server room housing the computing hardware was made possible with generous support from the PCMM Program at Boston Children’s Hospital. B.G. & M.W. were supported by the EPFL School of Life Sciences and by generous funding from CARIGEST SA.

The authors declare no competing financial interests.

## Author contributions

B. Gallusser, Mihir Sahasrabudhe and T. Kirchhausen conceptualized the computational aspects of the project; B. Gallusser, G. Maltese, G. Di Caprio, and M. Sahasrabudhe created, implemented, and used the computational pipeline to train the neural network, obtain models and make predictions; J. O’Connor helped set up and maintain the computing infrastructure; T. J. Vadakkan, B. Gallusser, G. Maltese and G. Di Caprio generated and curated the ground truth annotations used for training. G. Di Caprio analyzed the data on nuclear pores and coated structures. Anwesha Sanyal generated the cell lines used to image Cells 13a-17; Anwesha Sanyal and Elliott Somerville prepared Cells 13a-17 for FIB-SEM imaging. T. J. Vadakkan used FIB-SEM to image most cells used in this study (Cells 1-17); T. Kirchhausen supervised the imaging experiments; B. Gallusser, G. Di Caprio, G. Maltese, M. Sahasrabudhe, M. Weigert, and T. Kirchhausen participated drafting the manuscript; T. Kirchhausen contributed to the final manuscript in consultation with the authors.

---

**Figure S1.**
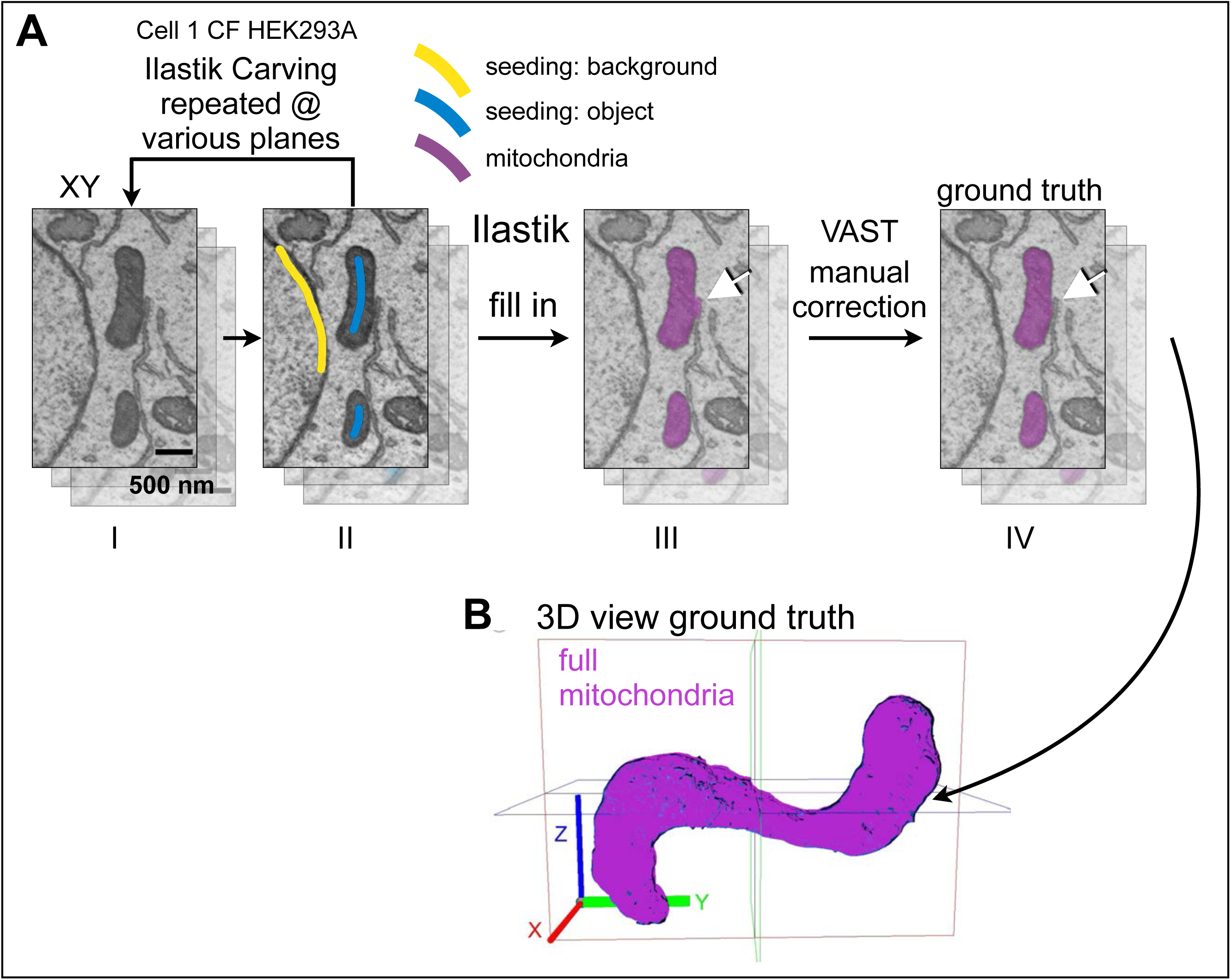
Ground truth annotation workflow for mitochondria. **(A)** Example to illustrate the sequential steps used with Ilastik Carving module to generate the ground truth annotation for a mitochondrion in Cell 1 HEK293A prepared by chemical fixation and visualized with ∼ 5 nm isotropic resolution. Coarse annotations for background (yellow) and object (blue) drawn in broadly spaced consecutive planes of the stack were used to seed the Ilastik Carving module from which a binary mask spaced along adjacent planes spaced 5 nm in the z-stack and corresponding to the mitochondria ground annotation was generated (magenta). Manual corrections using VAST are used as needed, to remove incorrectly assigned pixels, in this example corresponding to an adjacent ER (white arrow). **(B)** Volume rendering corresponding to the ground truth annotation of the mitochondrion shown in **(A)**. Scale bar, 500 nm.

**Figure S2.**
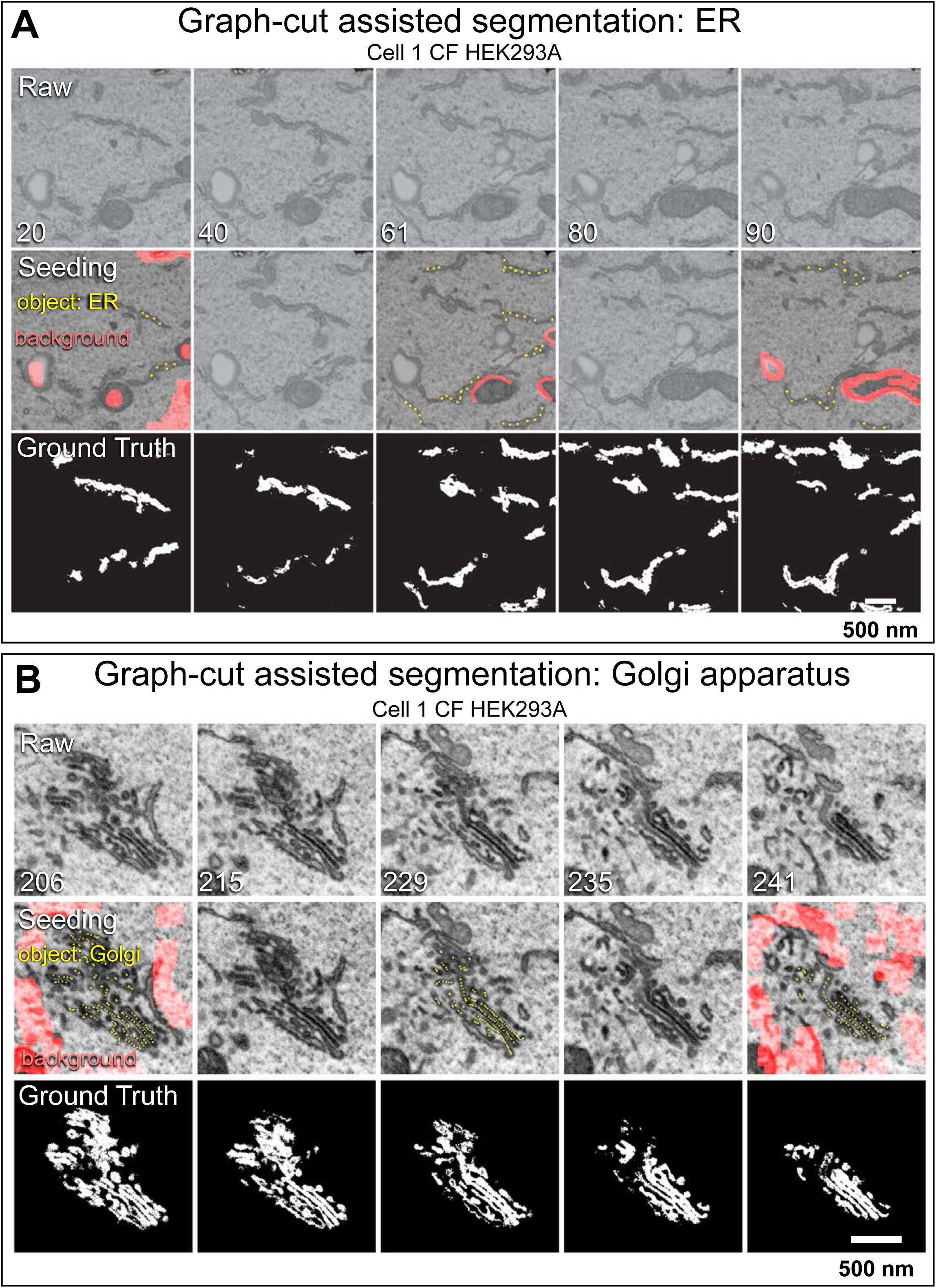
Ground truth annotation workflow for ER and Golgi apparatus. **(A, B)** Example of graph-cut assisted segmentation used to generate the ground truth annotation for ER **(A)** or Golgi apparatus **(B**) in Cell 1 HEK293A prepared by chemical fixation and visualized with ∼ 5 nm isotropic resolution. Coarse annotations for background (lines, solid areas in pink) and object (dotted lines in yellow) drawn in the indicated broadly spaced planes of the stack were used as seeds to obtain the ground truth annotations spaced 5 nm apart generated by the graph-cut assisted segmentation program.

**Figure S3.**
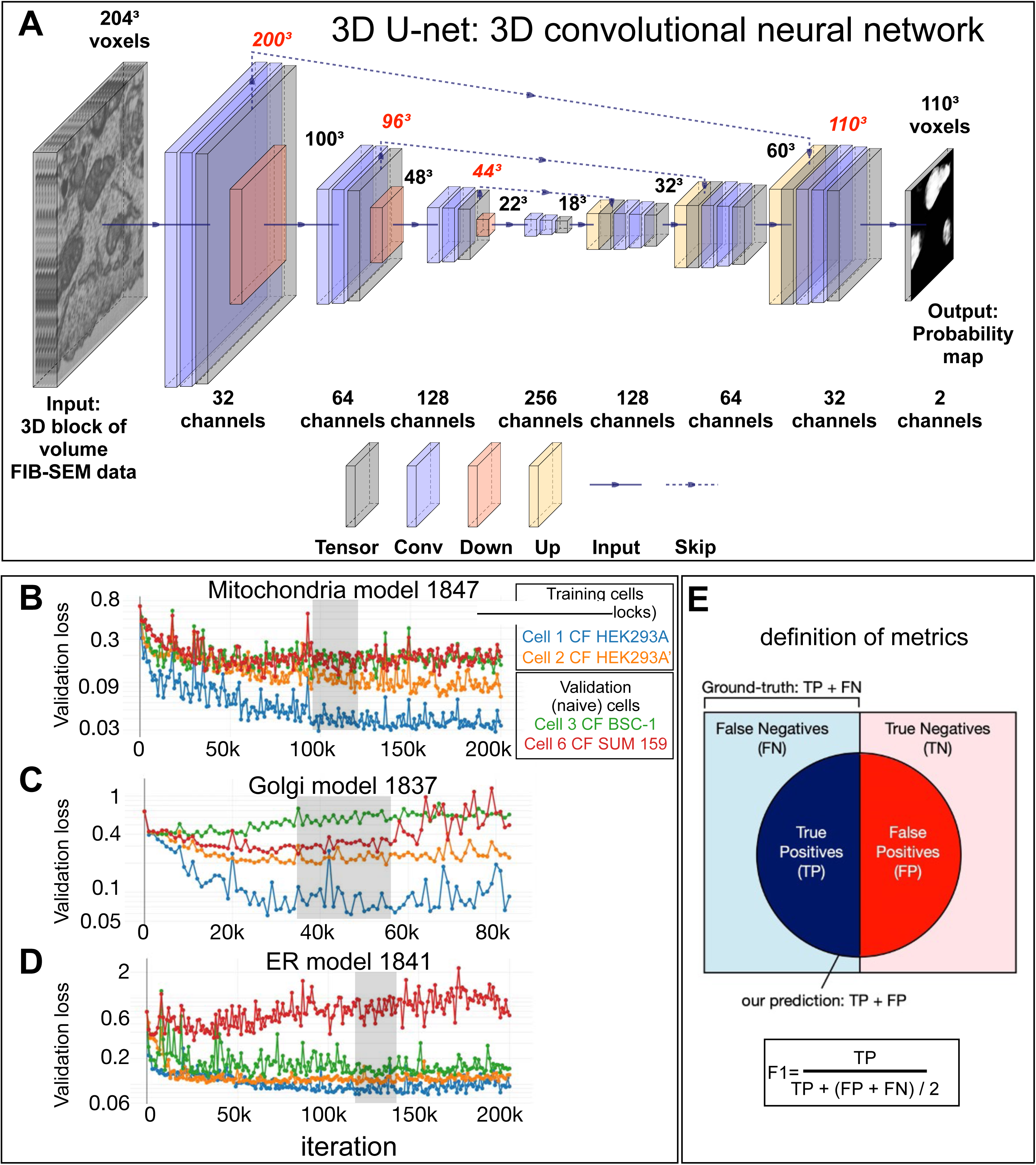
3D-Unet architecture, examples of network behavior during training, and F1 as a metric to compare ground truth annotations with model predictions. **(A)** Schematic representation of the steps used to train the 3D U-net encoder-decoder neural network. The input for the neural network mode are 3D blocks consisting of a stack of consecutive FIB-SEM images (size 204 x 204 x 204 voxels). The 3D block is subjected to consecutive 3 x 3 x 3 convolutions without padding (purple) and down sampling operators with 2 x 2x 2 max-pooling (pink), followed by consecutive up sampling by a factor of 2 (yellow) of the feature maps. During up sampling, the feature maps are concatenated with previous feature maps from the down sampling branch that had been exposed to central cropping; this step also includes consecutive 3 x 3 x 3 convolutions without padding (purple). The output of the neural network model is a feature map (size 110 x 110 x 110 voxels) of two channels, representing the foreground (FG) and background (BG = 1-FG) probability maps, respectively. Number of featured maps are denoted in red, spatial dimensions at the indicated steps in the neural network, in black. Figure designed based on PlotNeuralNet (https://github.com/HarisIqbal88/PlotNeuralNet) (adapted from (Sheridan et al., 2022). **(B-D)** Examples of plots of cross entropy loss used to evaluate the predicting behavior of the indicated neural network models for **(B)** Mitochondria, **(C)** Golgi or **(D)** ER obtained during training using FIB-SEM volume data of cells prepared by chemical fixation obtained at ∼ 5 nm resolution. Cross entropy values were obtained using ground truth annotations from the training set or from naïve cells not used during training, respectively. The gray area shows the first appearance of relatively stable cross-entropy loss and absence of major spikes obtained by the models during 20,000 consecutive training iterations; these models were then used to evaluate their network architecture and prediction performance. **(E)** Ground truth annotations consist of true positive (TP) and false negatives (FN) voxels and define presence or absence of a perfect match with the subcellular structure of interest. The model predicts voxels with true (TP) and false positives (FP) values, depending on whether it considers them as representing or not the structure of interest. F1, as defined in the figure, is used as a practical metric to evaluate the prediction accuracy of the neural network to identify the structure of interest. A perfect model prediction would yield F1=1 with FP=0, FN=0.

**Figure S4.**
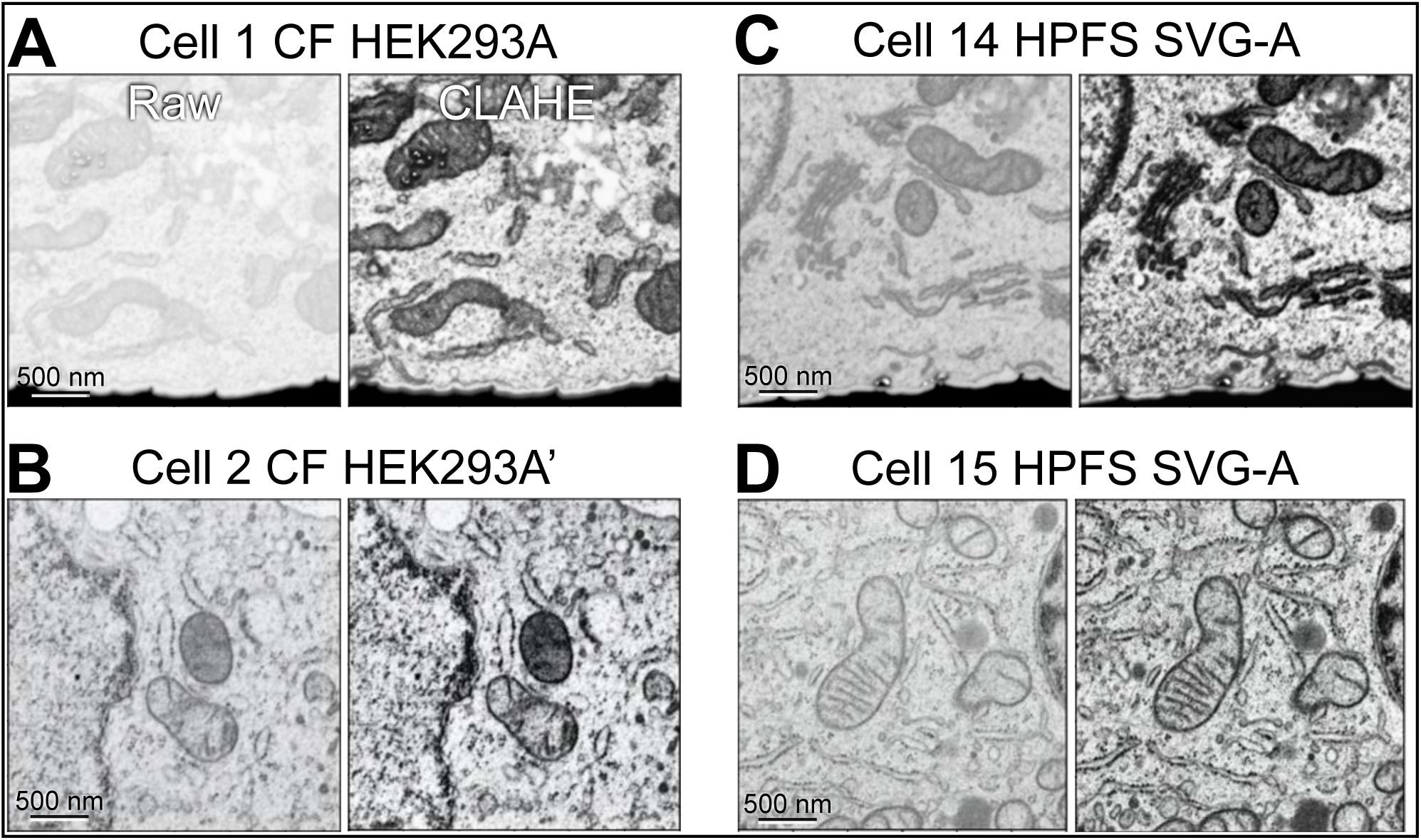
Use of CLAHE to equalize the contrast of FIB-SEM images. **(A-D)** Single plane views of FIB-SEM volume data after contrast equalization using CLAHE with a clip limit of 0.02. The samples were prepared by CF **(A, B)** or HFFS **(C, D)** and imaged at ∼ 5 nm isotropic resolution.

**Figure S5.**
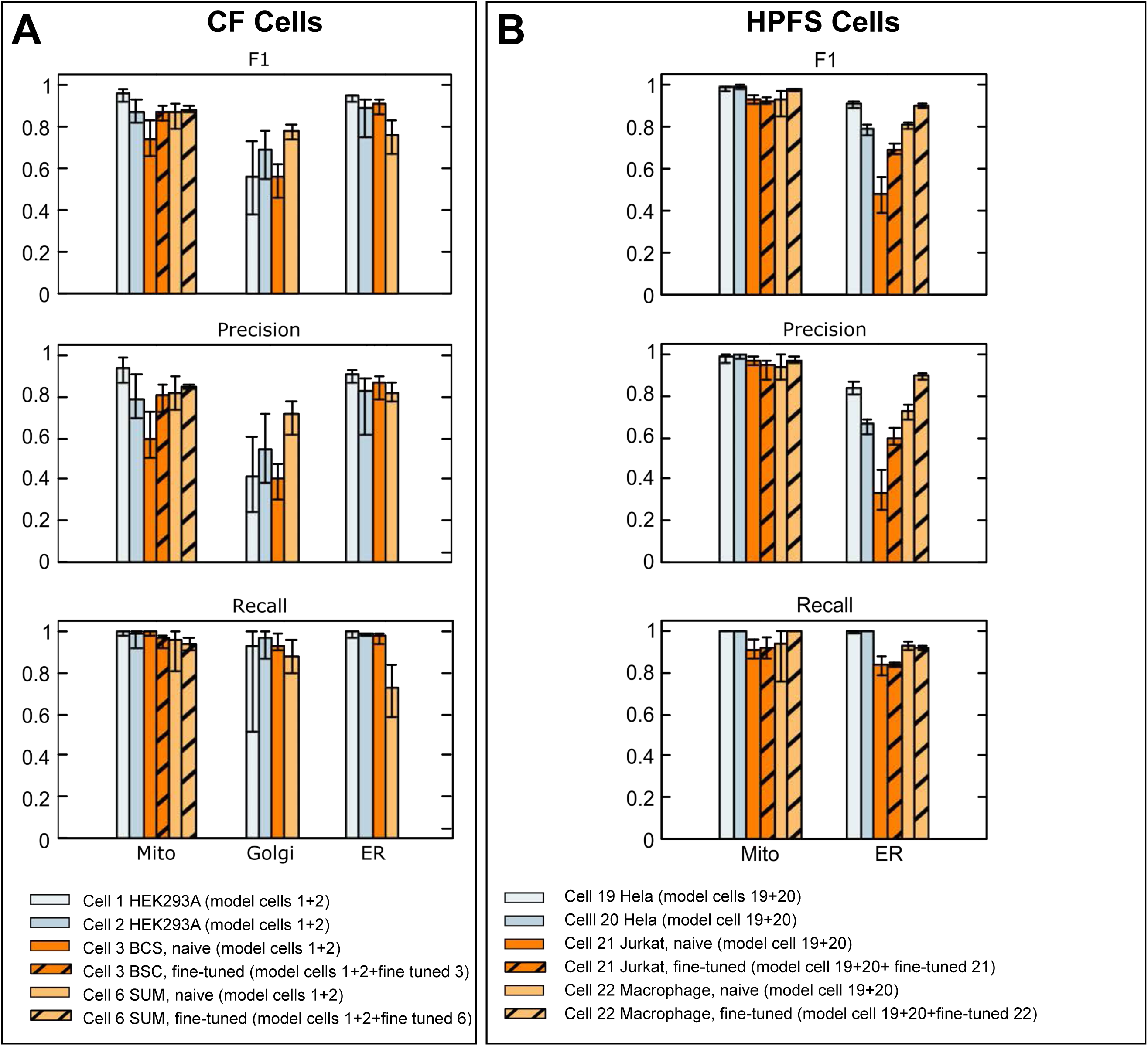
Comparison of metrics used to validate the prediction accuracy of neural models predicting mitochondria, ER and Golgi apparatus. Ground truth annotations from FIB-SEM volume data from the indicated cells at ∼ 5 nm isotropic resolution prepared by CF or HPFS were used for training to generate models for mitochondria, ER and Golgi apparatus. The histogram plots show F1, precision and recall metrics obtained using ground truth annotations not used for training. The results also show metrics obtained after fine-tunning with a small number of additional training iterations using ground truth annotations from the naïve cell. Details of datasets, ground-truth annotations and models are summarized in Tables S4, S5 and S2.

**Figure S6.**
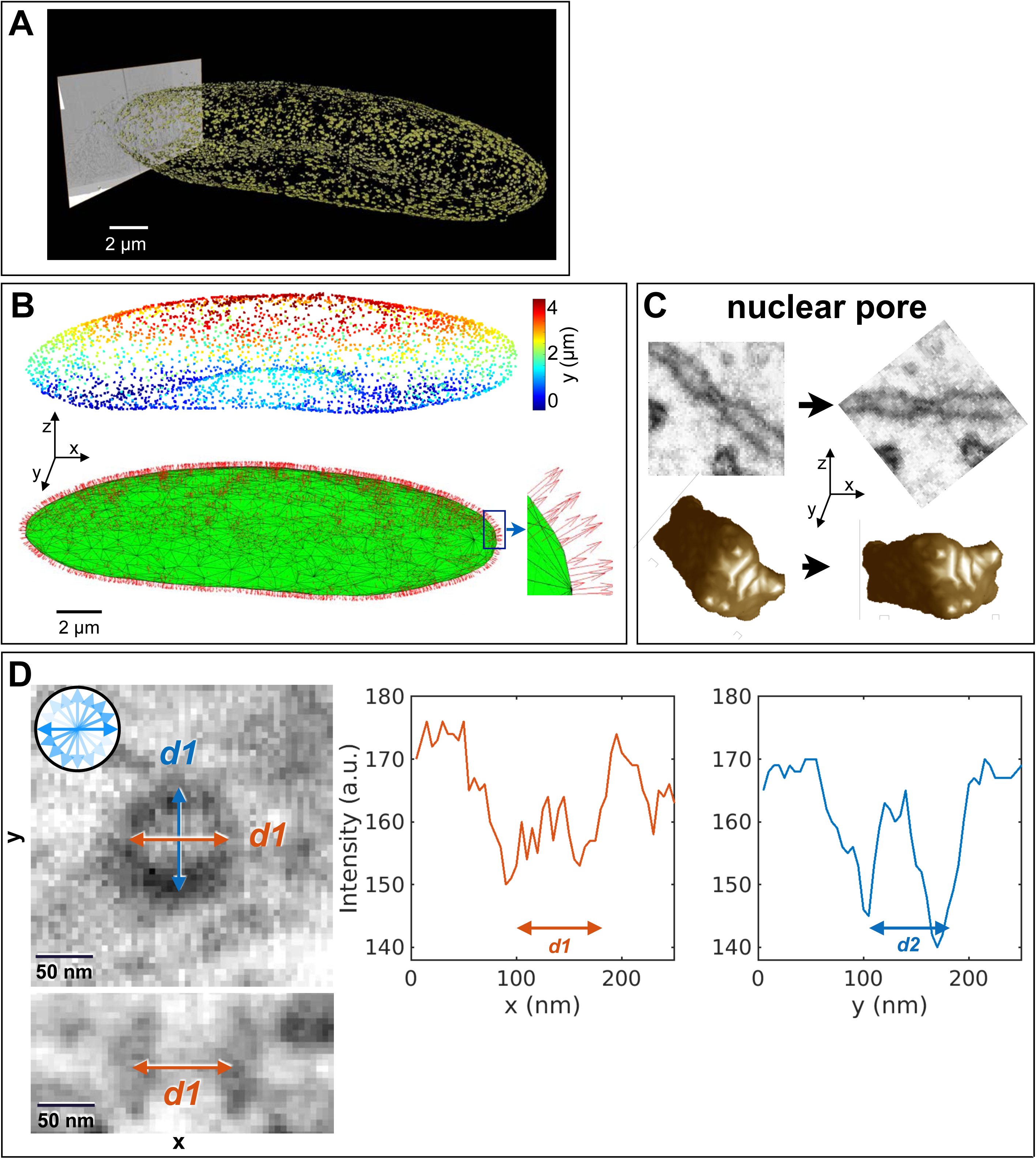
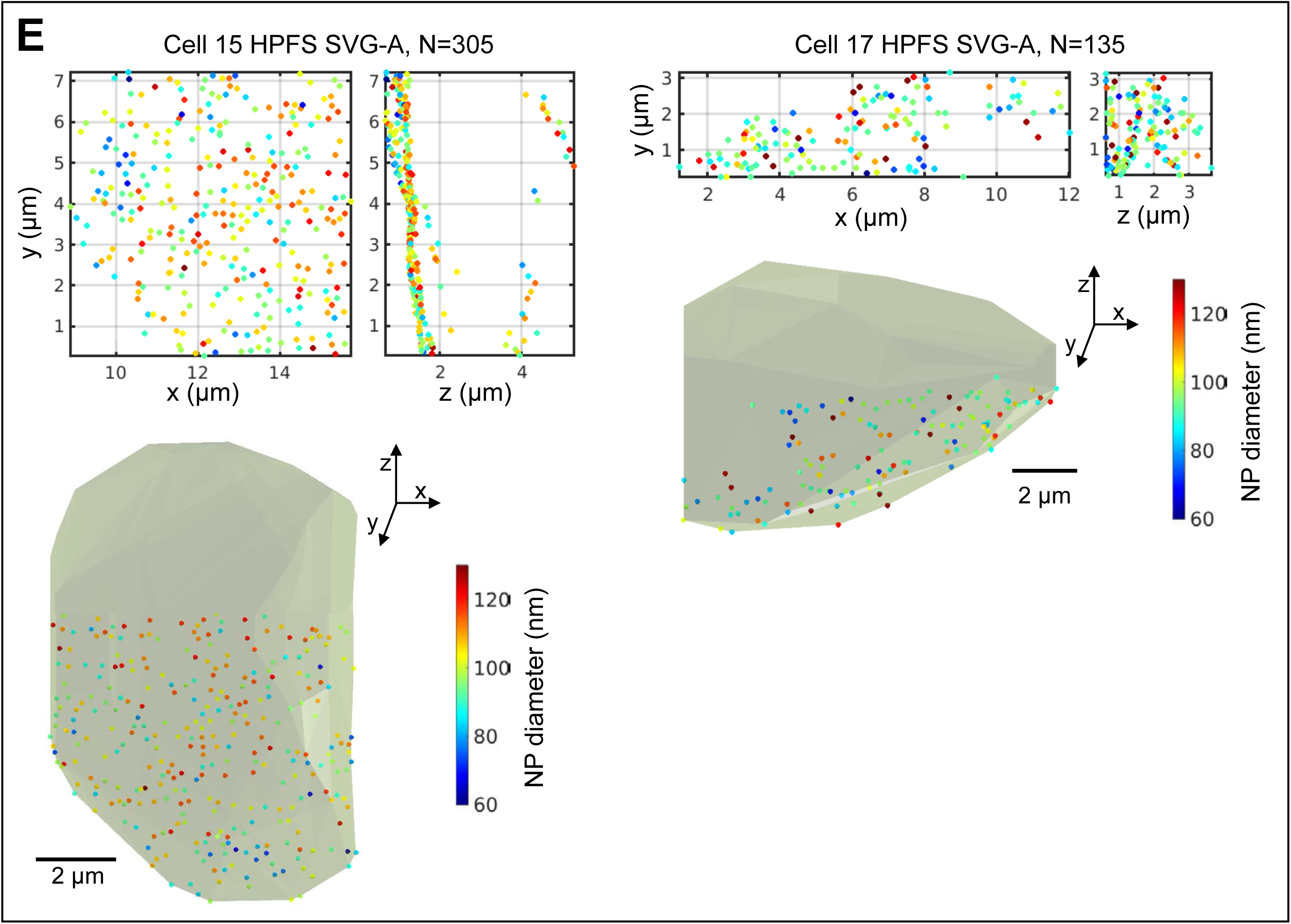
Steps to determine the diameter of the nuclear pore membrane. **(A)** Nuclear pore predictions for all the pores on the nuclear envelope of naïve interphase cell 19 (Hela-2) prepared by HPFS and visualized at 4 x 4 x 5.3 nm isotropic resolution. The nuclear pore predictions were obtained using model 1986 trained without fine tuning with ground truths annotations for Cell 13 (Hela) prepared by HPFS and imaged at ∼ 5 nm isotropic resolution. **(B)** Volume location of the centroid of each of the predicted nuclear pore’s color coded according to their relative position along the Z-axis (top panel) and surface rendition of the nuclear envelope (green) obtain by alpha-shape triangulation of the centroids (see Methods). Orthonormal vectors associated with each triangle are shown (red). **(C)** Example of realignment of a nuclear pore from its acquisition orientation in the FIB-SEM volume image to a new view with the nuclear envelope orthogonal to the Z-axis; side views and volume rendition of the nuclear pore prediction are shown. **(D)** Single plane on face and orthogonal views of a nuclear pore centered on the middle of the nuclear envelope (left panels) and examples of the intensity plots used to estimate the membrane pore diameters by determining the distance separating the two intensity minima along the indicated axis (right panels). The nuclear pore diameter is reported as the average of 10 values obtained 18 ° apart (inset in left panel). **(E)** Three-dimensional distribution of nuclear pores on the nuclear envelopes of Cells 15 and 17 color coded by a heat map as a function of membrane pore diameter.

**Figure S7.**
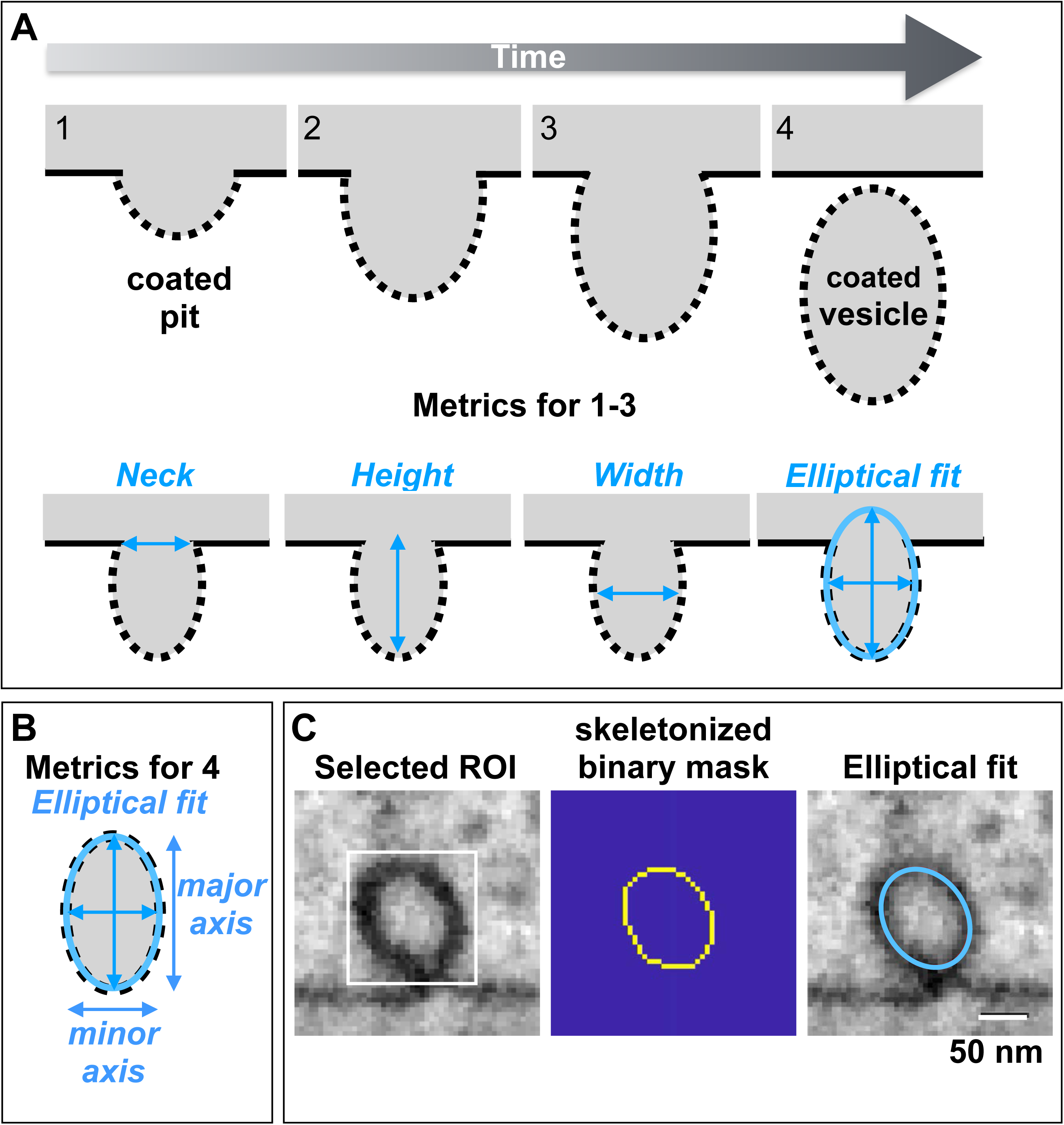
Definition of metrics used to characterize clathrin coated structures. **(A)** Schematic representation of the timeline to describe the formation of a clathrin coated pit mediated by the assembly of the clathrin coat (Kirchhausen et al., 2014). The last step mediated by fission of the membrane neck connecting the mature coated pit from the originating membrane results in formation of the fully formed coated vesicle. Metrics of neck width, pit height, full width at half maximum and major and minor axis of the fitted ellipse used to morphologically describe the clathrin coated pits are shown. **(B)** Metrics used to characterize clathrin coated vesicles. **(C)** Example of a single plane from a selected endocytic clathrin coated pit in a cell prepared by HPFS and imaged by FIB-SEM at ∼ 5 nm isotropic resolution. The darker voxels corresponding to the deformed membrane and the coat surrounding the pit (left panel) were segmented using an Otsu-based intensity threshold approach (Otsu, 1979) to generate a skeletonized binary mask (central panel) which was then used to fit the ellipse (right panel).

**Figure S8.**
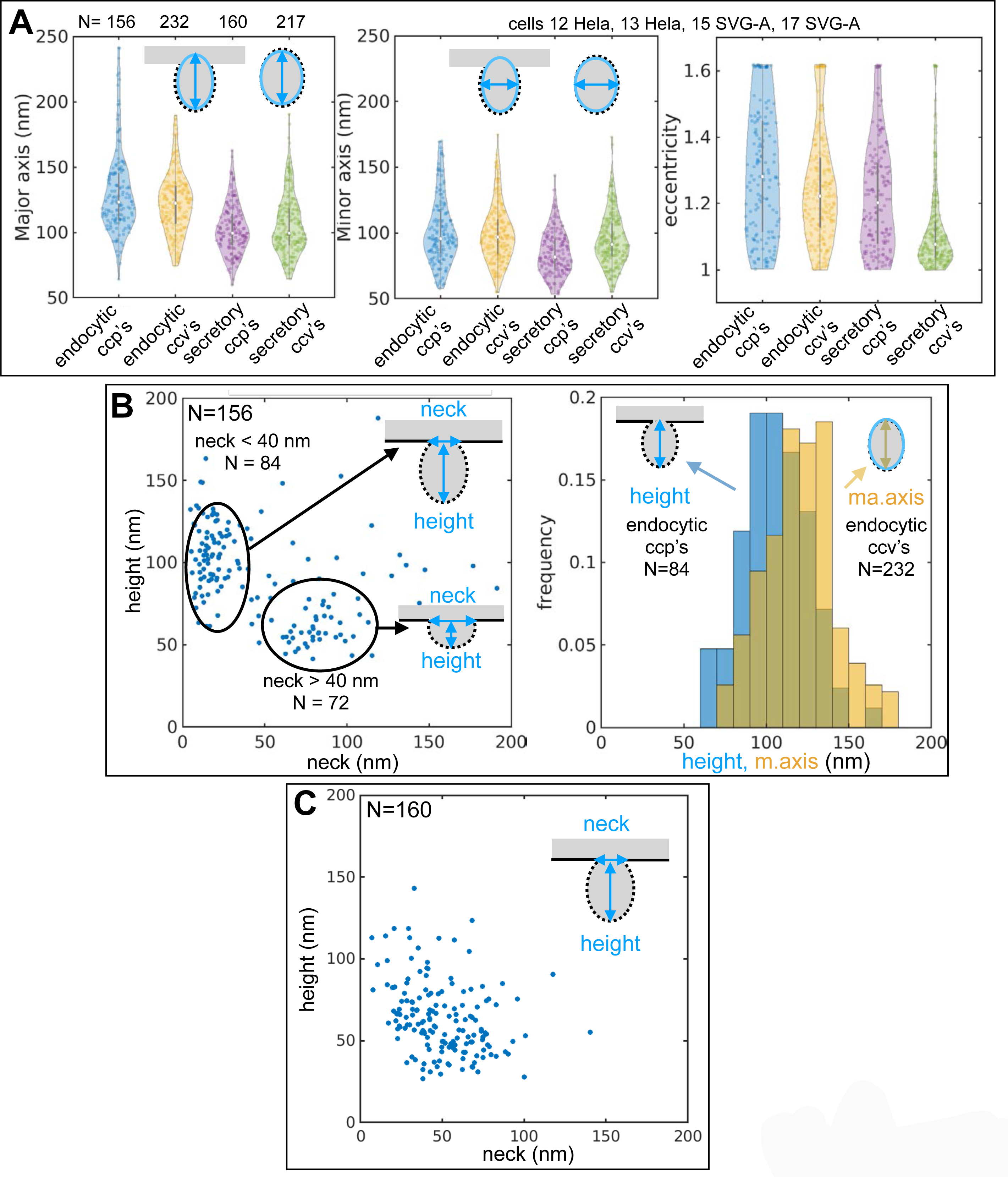
Characterization of clathrin coated pits and coated vesicles. Data shown in this figure for Cells 12, 13, 17 and 17 were generated using the coated pit model employed in Fig 7 obtained by training with ground truth annotations from Cell 12 prepared by HPFS and imaged at ∼ 5 nm isotropic resolution. **(A)** Violin plots of major and minor axis and eccentricity of the fitted ellipse of all pits and vesicles in the raw images of the structures identified by the coated pit model. **(B)** Scatter plot of height versus neck width of endocytic clathrin coated pits clustered in two groups associated with early and late stages of pit formation (left panel). The histogram compares height and major axis for the fitted ellipse of late endocytic coated pits and coated vesicles, respectively. **(C)** Scatter plot of height versus neck width of ‘secretory’ clathrin coated pits associated with internal membranes.

**Table S1.**
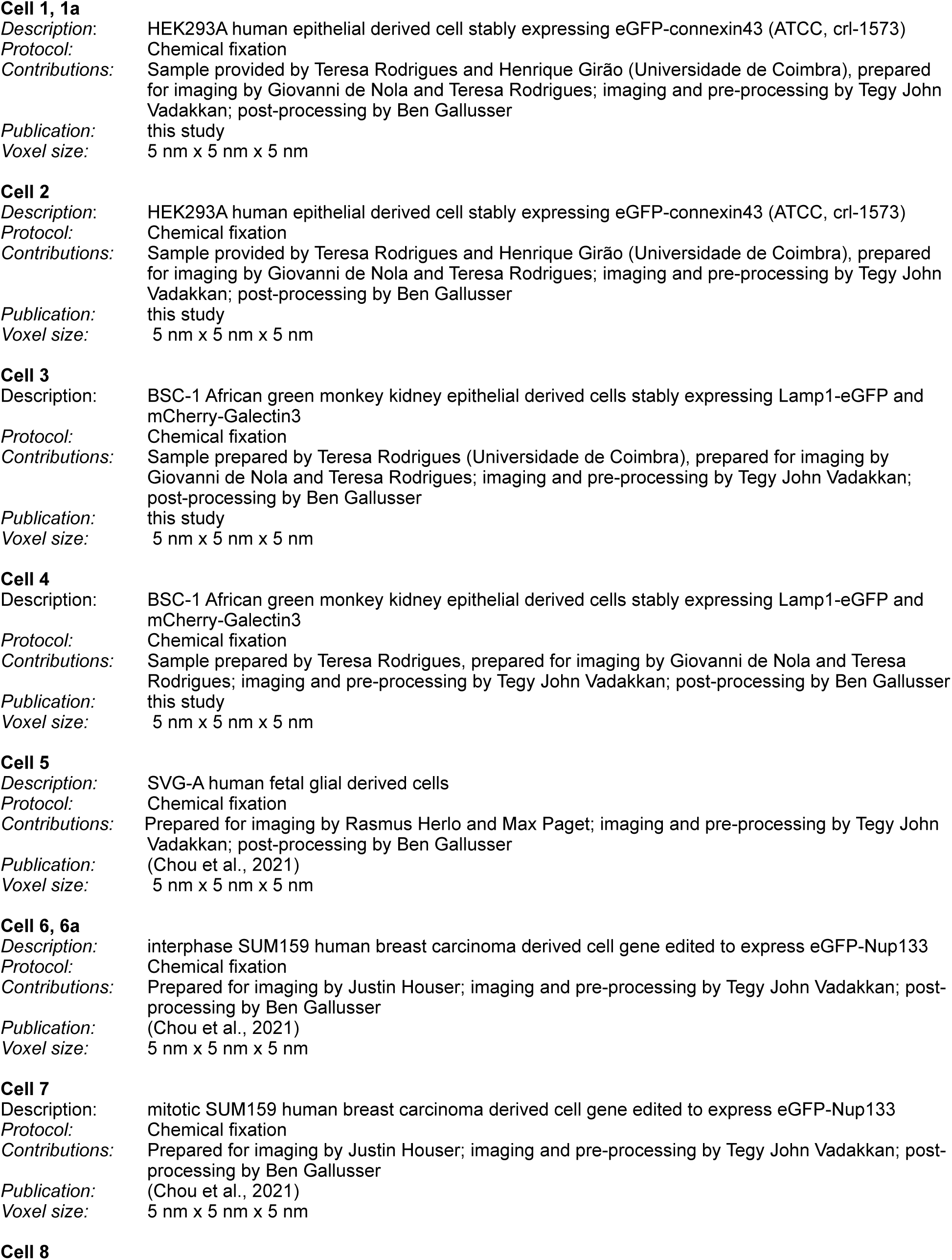
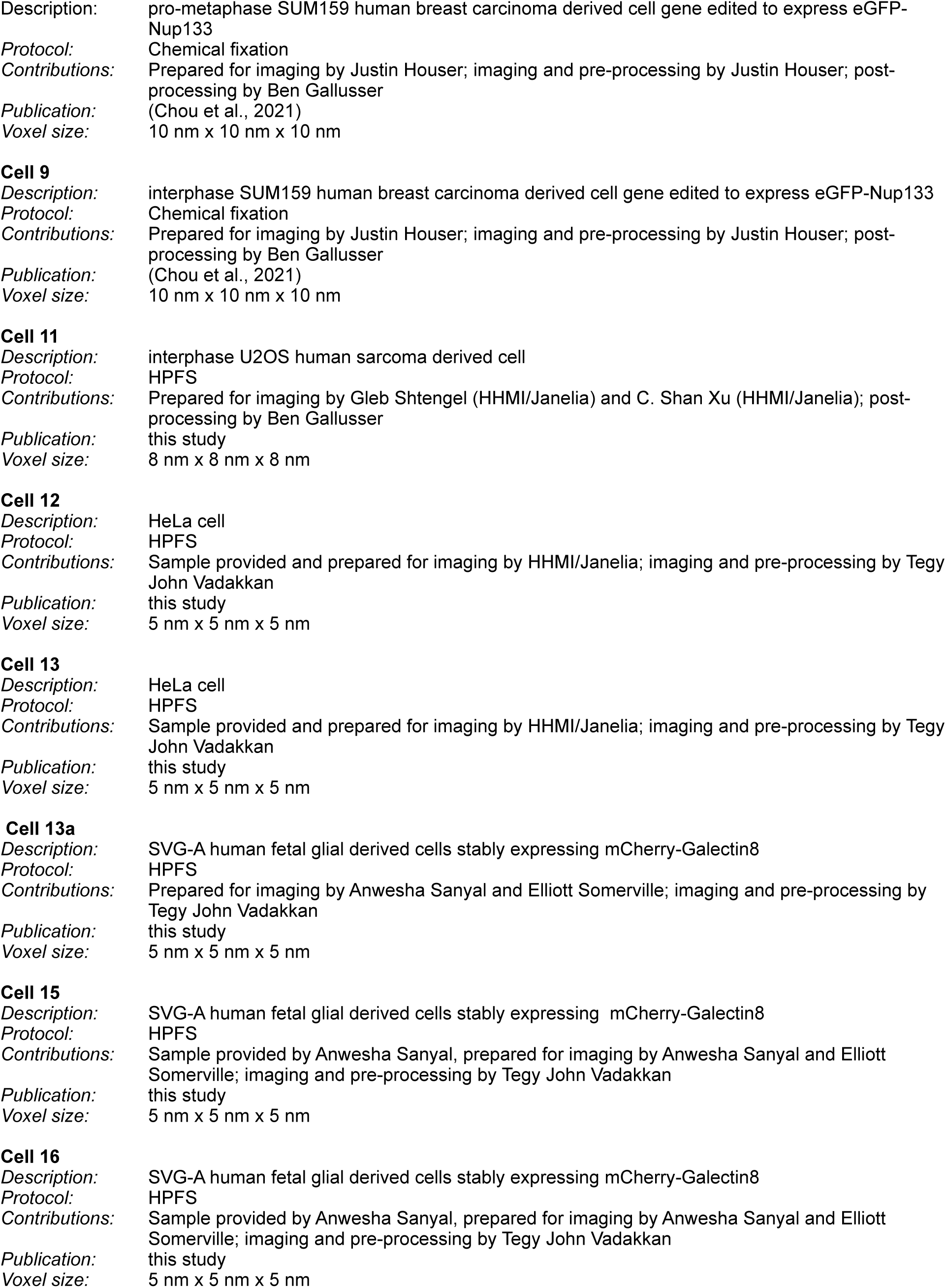
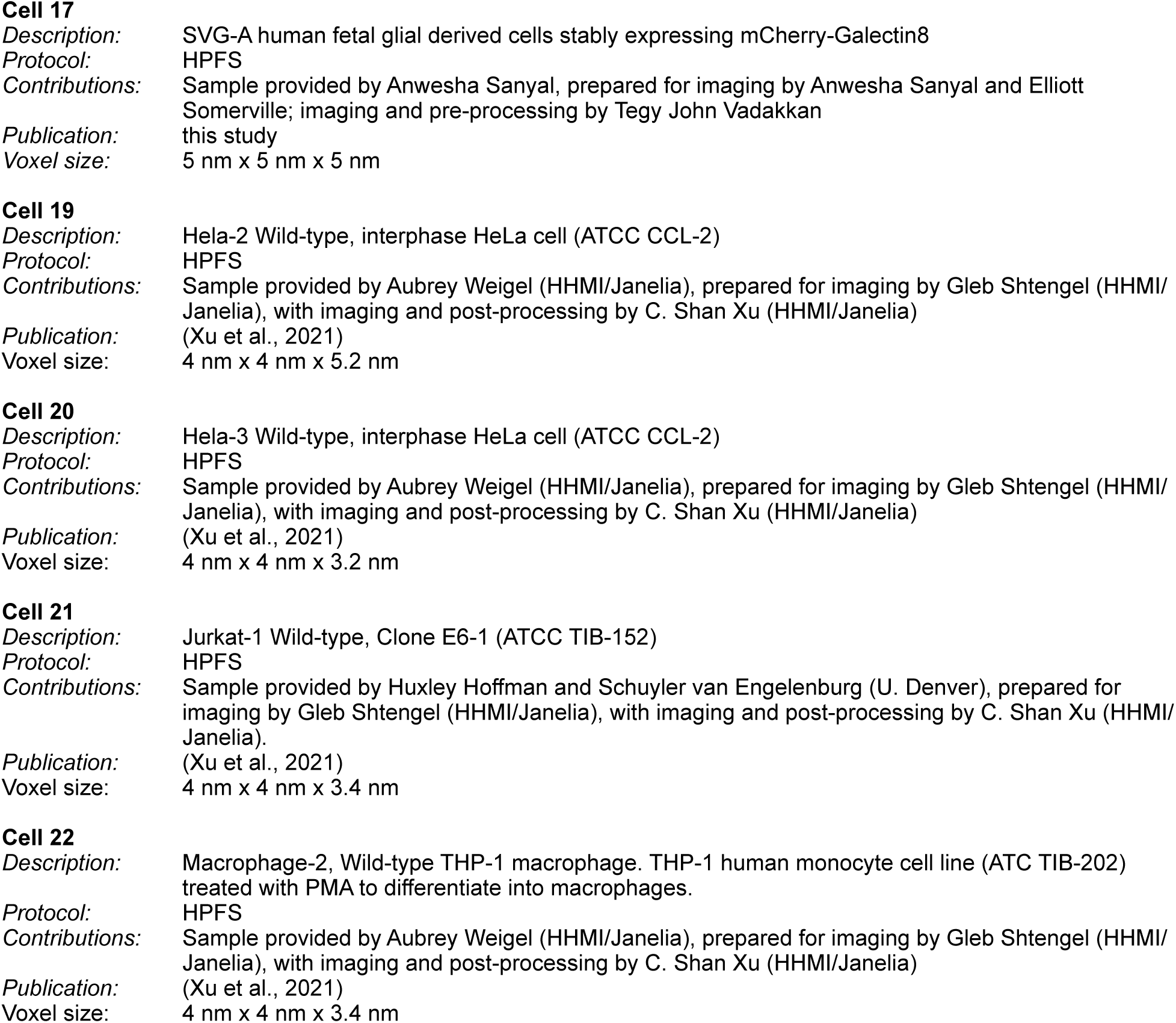
Cells used in this study

**Table S2.**
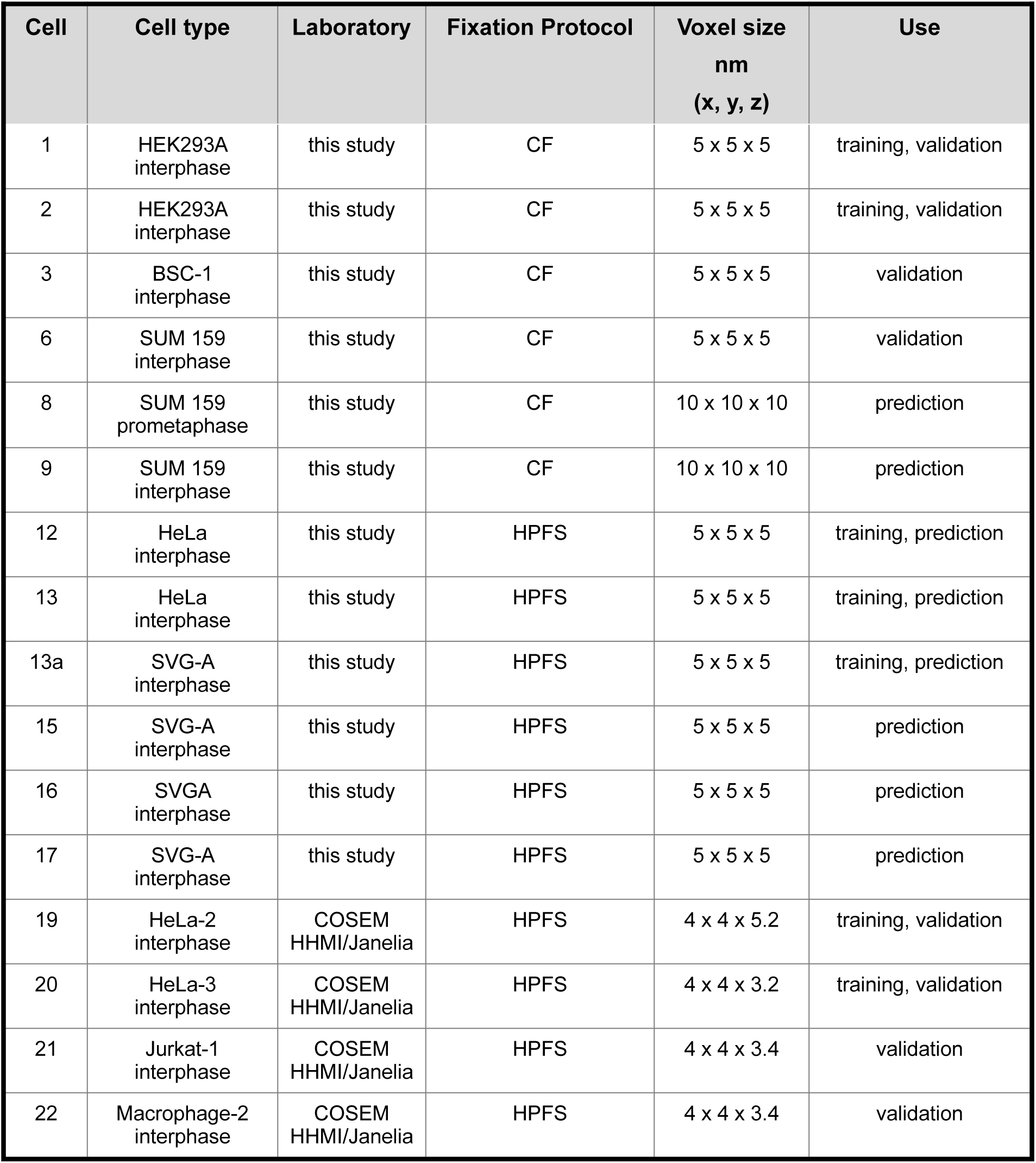
Size of hold out volumes containing ground truth annotations and their use for model training, validation or prediction. Description for each sample includes cell type, stage during cell cycle, fixation protocol (CF or HPFS), FIB-SEM resolution and use of the ground truth annotations for model training, validation or prediction.

**Table S3.**
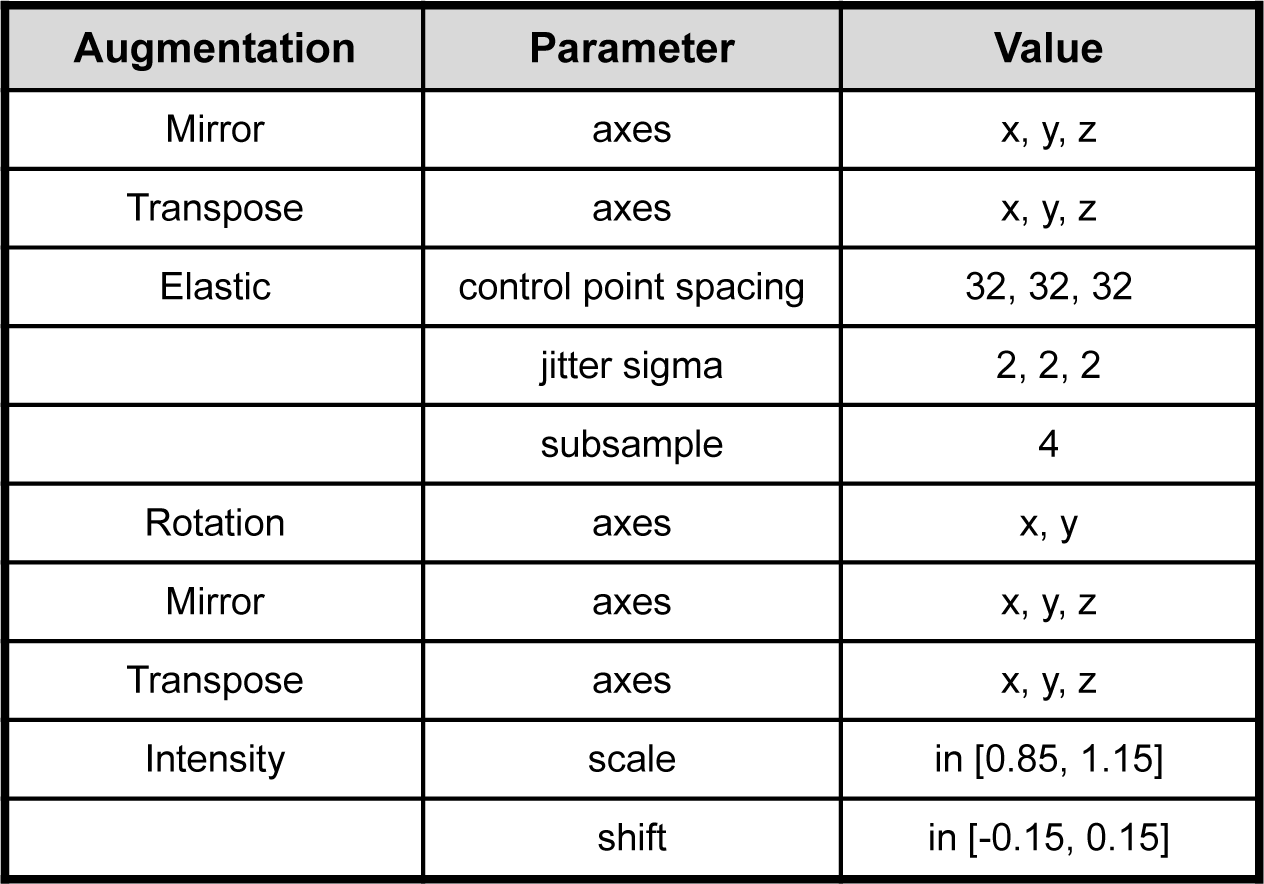
Types of data augmentation used in this study. Type of augmentation and parameters to transform the ground truth annotations used during model training. Their detailed description is found in GUNPOWDER (http://funkey.science/gunpowder/ api.html#augmentation-nodes). Since the rotation operation was performed only in two dimensions, it was necessary to perform the mirror and transpose operations twice in order to obtain all possible 3D orientations of a training block.

**Table S4.**
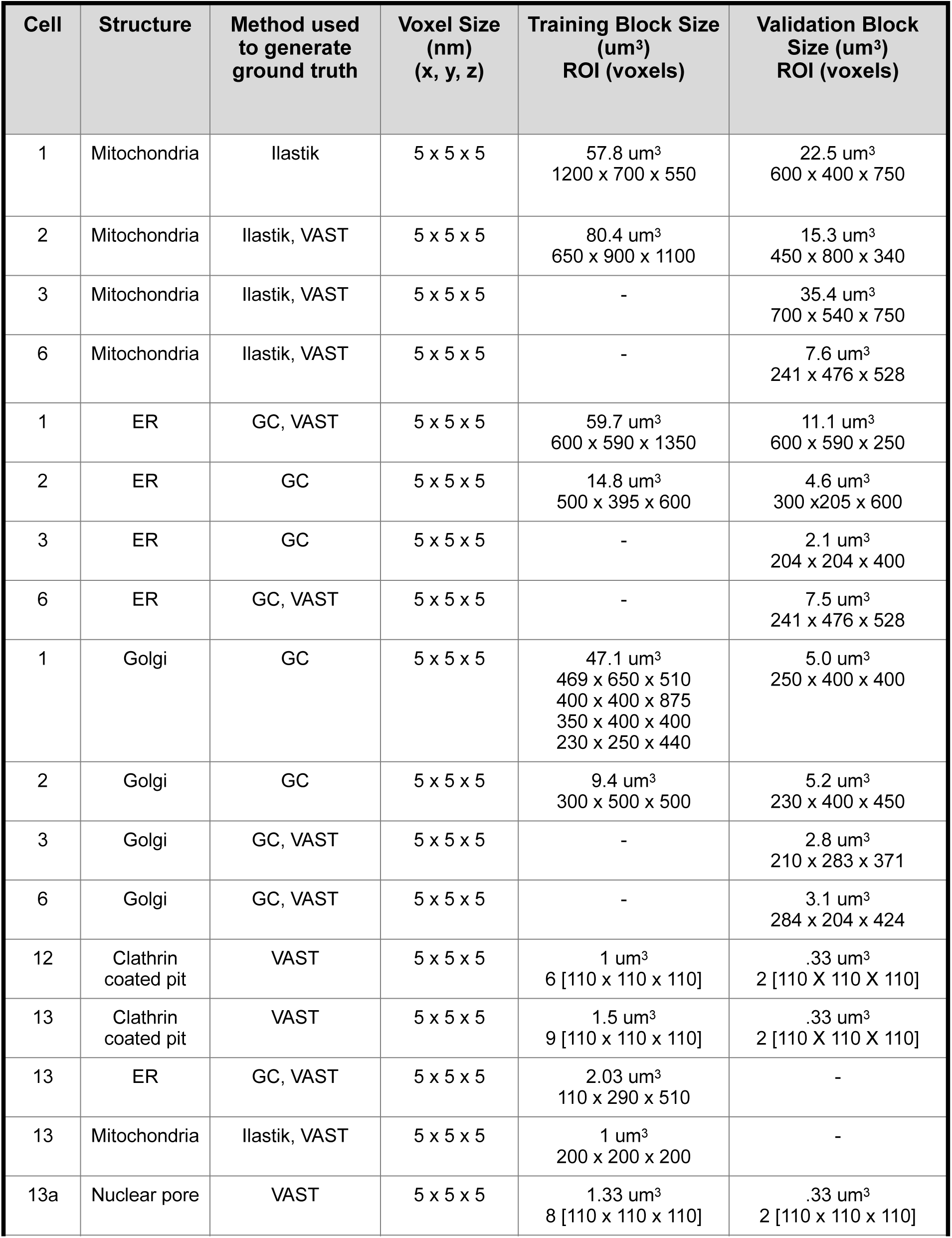
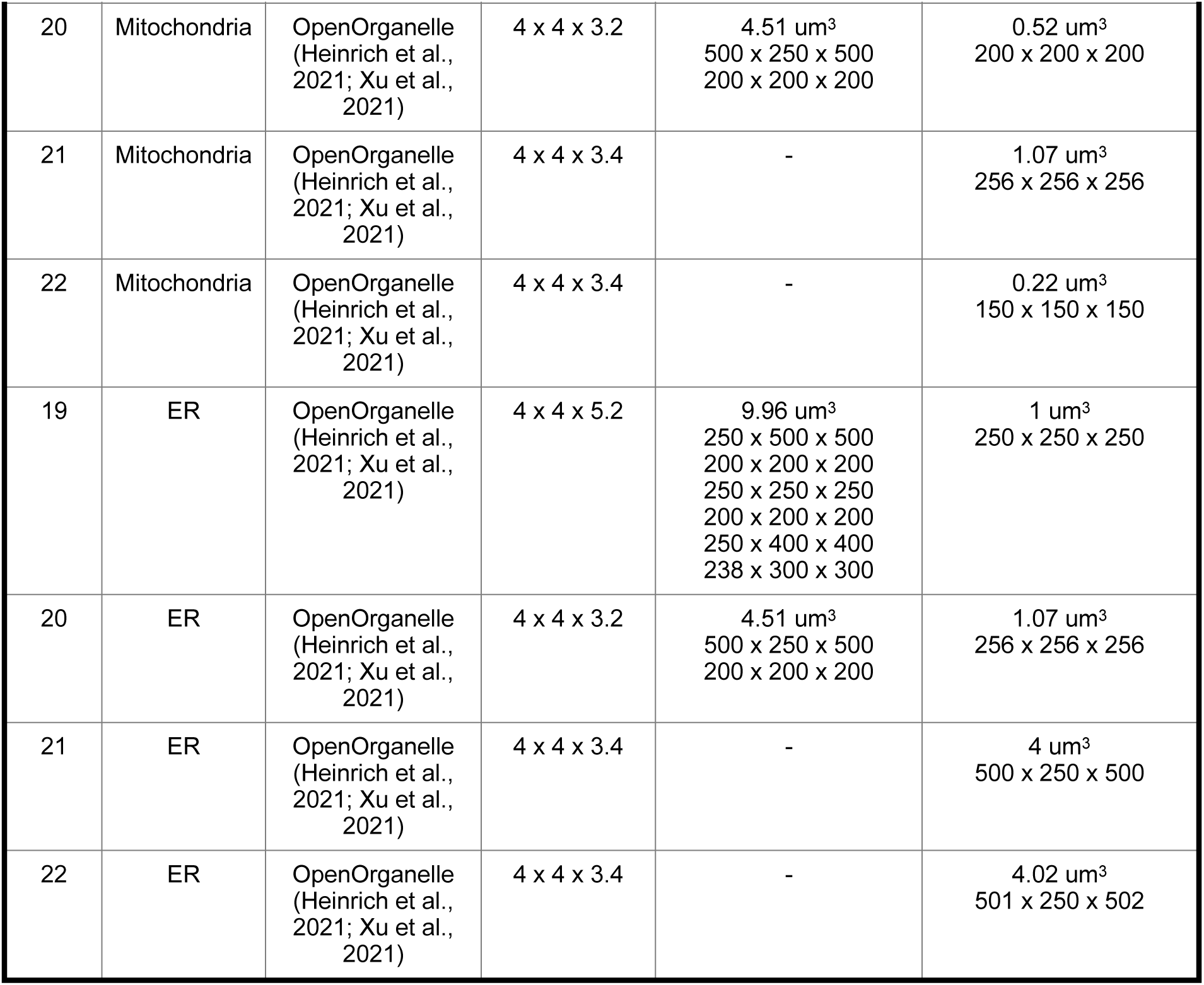
Procedures used to generate ground truth annotations. Description of cells, procedures used to generate ground truth annotations (see methods for details), resolution of FIB-SEM data and size of hold out volumes used for training and validation.

**Table S5.**
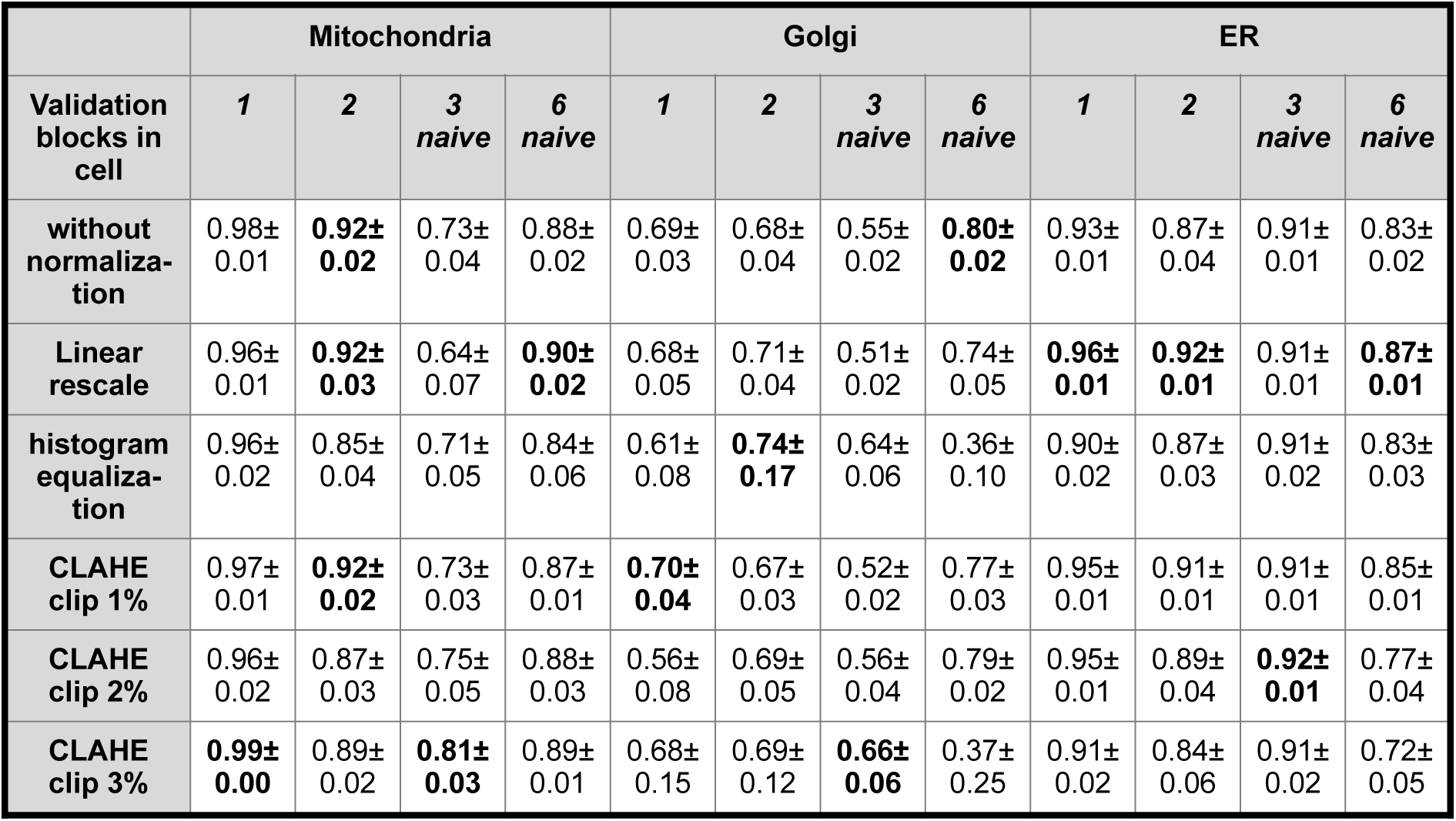
Effect of CLAHE on prediction performance. F1 prediction scores using the indicated cells were obtained with models trained with combined FIB-SEM data from cells 1 and 2 subjected to the indicated types of signal normalization. F1 score for each organelle corresponds to the average +/-SD from 20 consecutive predictions obtained every 1000 iterations initiated after about 150,000 training iterations. Best results are highlighted in bold. Inspection of the data shows no consistent improvement in the F1 predictions scores upon signal normalization including CLAHE.

**Table S6.**
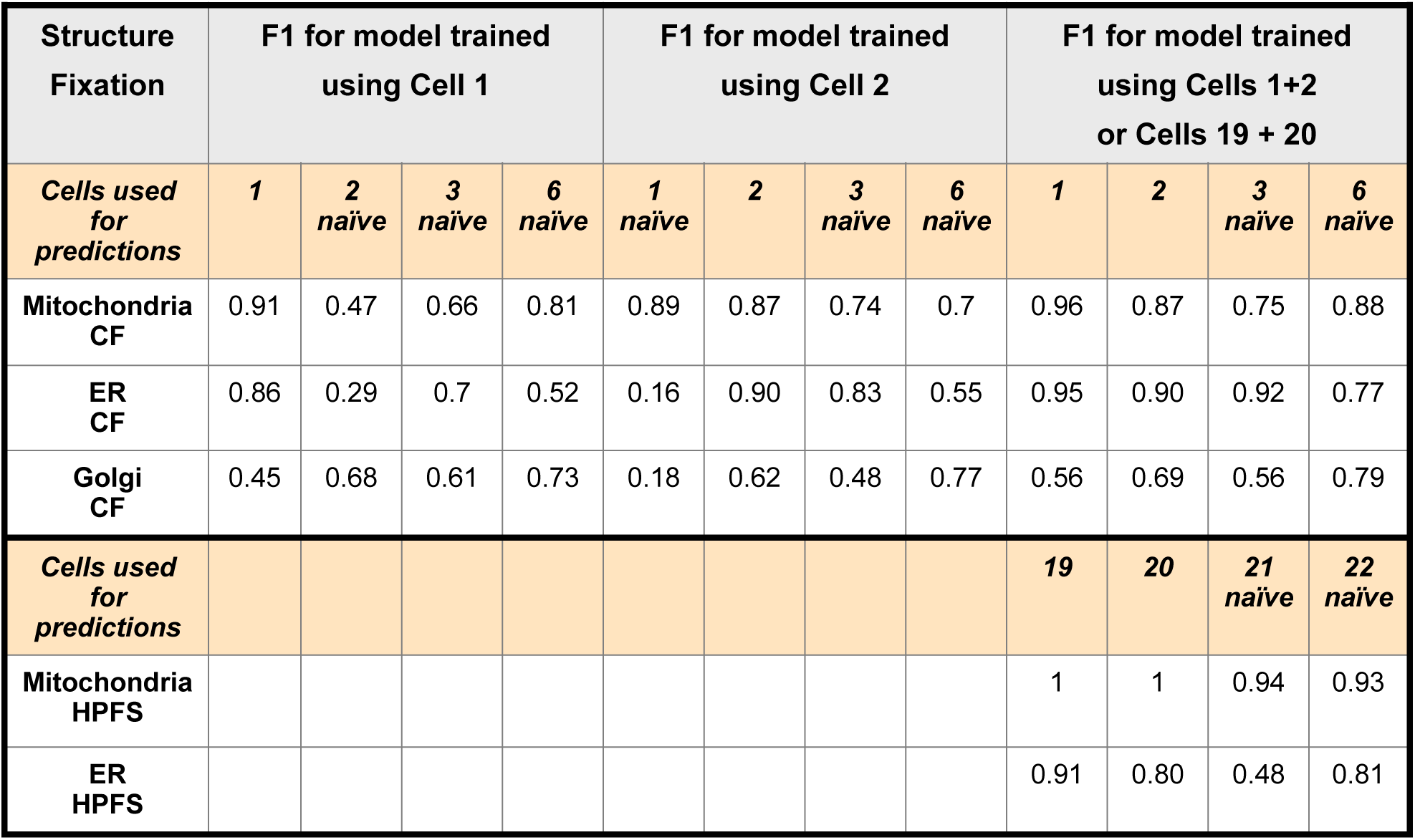
Comparative examples of predictive performance by models trained with data from one or two cells. A neural network was trained using ground truth annotations from the indicated cells, alone or in combination, and the resulting models then used to predict on images of the listed individual cells. The data show F1 prediction scores using validation datasets not employed for training.

**Table S7.**
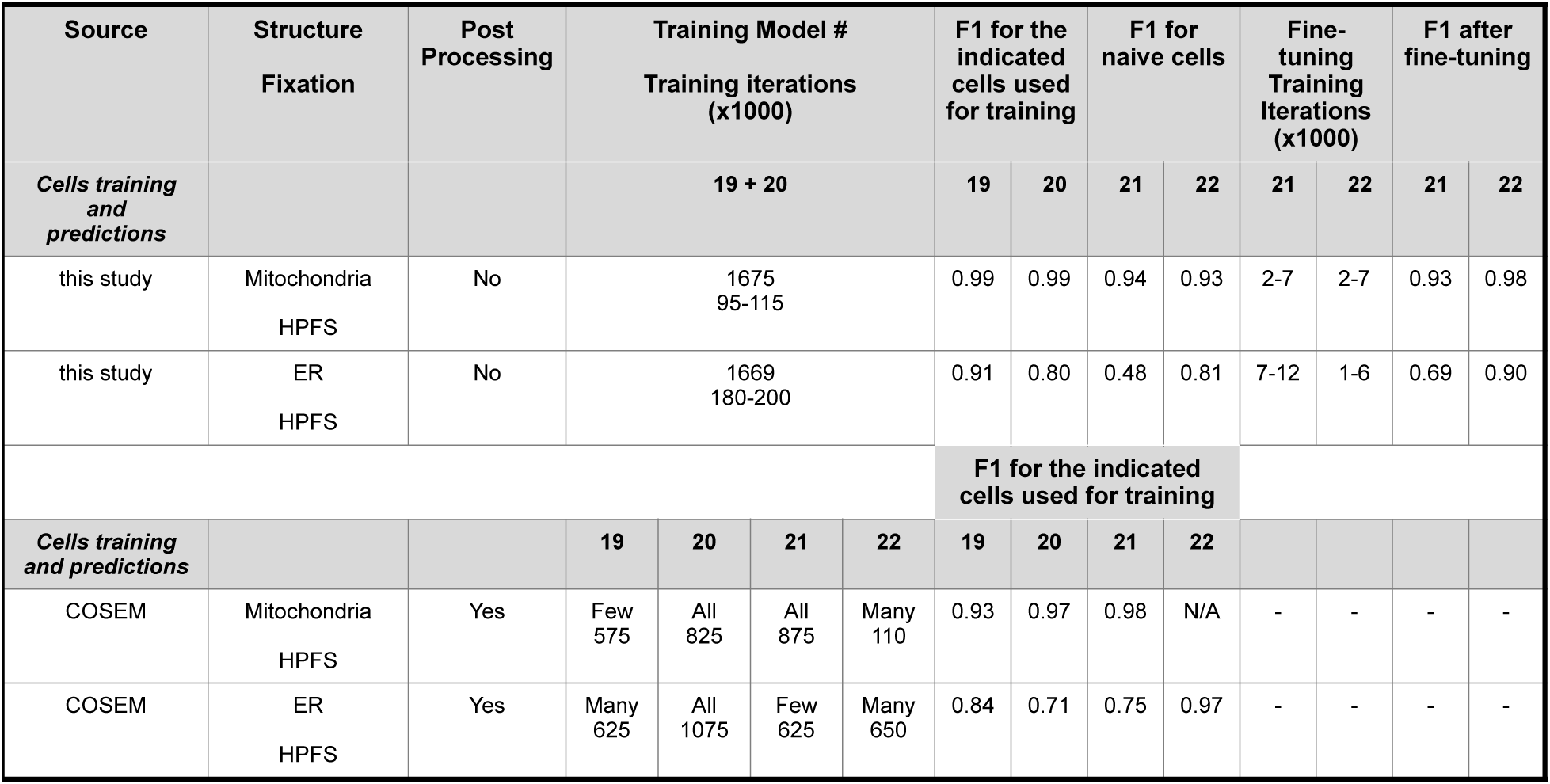
Comparison of model performance using the ASEM (this study) and COSEM pipelines. The neural networks used in this study or by the COSEM project (Heinrich et al., 2021) were trained to predict mitochondria and ER using ground truth annotations from the indicated cells, alone (COSEM project) or in combination (this study). The data show F1 prediction scores using validation datasets not employed for training. In this study we only used data from one organelle at a time to train a neural network and report the results for the defined number of training iterations chosen when the performance of the model reached stability; Reported F1 scores are the average of 20 consecutive values obtained every 1,000 training iterations determined after the indicated training iteration. The COSEM project simultaneously used data from more than one organelle to train the neural network according to the details described in (Heinrich et al., 2021). The F1 scores for the COSEM Project reported in Tables 1 & 2 (Heinrich et al., 2021) correspond to their best results by different trained networks and training iterations using ground annotations for ‘few’, ‘many’ or ‘all’ organelles including mitochondria and ER present within the hold-out block.

**Table S8.**
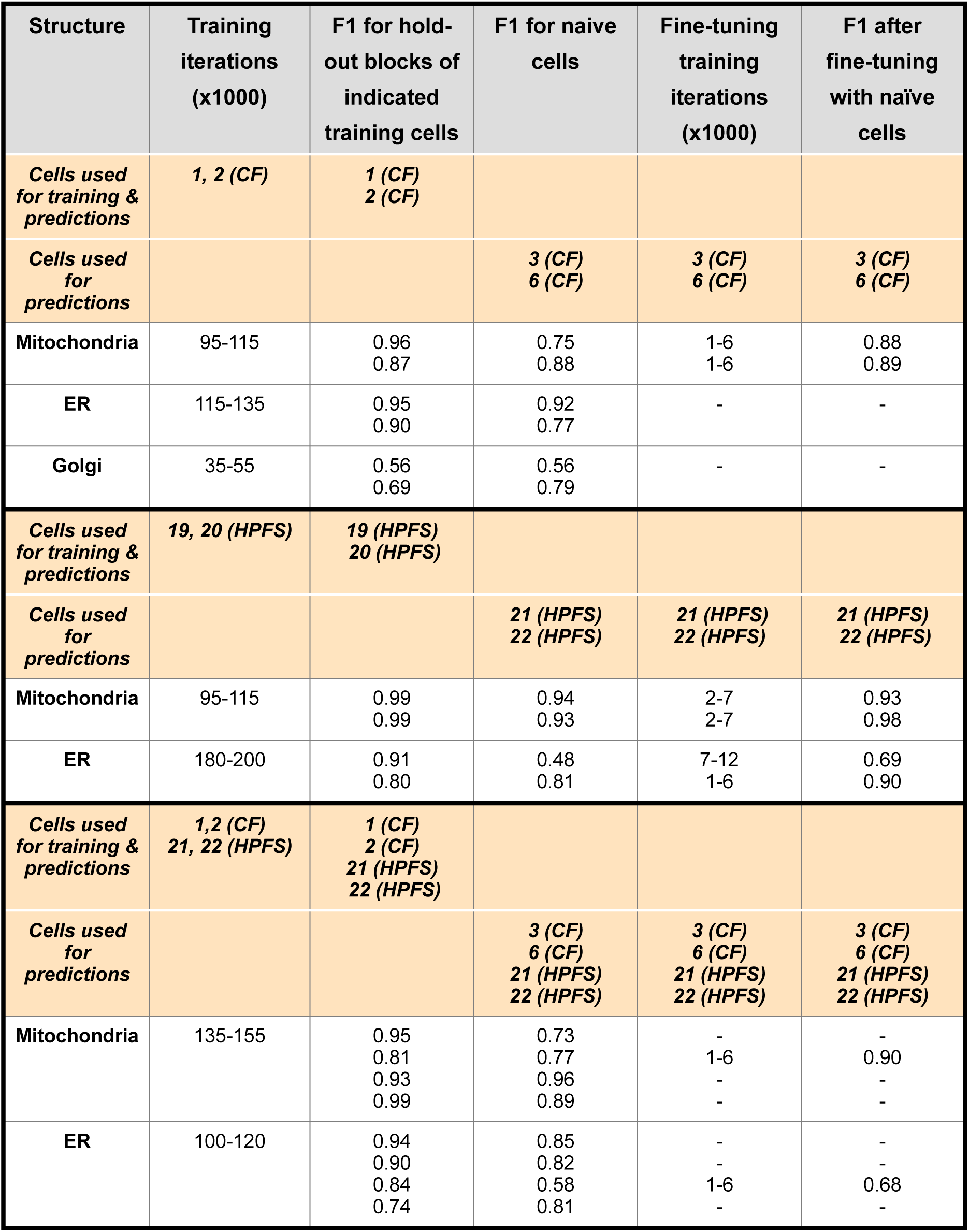
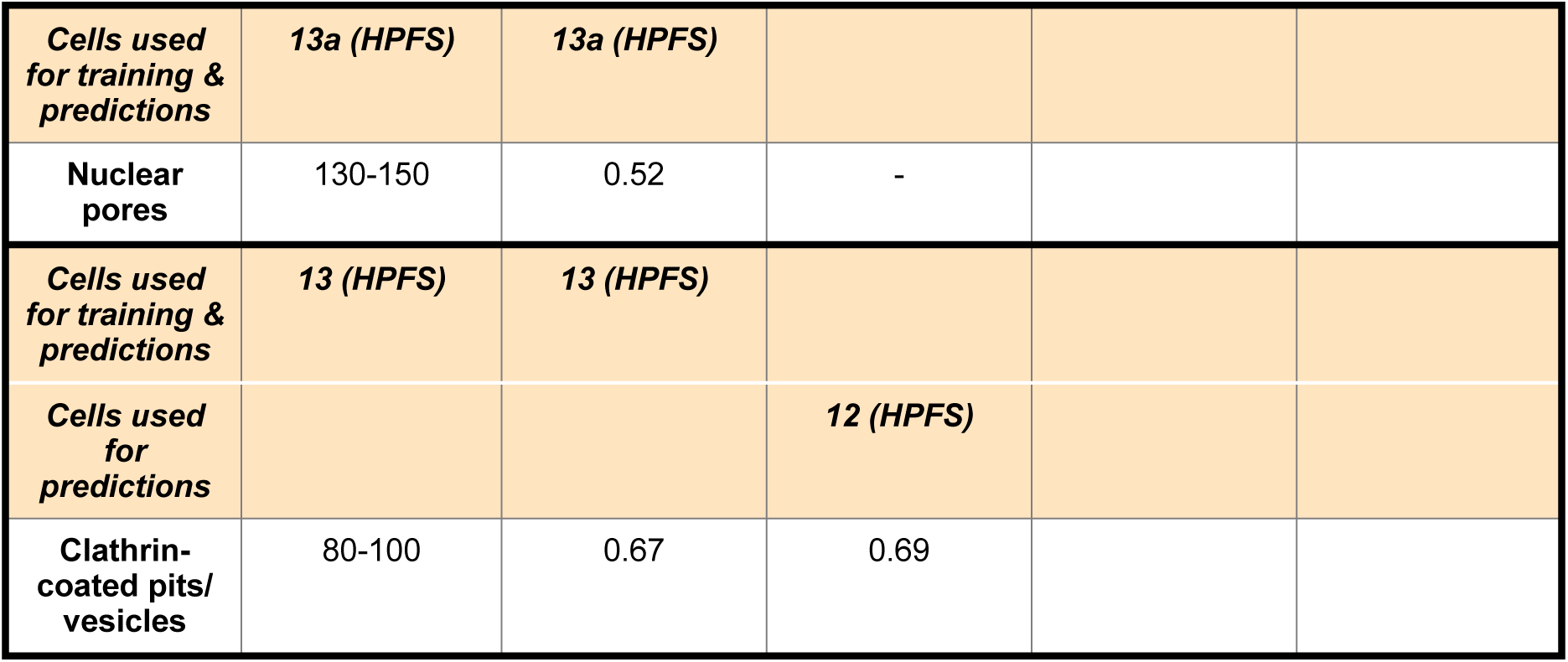
Comparative examples of predictive performance by models trained with data from cells prepared with the same or different fixation protocols. The neural network was trained using ground truth annotations from the indicated cells prepared with different fixation protocols, alone or in combination, and the resulting models then used to predict on images of the individual cells listed in the table. The data show F1 prediction scores using validation datasets not employed for training.

**Table S9.**
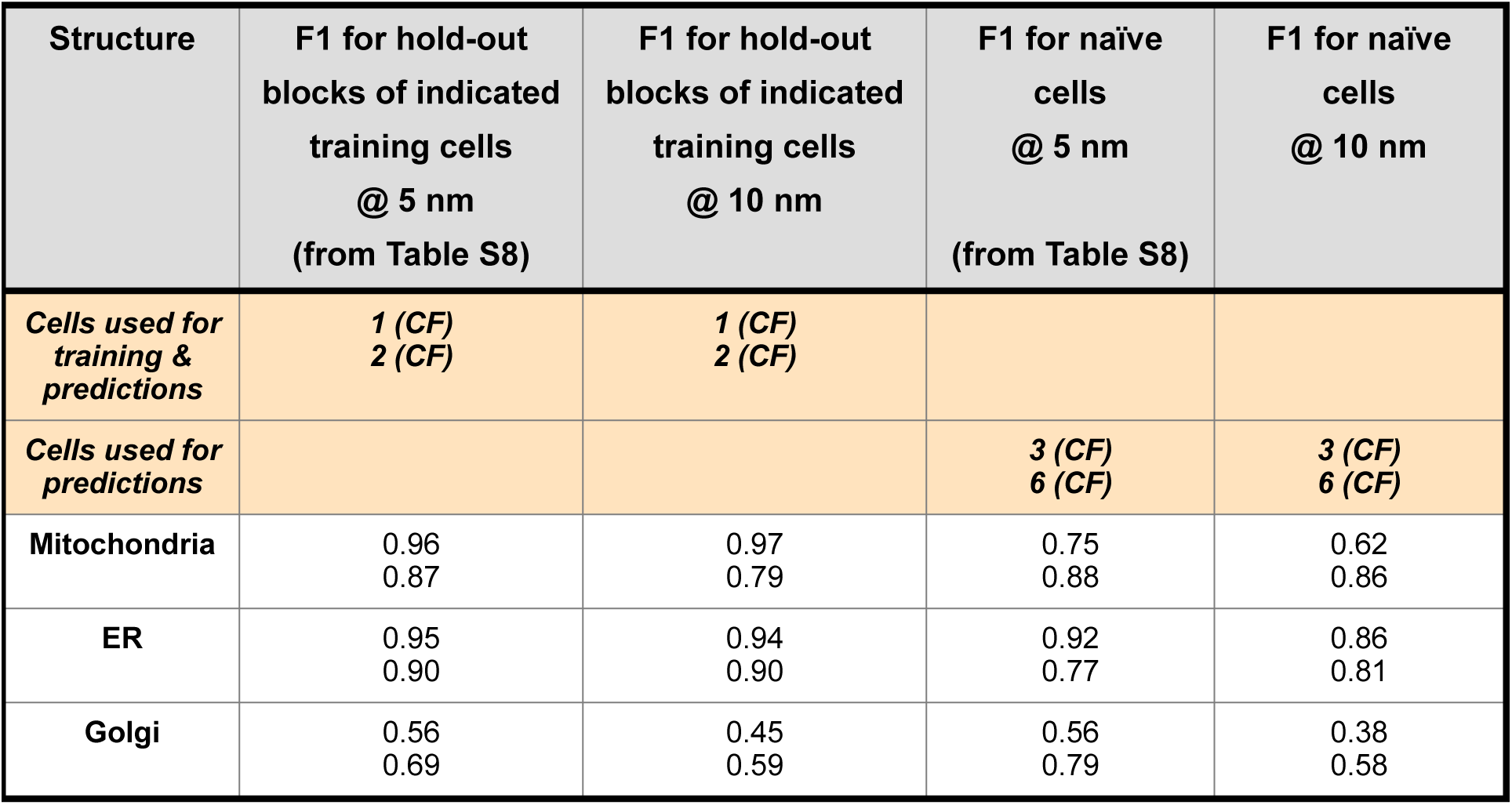
Effect of resolution on the predictive performance. The neural network was trained using ground truth annotations for the organelles from the indicated cells prepared by chemical fixation imaged at 5 nm (Data from Table S8). The resulting models were used to predict on the individual cells listed in the table. The same ground truth annotations were also isotropically downsampled to 10 nm, to then train new models used to predict on images of the same cells, but downsampled to 10 nm. The data show F1 prediction scores using validation datasets not employed for training.

**Table S10.**
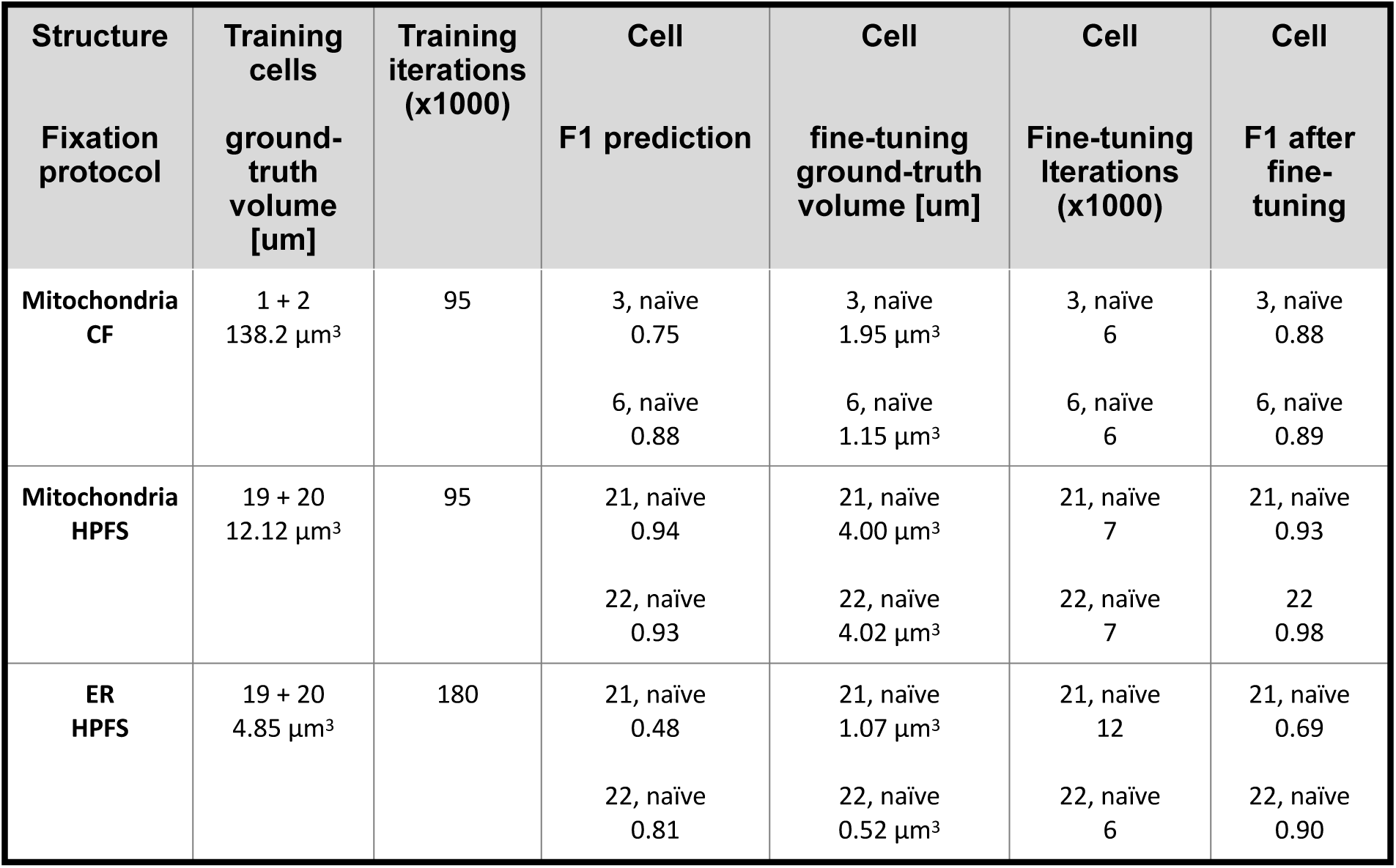
Summary of experiments to test the effect of fine-tuning. Description of models, cells, volumes containing ground truth annotations employed for model training and validation.

## Notes

### Competing Interest Statement

The authors have declared no competing interest.

#### Summary of Updates

Expand Material and methods Perform further analysis

https://github.com/kirchhausenlab/incasem

https://open.quiltdata.com/b/asem-project

## References

Achanta, R., Shaji, A., Smith, K., Lucchi, A., Fua, P., and Süsstrunk, S. (2012). SLIC Superpixels Compared to State-of-the-Art Superpixel Methods. Ieee T Pattern Anal 34, 2274– 2282.

Akisaka, T., Yoshida, H., Suzuki, R., and Takama, K. (2008). Adhesion structures and their cytoskeleton-membrane interactions at podosomes of osteoclasts in culture. Cell Tissue Res 331, 625–641.

Akkiraju, N., Edelsbrunner, H., Facello, M., Fu, F., Mucke, E., and Varella, C. Alpha shapes: definition and software. Proc. Internat. Comput. Geom. Software Workshop.

Berg, S., Kutra, D., Kroeger, T., Straehle, C.N., Kausler, B.X., Haubold, C., Schiegg, M., Ales, J., Beier, T., Rudy, M., et al. (2019). ilastik: interactive machine learning for (bio)image analysis. Nature Methods 16, 1226–1232.

Berger, D.R., Seung, H.S., and Lichtman, J.W. (2018). VAST (Volume Annotation and Segmentation Tool): Efficient Manual and Semi-Automatic Labeling of Large 3D Image Stacks. Front Neural Circuit 12, 88.

Boykov, Y., and Kolmogorov, V. (2004). An Experimental Comparison of Min-Cut/Max-Flow Algorithms for Energy Minimization in Vision. Ieee T Pattern Anal 26, 1124–1137.

Boykov, Y., Veksler, O., and Zabih, R. (2001). Fast approximate energy minimization via graph cuts. Ieee T Pattern Anal 23, 1222–1239.

Buhmann, J., Sheridan, A., Malin-Mayor, C., Schlegel, P., Gerhard, S., Kazimiers, T., Krause, R., Nguyen, T.M., Heinrich, L., Lee, W.-C.A., et al. (2021). Automatic detection of synaptic partners in a whole-brain Drosophila electron microscopy data set. Nat Methods 18, 771–774.

Chen, B.-C., Legant, W.R., Wang, K., Shao, L., Milkie, D.E., Davidson, M.W., Janetopoulos, C., Wu, X.S., Hammer, J.A., Liu, Z., et al. (2014). Lattice light-sheet microscopy: imaging molecules to embryos at high spatiotemporal resolution. Science 346, 1257998.

Chou, Y.-Y., Upadhyayula, S., Houser, J., He, K., Skillern, W., Scanavachi, G., Dang, S., Sanyal, A., Ohashi, K.G., Caprio, G.D., et al. (2021). Inherited nuclear pore substructures template post-mitotic pore assembly. Developmental Cell.

Çiçek, Ö., Abdulkadir, A., Lienkamp, S.S., Brox, T., and Ronneberger, O. (2016). Medical Image Computing and Computer-Assisted Intervention – MICCAI 2016, 19th International Conference, Athens, Greece, October 17-21, 2016, Proceedings, Part II. Lect Notes Comput Sc 424–432.

Ehrlich, M., Boll, W., Oijen, A. van, Hariharan, R., Chandran, K., Nibert, M.L., and Kirchhausen, T. (2004). Endocytosis by random initiation and stabilization of clathrin-coated pits. Cell 118, 591–605.

Gao, R., Asano, S.M., Upadhyayula, S., Pisarev, I., Milkie, D.E., Liu, T.-L., Singh, V., Graves, A., Huynh, G.H., Zhao, Y., et al. (2019). Cortical column and whole-brain imaging with molecular contrast and nanoscale resolution. Science (New York, N.Y.) 363, eaau8302.

Grove, J., Metcalf, D.J., Knight, A.E., Wavre-Shapton, S.T., Sun, T., Protonotarios, E.D., Griffin, L.D., Lippincott-Schwartz, J., and Marsh, M. (2014). Flat clathrin lattices: stable features of the plasma membrane. Molecular Biology of the Cell 25, 3581–3594.

Guay, M.D., Emam, Z.A.S., Anderson, A.B., Aronova, M.A., Pokrovskaya, I.D., Storrie, B., and Leapman, R.D. (2021). Dense cellular segmentation for EM using 2D–3D neural network ensembles. Sci Rep-Uk 11, 2561.

Haberl, M.G., Churas, C., Tindall, L., Boassa, D., Phan, S., Bushong, E.A., Madany, M., Akay, R., Deerinck, T.J., Peltier, S.T., et al. (2018). CDeep3M-Plug-and-Play cloud-based deep learning for image segmentation. Nature Methods 15, 677–680.

Haberl, M.G., Wong, W., Penticoff, S., Je, J., Madany, M., Borchardt, A., Boassa, D., Peltier, S.T., and Ellisman, M.H. (2020). CDeep3M-Preview: Online segmentation using the deep neural network model zoo. BioRxiv 16, 1233.

Heinrich, L., Bennett, D., Ackerman, D., Park, W., Bogovic, J., Eckstein, N., Petruncio, A., Clements, J., Pang, S., Xu, C.S., et al. (2021). Whole-cell organelle segmentation in volume electron microscopy. Nature 1–6.

Heuser, J. (1980). Three-dimensional visualization of coated vesicle formation in fibroblasts. J Cell Biol 84, 560–583.

Hoffman, D.P., Shtengel, G., Xu, C.S., Campbell, K.R., Freeman, M., Wang, L., Milkie, D.E., Pasolli, H.A., Iyer, N., Bogovic, J.A., et al. (2020). Correlative three-dimensional super-resolution and block-face electron microscopy of whole vitreously frozen cells. Science (New York, N.Y.) 367, eaaz5357.

Kingma, D.P., and Ba, J. (2014). Adam: A Method for Stochastic Optimization. Arxiv.

Kirchhausen, T. (1993). Coated pits and coated vesicles - sorting it all out. Current Opinion in Structural Biology 3, 182–188.

Kirchhausen, T. (2000). Clathrin. Annual Review of Biochemistry 69, 699–727.

Kirchhausen, T. (2009). Imaging endocytic clathrin structures in living cells. Trends in Cell Biology 19, 596–605.

Kirchhausen, T., Owen, D., and Harrison, S.C. (2014). Molecular structure, function, and dynamics of clathrin-mediated membrane traffic. Cold Spring Harbor Perspectives in Biology 6, a016725.

Knott, G., Marchman, H., Wall, D., and Lich, B. (2008). Serial Section Scanning Electron Microscopy of Adult Brain Tissue Using Focused Ion Beam Milling. J Neurosci 28, 2959–2964.

Kolmogorov, V., and Zabin, R. (2004). What energy functions can be minimized via graph cuts? Ieee T Pattern Anal 26, 147–159.

Liu, J., Li, L., Yang, Y., Hong, B., Chen, X., Xie, Q., and Han, H. (2020). Automatic Reconstruction of Mitochondria and Endoplasmic Reticulum in Electron Microscopy Volumes by Deep Learning. Front Neurosci-Switz 14, 599.

Liu, T.-L., Upadhyayula, S., Milkie, D.E., Singh, V., Wang, K., Swinburne, I.A., Mosaliganti, K.R., Collins, Z.M., Hiscock, T.W., Shea, J., et al. (2018). Observing the cell in its native state: Imaging subcellular dynamics in multicellular organisms. Science (New York, N.Y.) 360.

Lucchi, A., Smith, K., Achanta, R., Knott, G., and Fua, P. (2012). Supervoxel-Based Segmentation of Mitochondria in EM Image Stacks with Learned Shape Features. Ieee T Med Imaging 31, 474–486.

Maupin, P., and Pollard, T.D. (1983). Improved preservation and staining of HeLa cell actin filaments, clathrin-coated membranes, and other cytoplasmic structures by tannic acid-glutaraldehyde-saponin fixation. The Journal of Cell Biology 96, 51–62.

Mekuč, M.Ž., Bohak, C., Hudoklin, S., Kim, B.H., Romih, R., Kim, M.Y., and Marolt, M. (2020). Automatic segmentation of mitochondria and endolysosomes in volumetric electron microscopy data. Comput Biol Med 119, 103693.

Mekuč, M.Ž., Bohak, C., Boneš, E., Hudoklin, S., Romih, R., and Marolt, M. (2022). Automatic segmentation and reconstruction of intracellular compartments in volumetric electron microscopy data. Comput Meth Prog Bio 223, 106959.

Müller, A., Schmidt, D., Xu, C.S., Pang, S., D’Costa, J.V., Kretschmar, S., Münster, C., Kurth, T., Jug, F., Weigert, M., et al. (2020). 3D FIB-SEM reconstruction of microtubule–organelle interaction in whole primary mouse β cells. J Cell Biology 220, e202010039.

Otsu, N. (1979). A Threshold Selection Method from Gray-Level Histograms. Ieee Transactions Syst Man Cybern 9, 62–66.

Paszke, A., Gross, S., Massa, F., Lerer, A., Bradbury, J., Chanan, G., Killeen, T., Lin, Z., Gimelshein, N., Antiga, L., et al. (2019). PyTorch: An Imperative Style, High-Performance Deep Learning Library. Arxiv.

Pizer, S.M., Amburn, E.P., Austin, J.D., Cromartie, R., Geselowitz, A., Greer, T., Romeny, B. ter H., Zimmerman, J.B., and Zuiderveld, K. (1987). Adaptive histogram equalization and its variations. Comput Vis Graph Image Process 39, 355–368.

Saffarian, S., Cocucci, E., and Kirchhausen, T. (2009). Distinct dynamics of endocytic clathrin-coated pits and coated plaques. PLoS Biology 7, e1000191.

Schroeder, A.B., Dobson, E.T.A., Rueden, C.T., Tomancak, P., Jug, F., and Eliceiri, K.W. (2021). The ImageJ ecosystem: Open-source software for image visualization, processing, and analysis. Protein Sci 30, 234–249.

Schuller, A.P., Wojtynek, M., Mankus, D., Tatli, M., Kronenberg-Tenga, R., Regmi, S.G., Dip, P.V., Lytton-Jean, A.K.R., Brignole, E.J., Dasso, M., et al. (2021). The cellular environment shapes the nuclear pore complex architecture. Nature 1–5.

Shorten, C., and Khoshgoftaar, T.M. (2019). A survey on Image Data Augmentation for Deep Learning. J Big Data 6, 60.

Signoret, N., Hewlett, L., Wavre, S., Pelchen-Matthews, A., Oppermann, M., and Marsh, M. (2005). Agonist-induced Endocytosis of CC Chemokine Receptor 5 Is Clathrin Dependent. Mol Biol Cell 16, 902–917.

Studer, D., Humbel, B.M., and Chiquet, M. (2008). Electron microscopy of high pressure frozen samples: bridging the gap between cellular ultrastructure and atomic resolution. Histochem Cell Biol 130, 877–889.

Virtanen, P., Gommers, R., Oliphant, T.E., Haberland, M., Reddy, T., Cournapeau, D., Burovski, E., Peterson, P., Weckesser, W., Bright, J., et al. (2020). SciPy 1.0: fundamental algorithms for scientific computing in Python. Nat Methods 17, 261–272.

Walt, S. van der, Schönberger, J.L., Nunez-Iglesias, J., Boulogne, F., Warner, J.D., Yager, N., Gouillart, E., Yu, T., and contributors, scikit-image (2014). scikit-image: image processing in Python. Peerj 2, e453.

Wei, D., Lin, Z., Franco-Barranco, D., Wendt, N., Liu, X., Yin, W., Huang, X., Gupta, A., Jang, W.-D., Wang, X., et al. (2020). Medical Image Computing and Computer Assisted Intervention – MICCAI 2020, 23rd International Conference, Lima, Peru, October 4–8, 2020, Proceedings, Part V. Lect Notes Comput Sc 12265, 66–76.

Weiss, K., Khoshgoftaar, T.M., and Wang, D. (2016). A survey of transfer learning. J Big Data 3, 9.

Willy, N.M., Ferguson, J.P., Akatay, A., Huber, S., Djakbarova, U., Silahli, S., Cakez, C., Hasan, F., Chang, H.C., Travesset, A., et al. (2021). De novo endocytic clathrin coats develop curvature at early stages of their formation. Dev Cell 56, 3146–3159.e5.

Xu, C.S., Hayworth, K.J., Lu, Z., Grob, P., Hassan, A.M., García-Cerdán, J.G., Niyogi, K.K., Nogales, E., Weinberg, R.J., and Hess, H.F. (2017). Enhanced FIB-SEM systems for large-volume 3D imaging. ELife 6, 185.

Xu, C.S., Pang, S., Shtengel, G., Müller, A., Ritter, A.T., Hoffman, H.K., Takemura, S., Lu, Z., Pasolli, H.A., Iyer, N., et al. (2021). An open-access volume electron microscopy atlas of whole cells and tissues. Nature 599, 147–151.

Zeng, T., Murphy, R., Wu, B., and Ji, S. (2017). DeepEM3D: approaching human-level performance on 3D anisotropic EM image segmentation. Bioinformatics (Oxford, England) 33, 2555–2562.

Zimmerli, C.E., Allegretti, M., Rantos, V., Goetz, S.K., Obarska-Kosinska, A., Zagoriy, I., Halavatyi, A., Hummer, G., Mahamid, J., Kosinski, J., et al. (2021). Nuclear pores dilate and constrict in cellulo. Sci New York N Y 374, eabd9776.

Zuiderveld, K. (1994). Graphics Gems. Viii Image Process 474–485.

## References

Sheridan, A., Nguyen, T., Deb, D., Lee, W.-C.A., Saalfeld, S., Turaga, S., Manor, U., and Funke, J. (2022). Local Shape Descriptors for Neuron Segmentation. Biorxiv 2021.01.18.427039.

## Supplementary References

Chou, Y.-Y., Krupp, A., Kaynor, C., Gaudin, R., Ma, M., Cahir-McFarland, E., and Kirchhausen, T. (2016a). Inhibition of JCPyV infection mediated by targeted viral genome editing using CRISPR/Cas9. Scientific Reports 6, 36921.

Chou, Y.-Y., Cuevas, C., Carocci, M., Stubbs, S.H., Ma, M., Cureton, D.K., Chao, L., Evesson, F., He, K., Yang, P.L., et al. (2016b). Identification and characterization of a novel broad spectrum virus entry inhibitor. J Virol 90, 4494– 4510.

