## Supplementary material for "Deep neural network automated segmentation of cellular structures in volume electron microscopy": Tables and Supporting Figures

### Table S1. Cells used in this study

**Cell 1, 1a**
*Description*: HEK293A human epithelial derived cell stably expressing eGFP-connexin43 (ATCC, crl-1573)
*Protocol:* Chemical fixation
*Contributions:* Sample provided by Teresa Rodrigues and Henrique Girão (Universidade de Coimbra), prepared for imaging by Giovanni de Nola and Teresa Rodrigues; imaging and pre-processing by Tegy John Vadakkan; post-processing by Ben Gallusser
*Publication:* this study
*Voxel size:* 5 nm x 5 nm x 5 nm

**Cell 2**
*Description*: HEK293A human epithelial derived cell stably expressing eGFP-connexin43 (ATCC, crl-1573)
*Protocol:* Chemical fixation
*Contributions:* Sample provided by Teresa Rodrigues and Henrique Girão (Universidade de Coimbra), prepared for imaging by Giovanni de Nola and Teresa Rodrigues; imaging and pre-processing by Tegy John Vadakkan; post-processing by Ben Gallusser
*Publication:* this study
*Voxel size:* 5 nm x 5 nm x 5 nm

**Cell 3**
Description: BSC-1 African green monkey kidney epithelial derived cells stably expressing Lamp1-eGFP and mCherry-Galectin3
*Protocol:* Chemical fixation
*Contributions:* Sample prepared by Teresa Rodrigues (Universidade de Coimbra), prepared for imaging by Giovanni de Nola and Teresa Rodrigues; imaging and pre-processing by Tegy John Vadakkan; post-processing by Ben Gallusser
*Publication:* this study
*Voxel size:* 5 nm x 5 nm x 5 nm

**Cell 4**
Description: BSC-1 African green monkey kidney epithelial derived cells stably expressing Lamp1-eGFP and mCherry-Galectin3
*Protocol:* Chemical fixation
*Contributions:* Sample prepared by Teresa Rodrigues, prepared for imaging by Giovanni de Nola and Teresa Rodrigues; imaging and pre-processing by Tegy John Vadakkan; post-processing by Ben Gallusser
*Publication:* this study
*Voxel size:* 5 nm x 5 nm x 5 nm

**Cell 5**
*Description:* SVG-A human fetal glial derived cells
*Protocol:* Chemical fixation
*Contributions:* Prepared for imaging by Rasmus Herlo and Max Paget; imaging and pre-processing by Tegy John Vadakkan; post-processing by Ben Gallusser
*Publication:* (Chou et al., 2021)
*Voxel size:* 5 nm x 5 nm x 5 nm

**Cell 6, 6a**
*Description:* interphase SUM159 human breast carcinoma derived cell gene edited to express eGFP-Nup133
*Protocol:* Chemical fixation
*Contributions:* Prepared for imaging by Justin Houser; imaging and pre-processing by Tegy John Vadakkan; post-processing by Ben Gallusser
*Publication:* (Chou et al., 2021)
*Voxel size:* 5 nm x 5 nm x 5 nm

**Cell 7**
Description: mitotic SUM159 human breast carcinoma derived cell gene edited to express eGFP-Nup133
*Protocol:* Chemical fixation
*Contributions:* Prepared for imaging by Justin Houser; imaging and pre-processing by Tegy John Vadakkan; post-processing by Ben Gallusser
*Publication:* (Chou et al., 2021)
*Voxel size:* 5 nm x 5 nm x 5 nm

**Cell 8**

Description: pro-metaphase SUM159 human breast carcinoma derived cell gene edited to express eGFP-Nup133
*Protocol:* Chemical fixation
*Contributions:* Prepared for imaging by Justin Houser; imaging and pre-processing by Justin Houser; post-processing by Ben Gallusser
*Publication:* (Chou et al., 2021)
*Voxel size:* 10 nm x 10 nm x 10 nm

**Cell 9**
*Description:* interphase SUM159 human breast carcinoma derived cell gene edited to express eGFP-Nup133
*Protocol:* Chemical fixation
*Contributions:* Prepared for imaging by Justin Houser; imaging and pre-processing by Justin Houser; post-processing by Ben Gallusser
*Publication:* (Chou et al., 2021)
*Voxel size:* 10 nm x 10 nm x 10 nm

**Cell 11**
*Description:* interphase U2OS human sarcoma derived cell
*Protocol:* HPFS
*Contributions:* Prepared for imaging by Gleb Shtengel (HHMI/Janelia) and C. Shan Xu (HHMI/Janelia); post-processing by Ben Gallusser
*Publication:* this study
*Voxel size:* 8 nm x 8 nm x 8 nm

**Cell 12**
*Description:* HeLa cell
*Protocol:* HPFS
*Contributions:* Sample provided and prepared for imaging by HHMI/Janelia; imaging and pre-processing by Tegy John Vadakkan
*Publication:* this study
*Voxel size:* 5 nm x 5 nm x 5 nm

**Cell 13**
*Description:* HeLa cell
*Protocol:* HPFS
*Contributions:* Sample provided and prepared for imaging by HHMI/Janelia; imaging and pre-processing by Tegy John Vadakkan
*Publication:* this study
*Voxel size:* 5 nm x 5 nm x 5 nm

**Cell 13a**
*Description:* SVG-A human fetal glial derived cells stably expressing mCherry-Galectin8
*Protocol:* HPFS
*Contributions:* Prepared for imaging by Anwesha Sanyal and Elliott Somerville; imaging and pre-processing by Tegy John Vadakkan
*Publication:* this study
*Voxel size:* 5 nm x 5 nm x 5 nm

**Cell 15**
*Description:* SVG-A human fetal glial derived cells stably expressing mCherry-Galectin8
*Protocol:* HPFS
*Contributions:* Sample provided by Anwesha Sanyal, prepared for imaging by Anwesha Sanyal and Elliott Somerville; imaging and pre-processing by Tegy John Vadakkan
*Publication:* this study
*Voxel size:* 5 nm x 5 nm x 5 nm

**Cell 16**
*Description:* SVG-A human fetal glial derived cells stably expressing mCherry-Galectin8
*Protocol:* HPFS
*Contributions:* Sample provided by Anwesha Sanyal, prepared for imaging by Anwesha Sanyal and Elliott Somerville; imaging and pre-processing by Tegy John Vadakkan
*Publication:* this study
*Voxel size:* 5 nm x 5 nm x 5 nm

**Cell 17**
*Description:* SVG-A human fetal glial derived cells stably expressing mCherry-Galectin8
*Protocol:* HPFS
*Contributions:* Sample provided by Anwesha Sanyal, prepared for imaging by Anwesha Sanyal and Elliott Somerville; imaging and pre-processing by Tegy John Vadakkan
*Publication:* this study
*Voxel size:* 5 nm x 5 nm x 5 nm

**Cell 19**
*Description:* Hela-2 Wild-type, interphase HeLa cell (ATCC CCL-2)
*Protocol:* HPFS
*Contributions:* Sample provided by Aubrey Weigel (HHMI/Janelia), prepared for imaging by Gleb Shtengel (HHMI/Janelia), with imaging and post-processing by C. Shan Xu (HHMI/Janelia)
*Publication:* (Xu et al., 2021)
Voxel size: 4 nm x 4 nm x 5.2 nm

**Cell 20**
*Description:* Hela-3 Wild-type, interphase HeLa cell (ATCC CCL-2)
*Protocol:* HPFS
*Contributions:* Sample provided by Aubrey Weigel (HHMI/Janelia), prepared for imaging by Gleb Shtengel (HHMI/Janelia), with imaging and post-processing by C. Shan Xu (HHMI/Janelia)
*Publication:* (Xu et al., 2021)
Voxel size: 4 nm x 4 nm x 3.2 nm

**Cell 21**
*Description:* Jurkat-1 Wild-type, Clone E6-1 (ATCC TIB-152)
*Protocol:* HPFS
*Contributions:* Sample provided by Huxley Hoffman and Schuyler van Engelenburg (U. Denver), prepared for imaging by Gleb Shtengel (HHMI/Janelia), with imaging and post-processing by C. Shan Xu (HHMI/Janelia).
*Publication:* (Xu et al., 2021)
Voxel size: 4 nm x 4 nm x 3.4 nm

**Cell 22**
*Description:* Macrophage-2, Wild-type THP-1 macrophage. THP-1 human monocyte cell line (ATC TIB-202) treated with PMA to differentiate into macrophages.
*Protocol:* HPFS
*Contributions:* Sample provided by Aubrey Weigel (HHMI/Janelia), prepared for imaging by Gleb Shtengel (HHMI/Janelia), with imaging and post-processing by C. Shan Xu (HHMI/Janelia)
*Publication:* (Xu et al., 2021)
Voxel size: 4 nm x 4 nm x 3.4 nm

Description for each sample includes cell type, fixation protocol (CF or HPFS), source and FIB-SEM resolution**.**

#### Supplementary References

Chou, Y.-Y., Krupp, A., Kaynor, C., Gaudin, R., Ma, M., Cahir-McFarland, E., and Kirchhausen, T. (2016a). Inhibition of JCPyV infection mediated by targeted viral genome editing using CRISPR/Cas9. Scientific Reports *6*, 36921.

Chou, Y.-Y., Cuevas, C., Carocci, M., Stubbs, S.H., Ma, M., Cureton, D.K., Chao, L., Evesson, F., He, K., Yang, P.L., et al. (2016b). Identification and characterization of a novel broad spectrum virus entry inhibitor. J Virol *90*, 4494–4510.

Chou, Y.-Y., Upadhyayula, S., Houser, J., He, K., Skillern, W., Scanavachi, G., Dang, S., Sanyal, A., Ohashi, K.G., Caprio, G.D., et al. (2021). Inherited nuclear pore substructures template post-mitotic pore assembly. Developmental Cell.

Ehrlich, M., Boll, W., Oijen, A. van, Hariharan, R., Chandran, K., Nibert, M.L., and Kirchhausen, T. (2004). Endocytosis by random initiation and stabilization of clathrin-coated pits. Cell *118*, 591–605.

Xu, C.S., Pang, S., Shtengel, G., Müller, A., Ritter, A.T., Hoffman, H.K., Takemura, S., Lu, Z., Pasolli, H.A., Iyer, N., et al. (2021). An open-access volume electron microscopy atlas of whole cells and tissues. Nature *599*, 147–151.

**Table S2. Size of hold out volumes containing ground truth annotations and their use for model training, validation or prediction**

| Cell | Cell type | Laboratory | Fixation Protocol | Voxel size nm (x, y, z) | Use |
|---|---|---|---|---|---|
| 1 | HEK293A interphase | this study | CF | 5 x 5 x 5 | training, validation |
| 2 | HEK293A interphase | this study | CF | 5 x 5 x 5 | training, validation |
| 3 | BSC-1 interphase | this study | CF | 5 x 5 x 5 | validation |
| 6 | SUM 159 interphase | this study | CF | 5 x 5 x 5 | validation |
| 8 | SUM 159 prometaphase | this study | CF | 10 x 10 x 10 | prediction |
| 9 | SUM 159 interphase | this study | CF | 10 x 10 x 10 | prediction |
| 12 | HeLa interphase | this study | HPFS | 5 x 5 x 5 | training, prediction |
| 13 | HeLa interphase | this study | HPFS | 5 x 5 x 5 | training, prediction |
| 13a | SVG-A interphase | this study | HPFS | 5 x 5 x 5 | training, prediction |
| 15 | SVG-A interphase | this study | HPFS | 5 x 5 x 5 | prediction |
| 16 | SVGA interphase | this study | HPFS | 5 x 5 x 5 | prediction |
| 17 | SVG-A interphase | this study | HPFS | 5 x 5 x 5 | prediction |
| 19 | HeLa-2 interphase | COSEM HHMI/Janelia | HPFS | 4 x 4 x 5.2 | training, validation |
| 20 | HeLa-3 interphase | COSEM HHMI/Janelia | HPFS | 4 x 4 x 3.2 | training, validation |
| 21 | Jurkat-1 interphase | COSEM HHMI/Janelia | HPFS | 4 x 4 x 3.4 | validation |
| 22 | Macrophage-2 interphase | COSEM HHMI/Janelia | HPFS | 4 x 4 x 3.4 | validation |

Description for each sample includes cell type, stage during cell cycle, fixation protocol (CF or HPFS), FIB-SEM resolution and use of the ground truth annotations for model training, validation or prediction.

#### Table S3. Types of data augmentation used in this study

| Augmentation | Parameter | Value |
|---|---|---|
| Mirror | axes | x, y, z |
| Transpose | axes | x, y, z |
| Elastic | control point spacing | 32, 32, 32 |
| | jitter sigma | 2, 2, 2 |
| | subsample | 4 |
| Rotation | axes | x, y |
| Mirror | axes | x, y, z |
| Transpose | axes | x, y, z |
| Intensity | scale | in [0.85, 1.15] |
| | shift | in [-0.15, 0.15] |

Type of augmentation and parameters to transform the ground truth annotations used during model training. Their detailed description is found in GUNPOWDER (http://funkey.science/gunpowder/api.html#augmentation-nodes). Since the rotation operation was performed only in two dimensions, it was necessary to perform the mirror and transpose operations twice in order to obtain all possible 3D orientations of a training block.

#### Table S4. Procedures used to generate ground truth annotations

| Cell | Structure | Method used to generate ground truth | Voxel Size (nm) (x, y, z) | Training Block Size ($um^3$) ROI (voxels) | Validation Block Size ($um^3$) ROI (voxels) |
|---|---|---|---|---|---|
| 1 | Mitochondria | Ilastik | 5 x 5 x 5 | 57.8 $um^3$<br>1200 x 700 x 550 | 22.5 $um^3$<br>600 x 400 x 750 |
| 2 | Mitochondria | Ilastik, VAST | 5 x 5 x 5 | 80.4 $um^3$<br>650 x 900 x 1100 | 15.3 $um^3$<br>450 x 800 x 340 |
| 3 | Mitochondria | Ilastik, VAST | 5 x 5 x 5 | - | 35.4 $um^3$<br>700 x 540 x 750 |
| 6 | Mitochondria | Ilastik, VAST | 5 x 5 x 5 | - | 7.6 $um^3$<br>241 x 476 x 528 |
| 1 | ER | GC, VAST | 5 x 5 x 5 | 59.7 $um^3$<br>600 x 590 x 1350 | 11.1 $um^3$<br>600 x 590 x 250 |
| 2 | ER | GC | 5 x 5 x 5 | 14.8 $um^3$<br>500 x 395 x 600 | 4.6 $um^3$<br>300 x205 x 600 |
| 3 | ER | GC | 5 x 5 x 5 | - | 2.1 $um^3$<br>204 x 204 x 400 |
| 6 | ER | GC, VAST | 5 x 5 x 5 | - | 7.5 $um^3$<br>241 x 476 x 528 |
| 1 | Golgi | GC | 5 x 5 x 5 | 47.1 $um^3$<br>469 x 650 x 510<br>400 x 400 x 875<br>350 x 400 x 400<br>230 x 250 x 440 | 5.0 $um^3$<br>250 x 400 x 400 |
| 2 | Golgi | GC | 5 x 5 x 5 | 9.4 $um^3$<br>300 x 500 x 500 | 5.2 $um^3$<br>230 x 400 x 450 |
| 3 | Golgi | GC, VAST | 5 x 5 x 5 | - | 2.8 $um^3$<br>210 x 283 x 371 |
| 6 | Golgi | GC, VAST | 5 x 5 x 5 | - | 3.1 $um^3$<br>284 x 204 x 424 |
| 12 | Clathrin coated pit | VAST | 5 x 5 x 5 | 1 $um^3$<br>6 [110 x 110 x 110] | .33 $um^3$<br>2 [110 X 110 X 110] |
| 13 | Clathrin coated pit | VAST | 5 x 5 x 5 | 1.5 $um^3$<br>9 [110 x 110 x 110] | .33 $um^3$<br>2 [110 X 110 X 110] |
| 13 | ER | GC, VAST | 5 x 5 x 5 | 2.03 $um^3$<br>110 x 290 x 510 | - |
| 13 | Mitochondria | Ilastik, VAST | 5 x 5 x 5 | 1 $um^3$<br>200 x 200 x 200 | - |
| 13a | Nuclear pore | VAST | 5 x 5 x 5 | 1.33 $um^3$<br>8 [110 x 110 x 110] | .33 $um^3$<br>2 [110 x 110 x 110] |

| | | | | | |
|---|---|---|---|---|---|
| 20 | Mitochondria | OpenOrganelle (Heinrich et al., 2021; Xu et al., 2021) | 4 x 4 x 3.2 | 4.51 $um^3$<br>500 x 250 x 500<br>200 x 200 x 200 | 0.52 $um^3$<br>200 x 200 x 200 |
| 21 | Mitochondria | OpenOrganelle (Heinrich et al., 2021; Xu et al., 2021) | 4 x 4 x 3.4 | - | 1.07 $um^3$<br>256 x 256 x 256 |
| 22 | Mitochondria | OpenOrganelle (Heinrich et al., 2021; Xu et al., 2021) | 4 x 4 x 3.4 | - | 0.22 $um^3$<br>150 x 150 x 150 |
| 19 | ER | OpenOrganelle (Heinrich et al., 2021; Xu et al., 2021) | 4 x 4 x 5.2 | 9.96 $um^3$<br>250 x 500 x 500<br>200 x 200 x 200<br>250 x 250 x 250<br>200 x 200 x 200<br>250 x 400 x 400<br>238 x 300 x 300 | 1 $um^3$<br>250 x 250 x 250 |
| 20 | ER | OpenOrganelle (Heinrich et al., 2021; Xu et al., 2021) | 4 x 4 x 3.2 | 4.51 $um^3$<br>500 x 250 x 500<br>200 x 200 x 200 | 1.07 $um^3$<br>256 x 256 x 256 |
| 21 | ER | OpenOrganelle (Heinrich et al., 2021; Xu et al., 2021) | 4 x 4 x 3.4 | - | 4 $um^3$<br>500 x 250 x 500 |
| 22 | ER | OpenOrganelle (Heinrich et al., 2021; Xu et al., 2021) | 4 x 4 x 3.4 | - | 4.02 $um^3$<br>501 x 250 x 502 |

Description of cells, procedures used to generate ground truth annotations (see methods for details), resolution of FIB-SEM data and size of hold out volumes used for training and validation.

#### Table S5. Effect of CLAHE on prediction performance

| | Mitochondria | | | | Golgi | | | | ER | | | |
|---|---|---|---|---|---|---|---|---|---|---|---|---|
| **Validation blocks in cell** | ***1*** | ***2*** | ***3 naive*** | ***6 naive*** | ***1*** | ***2*** | ***3 naive*** | ***6 naive*** | ***1*** | ***2*** | ***3 naive*** | ***6 naive*** |
| **without normaliza-tion** | 0.98± 0.01 | **0.92± 0.02** | 0.73± 0.04 | 0.88± 0.02 | 0.69± 0.03 | 0.68± 0.04 | 0.55± 0.02 | **0.80± 0.02** | 0.93± 0.01 | 0.87± 0.04 | 0.91± 0.01 | 0.83± 0.02 |
| **Linear rescale** | 0.96± 0.01 | **0.92± 0.03** | 0.64± 0.07 | **0.90± 0.02** | 0.68± 0.05 | 0.71± 0.04 | 0.51± 0.02 | 0.74± 0.05 | **0.96± 0.01** | **0.92± 0.01** | 0.91± 0.01 | **0.87± 0.01** |
| **histogram equaliza-tion** | 0.96± 0.02 | 0.85± 0.04 | 0.71± 0.05 | 0.84± 0.06 | 0.61± 0.08 | **0.74± 0.17** | 0.64± 0.06 | 0.36± 0.10 | 0.90± 0.02 | 0.87± 0.03 | 0.91± 0.02 | 0.83± 0.03 |
| **CLAHE clip 1%** | 0.97± 0.01 | **0.92± 0.02** | 0.73± 0.03 | 0.87± 0.01 | **0.70± 0.04** | 0.67± 0.03 | 0.52± 0.02 | 0.77± 0.03 | 0.95± 0.01 | 0.91± 0.01 | 0.91± 0.01 | 0.85± 0.01 |
| **CLAHE clip 2%** | 0.96± 0.02 | 0.87± 0.03 | 0.75± 0.05 | 0.88± 0.03 | 0.56± 0.08 | 0.69± 0.05 | 0.56± 0.04 | 0.79± 0.02 | 0.95± 0.01 | 0.89± 0.04 | **0.92± 0.01** | 0.77± 0.04 |
| **CLAHE clip 3%** | **0.99± 0.00** | 0.89± 0.02 | **0.81± 0.03** | 0.89± 0.01 | 0.68± 0.15 | 0.69± 0.12 | **0.66± 0.06** | 0.37± 0.25 | 0.91± 0.02 | 0.84± 0.06 | 0.91± 0.02 | 0.72± 0.05 |

F1 prediction scores using the indicated cells were obtained with models trained with combined FIB-SEM data from cells 1 and 2 subjected to the indicated types of signal normalization. F1 score for each organelle corresponds to the average +/- SD from 20 consecutive predictions obtained every 1000 iterations initiated after about 150,000 training iterations. Best results are highlighted in bold. Inspection of the data shows no consistent improvement in the F1 predictions scores upon signal normalization including CLAHE.

#### Table S6. Comparative examples of predictive performance by models trained with data from one or two cells

| Structure Fixation | F1 for model trained using Cell 1 | | | | F1 for model trained using Cell 2 | | | | F1 for model trained using Cells 1+2 or Cells 19 + 20 | | | |
|---|---|---|---|---|---|---|---|---|---|---|---|---|
| ***Cells used for predictions*** | ***1*** | ***2 naïve*** | ***3 naïve*** | ***6 naïve*** | ***1 naïve*** | ***2*** | ***3 naïve*** | ***6 naïve*** | ***1*** | ***2*** | ***3 naïve*** | ***6 naïve*** |
| **Mitochondria CF** | 0.91 | 0.47 | 0.66 | 0.81 | 0.89 | 0.87 | 0.74 | 0.7 | 0.96 | 0.87 | 0.75 | 0.88 |
| **ER CF** | 0.86 | 0.29 | 0.7 | 0.52 | 0.16 | 0.90 | 0.83 | 0.55 | 0.95 | 0.90 | 0.92 | 0.77 |
| **Golgi CF** | 0.45 | 0.68 | 0.61 | 0.73 | 0.18 | 0.62 | 0.48 | 0.77 | 0.56 | 0.69 | 0.56 | 0.79 |
| ***Cells used for predictions*** | | | | | | | | | ***19*** | ***20*** | ***21 naïve*** | ***22 naïve*** |
| **Mitochondria HPFS** | | | | | | | | | 1 | 1 | 0.94 | 0.93 |
| **ER HPFS** | | | | | | | | | 0.91 | 0.80 | 0.48 | 0.81 |

A neural network was trained using ground truth annotations from the indicated cells, alone or in combination, and the resulting models then used to predict on images of the listed individual cells. The data show F1 prediction scores using validation datasets not employed for training.

#### Table S7. Comparison of model performance using the ASEM (this study) and COSEM pipelines

| **Source** | **Structure**<br>**Fixation** | **Post Processing** | **Training Model #**<br>**Training iterations (x1000)** | | | | **F1 for the indicated cells used for training** | | **F1 for naive cells** | | **Fine-tuning Training Iterations (x1000)** | | **F1 after fine-tuning** | |
|---|---|---|---|---|---|---|---|---|---|---|---|---|---|---|
| ***Cells training and predictions*** | | | **19 + 20** | | | | **19** | **20** | **21** | **22** | **21** | **22** | **21** | **22** |
| this study | Mitochondria<br>HPFS | No | 1675<br>95-115 | | | | 0.99 | 0.99 | 0.94 | 0.93 | 2-7 | 2-7 | 0.93 | 0.98 |
| this study | ER<br>HPFS | No | 1669<br>180-200 | | | | 0.91 | 0.80 | 0.48 | 0.81 | 7-12 | 1-6 | 0.69 | 0.90 |
| | | | | | | | **F1 for the indicated cells used for training** | | | | | | | |
| ***Cells training and predictions*** | | | **19** | **20** | **21** | **22** | **19** | **20** | **21** | **22** | | | | |
| COSEM | Mitochondria<br>HPFS | Yes | Few<br>575 | All<br>825 | All<br>875 | Many<br>110 | 0.93 | 0.97 | 0.98 | N/A | - | - | - | - |
| COSEM | ER<br>HPFS | Yes | Many<br>625 | All<br>1075 | Few<br>625 | Many<br>650 | 0.84 | 0.71 | 0.75 | 0.97 | - | - | - | - |

The neural networks used in this study or by the COSEM project (Heinrich et al., 2021) were trained to predict mitochondria and ER using ground truth annotations from the indicated cells, alone (COSEM project) or in combination (this study). The data show F1 prediction scores using validation datasets not employed for training. In this study we only used data from one organelle at a time to train a neural network and report the results for the defined number of training iterations chosen when the performance of the model reached stability; Reported F1 scores are the average of 20 consecutive values obtained every 1,000 training iterations determined after the indicated training iteration. The COSEM project simultaneously used data from more than one organelle to train the neural network according to the details described in (Heinrich et al., 2021). The F1 scores for the COSEM Project reported in Tables 1 & 2 (Heinrich et al., 2021) correspond to their best results by different trained networks and training iterations using ground annotations for ‘few’, ‘many’ or ‘all’ organelles including mitochondria and ER present within the hold-out block.

**Table S8. Comparative examples of predictive performance by models trained with data from cells prepared with the same or different fixation protocols**

| Structure | Training iterations (x1000) | F1 for hold-out blocks of indicated training cells | F1 for naive cells | Fine-tuning training iterations (x1000) | F1 after fine-tuning with naïve cells |
|---|---|---|---|---|---|
| ***Cells used for training & predictions*** | ***1, 2 (CF)*** | ***1 (CF)<br>2 (CF)*** | | | |
| ***Cells used for predictions*** | | | ***3 (CF)<br>6 (CF)*** | ***3 (CF)<br>6 (CF)*** | ***3 (CF)<br>6 (CF)*** |
| **Mitochondria** | 95-115 | 0.96<br>0.87 | 0.75<br>0.88 | 1-6<br>1-6 | 0.88<br>0.89 |
| **ER** | 115-135 | 0.95<br>0.90 | 0.92<br>0.77 | - | - |
| **Golgi** | 35-55 | 0.56<br>0.69 | 0.56<br>0.79 | - | - |
| ***Cells used for training & predictions*** | ***19, 20 (HPFS)*** | ***19 (HPFS)<br>20 (HPFS)*** | | | |
| ***Cells used for predictions*** | | | ***21 (HPFS)<br>22 (HPFS)*** | ***21 (HPFS)<br>22 (HPFS)*** | ***21 (HPFS)<br>22 (HPFS)*** |
| **Mitochondria** | 95-115 | 0.99<br>0.99 | 0.94<br>0.93 | 2-7<br>2-7 | 0.93<br>0.98 |
| **ER** | 180-200 | 0.91<br>0.80 | 0.48<br>0.81 | 7-12<br>1-6 | 0.69<br>0.90 |
| ***Cells used for training & predictions*** | ***1,2 (CF)<br>21, 22 (HPFS)*** | ***1 (CF)<br>2 (CF)<br>21 (HPFS)<br>22 (HPFS)*** | | | |
| ***Cells used for predictions*** | | | ***3 (CF)<br>6 (CF)<br>21 (HPFS)<br>22 (HPFS)*** | ***3 (CF)<br>6 (CF)<br>21 (HPFS)<br>22 (HPFS)*** | ***3 (CF)<br>6 (CF)<br>21 (HPFS)<br>22 (HPFS)*** |
| **Mitochondria** | 135-155 | 0.95<br>0.81<br>0.93<br>0.99 | 0.73<br>0.77<br>0.96<br>0.89 | -<br>1-6<br>-<br>- | -<br>0.90<br>-<br>- |
| **ER** | 100-120 | 0.94<br>0.90<br>0.84<br>0.74 | 0.85<br>0.82<br>0.58<br>0.81 | -<br>-<br>1-6<br>- | -<br>-<br>0.68<br>- |

| | | | |
|---|---|---|---|
| ***Cells used for training & predictions*** | ***13a (HPFS)*** | ***13a (HPFS)*** | |
| **Nuclear pores** | 130-150 | 0.52 | - |
| ***Cells used for training & predictions*** | ***13 (HPFS)*** | ***13 (HPFS)*** | |
| ***Cells used for predictions*** | | | ***12 (HPFS)*** |
| **Clathrin-coated pits/ vesicles** | 80-100 | 0.67 | 0.69 |

The neural network was trained using ground truth annotations from the indicated cells prepared with different fixation protocols, alone or in combination, and the resulting models then used to predict on images of the individual cells listed in the table. The data show F1 prediction scores using validation datasets not employed for training.

#### Table S9. Effect of resolution on the predictive performance

| Structure | F1 for hold-out blocks of indicated training cells @ 5 nm (from Table S8) | F1 for hold-out blocks of indicated training cells @ 10 nm | F1 for naïve cells @ 5 nm (from Table S8) | F1 for naïve cells @ 10 nm |
|---|---|---|---|---|
| ***Cells used for training & predictions*** | ***1 (CF)<br>2 (CF)*** | ***1 (CF)<br>2 (CF)*** | | |
| ***Cells used for predictions*** | | | ***3 (CF)<br>6 (CF)*** | ***3 (CF)<br>6 (CF)*** |
| **Mitochondria** | 0.96<br>0.87 | 0.97<br>0.79 | 0.75<br>0.88 | 0.62<br>0.86 |
| **ER** | 0.95<br>0.90 | 0.94<br>0.90 | 0.92<br>0.77 | 0.86<br>0.81 |
| **Golgi** | 0.56<br>0.69 | 0.45<br>0.59 | 0.56<br>0.79 | 0.38<br>0.58 |

The neural network was trained using ground truth annotations for the organelles from the indicated cells prepared by chemical fixation imaged at 5 nm (Data from Table S8). The resulting models were used to predict on the individual cells listed in the table. The same ground truth annotations were also isotropically downsampled to 10 nm, to then train new models used to predict on images of the same cells, but downsampled to 10 nm. The data show F1 prediction scores using validation datasets not employed for training.

#### Table S10. Summary of experiments to test the effect of fine-tuning

| Structure<br><br>Fixation protocol | Training cells<br><br>ground-truth volume [um] | Training iterations (x1000) | Cell<br><br>F1 prediction | Cell<br><br>fine-tuning ground-truth volume [um] | Cell<br><br>Fine-tuning Iterations (x1000) | Cell<br><br>F1 after fine-tuning |
|---|---|---|---|---|---|---|
| **Mitochondria<br>CF** | 1 + 2<br>138.2 μm³ | 95 | 3, naïve<br>0.75<br><br>6, naïve<br>0.88 | 3, naïve<br>1.95 μm³<br><br>6, naïve<br>1.15 μm³ | 3, naïve<br>6<br><br>6, naïve<br>6 | 3, naïve<br>0.88<br><br>6, naïve<br>0.89 |
| **Mitochondria<br>HPFS** | 19 + 20<br>12.12 μm³ | 95 | 21, naïve<br>0.94<br><br>22, naïve<br>0.93 | 21, naïve<br>4.00 μm³<br><br>22, naïve<br>4.02 μm³ | 21, naïve<br>7<br><br>22, naïve<br>7 | 21, naïve<br>0.93<br><br>22<br>0.98 |
| **ER<br>HPFS** | 19 + 20<br>4.85 μm³ | 180 | 21, naïve<br>0.48<br><br>22, naïve<br>0.81 | 21, naïve<br>1.07 μm³<br><br>22, naïve<br>0.52 μm³ | 21, naïve<br>12<br><br>22, naïve<br>6 | 21, naïve<br>0.69<br><br>22, naïve<br>0.90 |

Sheridan, A., Nguyen, T., Deb, D., Lee, W.-C.A., Saalfeld, S., Turaga, S., Manor, U., and Funke, J. (2022). Local Shape Descriptors for Neuron Segmentation. Biorxiv 2021.01.18.427039.

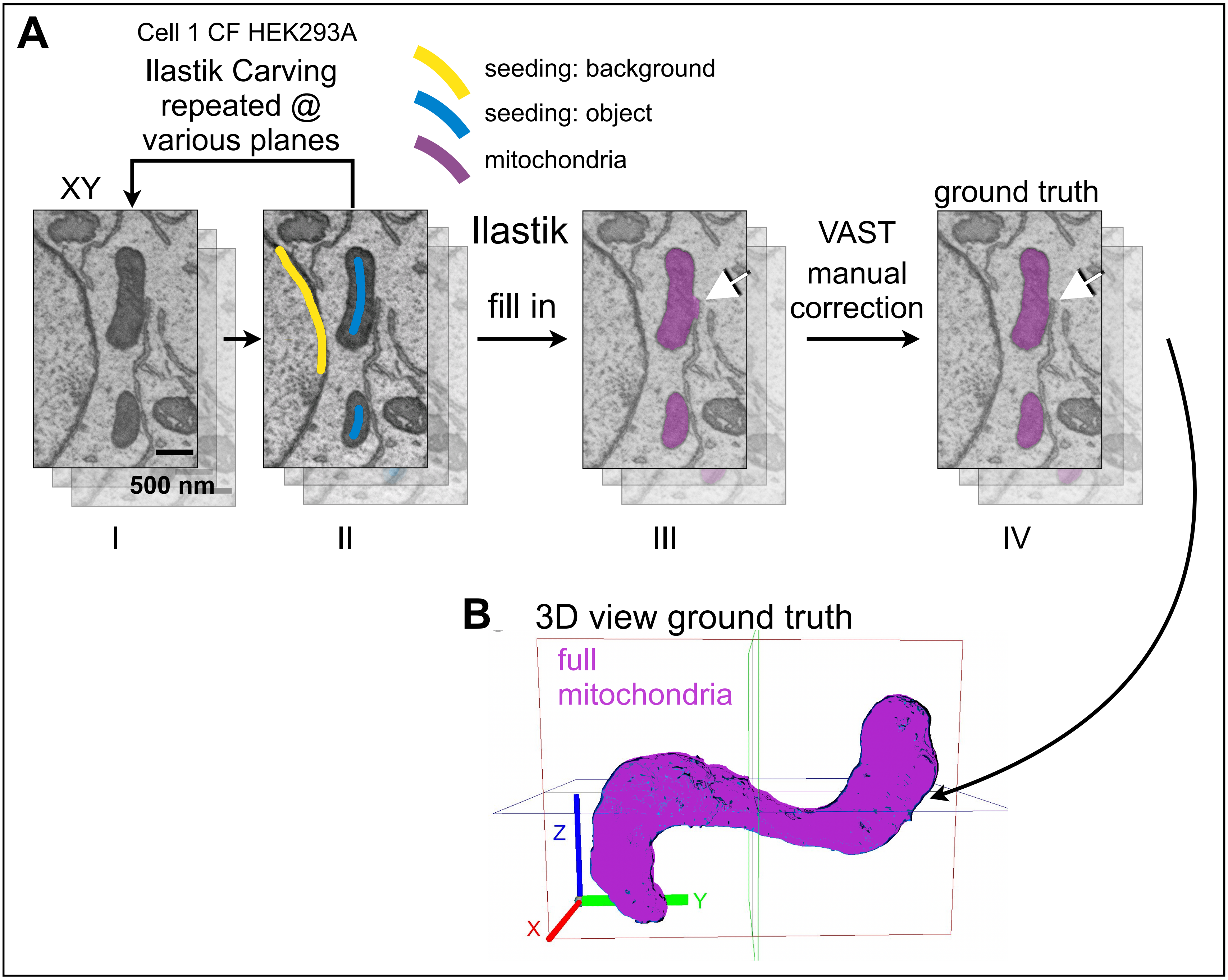
A
Cell 1 CF HEK293A
Ilastik Carving repeated @ various planes
seeding: background
seeding: object
mitochondria
XY
500 nm
Ilastik
fill in
VAST
manual
correction
ground truth
I
II
III
IV
B
3D view ground truth
full mitochondria
Z
Y
X

### Figure S1

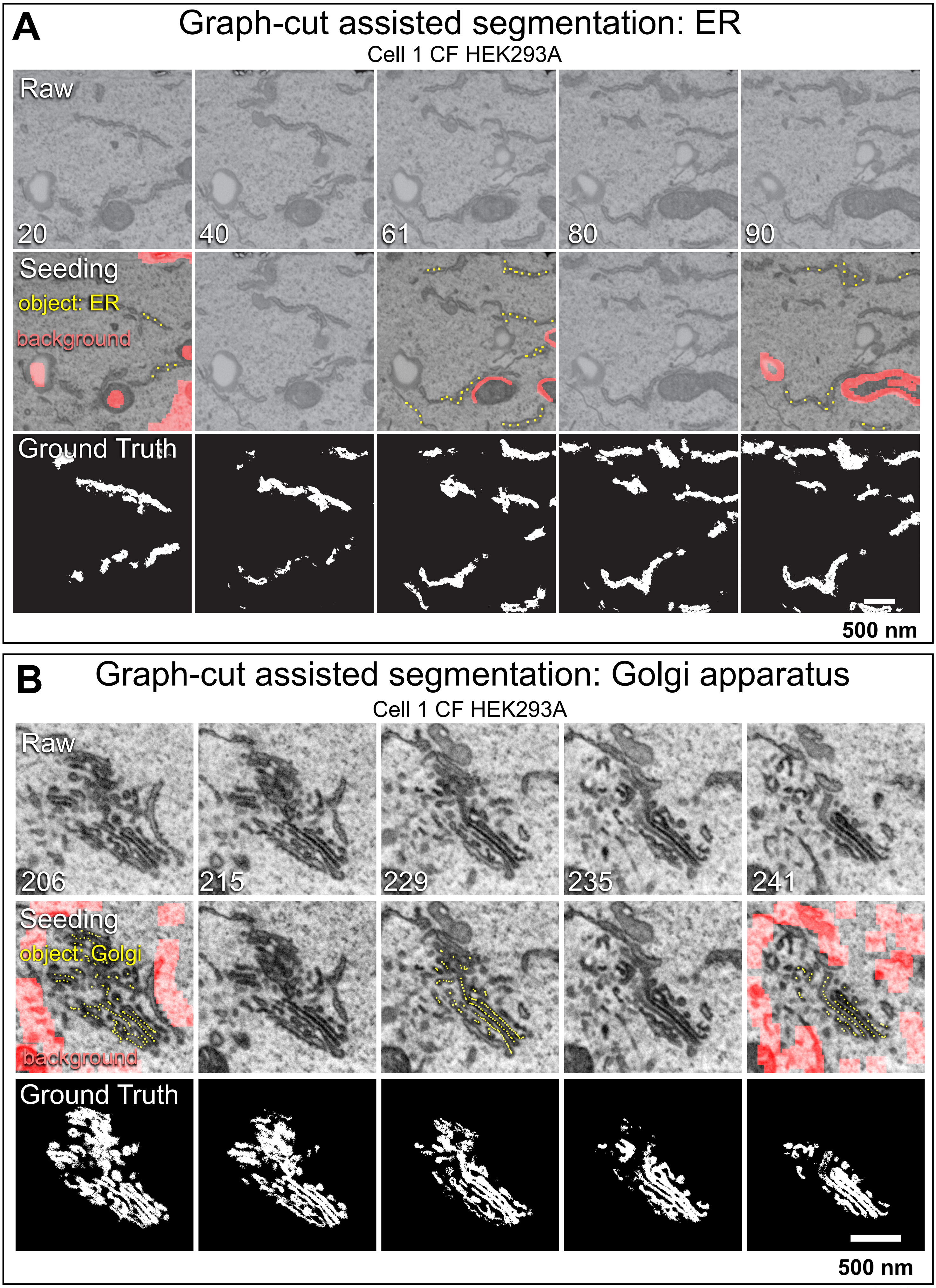

A
Graph-cut assisted segmentation: ER
Cell 1 CF HEK293A
Raw
20
40
61
80
90
Seeding
object: ER
background
Ground Truth
500 nm
B
Graph-cut assisted segmentation: Golgi apparatus
Cell 1 CF HEK293A
Raw
206
215
229
235
241
Seeding
object: Golgi
background
Ground Truth
500 nm

Figure S2

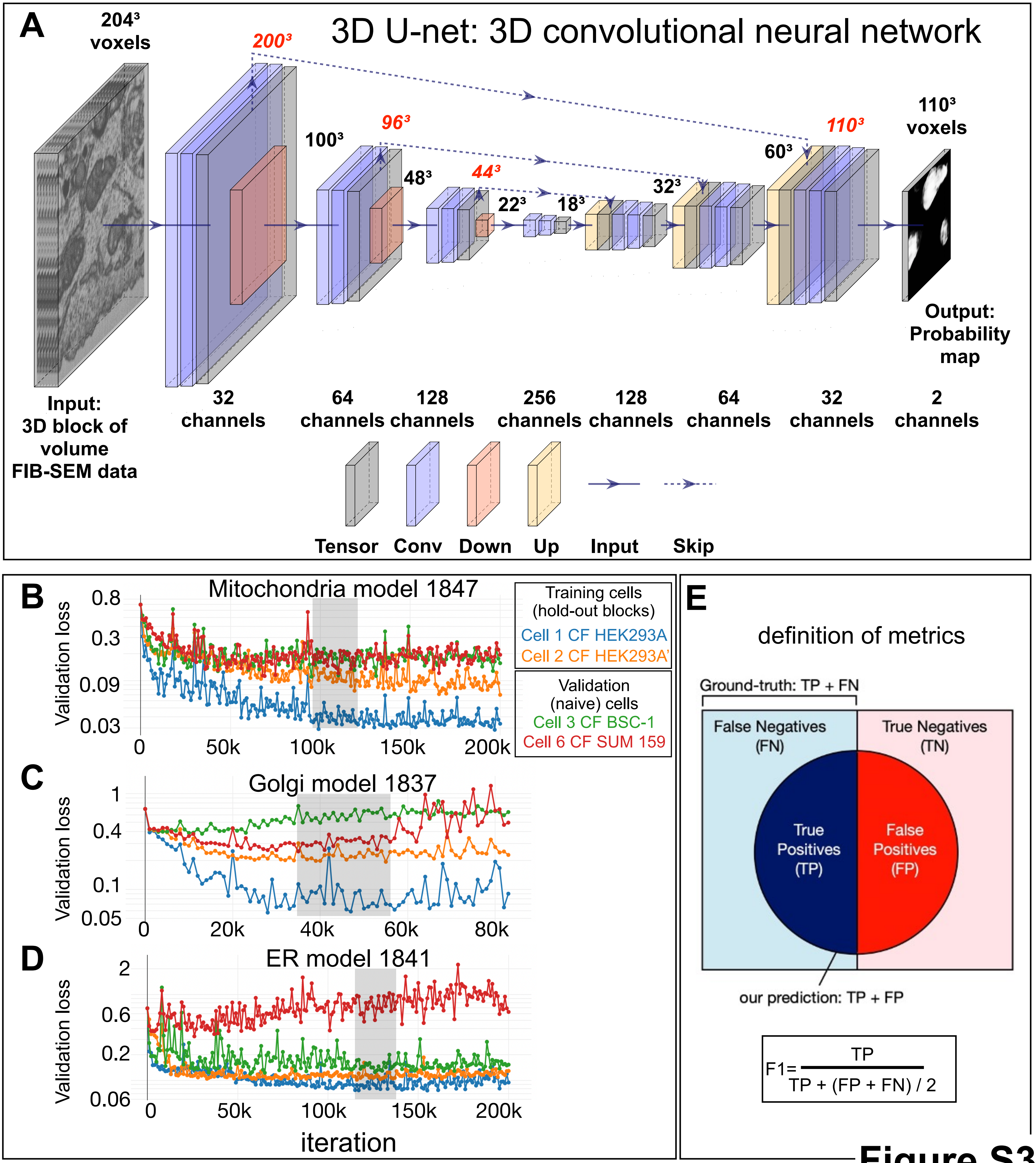
A
3D U-net: 3D convolutional neural network
204³ voxels
200³
100³
96³
48³
44³
22³
18³
32³
60³
110³
110³ voxels
Output: Probability map
Input: 3D block of volume FIB-SEM data
32 channels
64 channels
128 channels
256 channels
128 channels
64 channels
32 channels
2 channels
Tensor
Conv
Down
Up
Input
Skip
B
Mitochondria model 1847
Validation loss
0.8
0.3
0.09
0.03
0
50k
100k
150k
200k
Training cells (hold-out blocks)
Cell 1 CF HEK293A
Cell 2 CF HEK293A'
Validation (naive) cells
Cell 3 CF BSC-1
Cell 6 CF SUM 159
C
Golgi model 1837
Validation loss
1
0.4
0.1
0.05
0
20k
40k
60k
80k
D
ER model 1841
Validation loss
2
0.6
0.2
0.06
0
50k
100k
150k
200k
iteration
E
definition of metrics
Ground-truth: TP + FN
False Negatives (FN)
True Negatives (TN)
True Positives (TP)
False Positives (FP)
our prediction: TP + FP
F1= TP / (TP + (FP + FN) / 2)

**Figure S3**

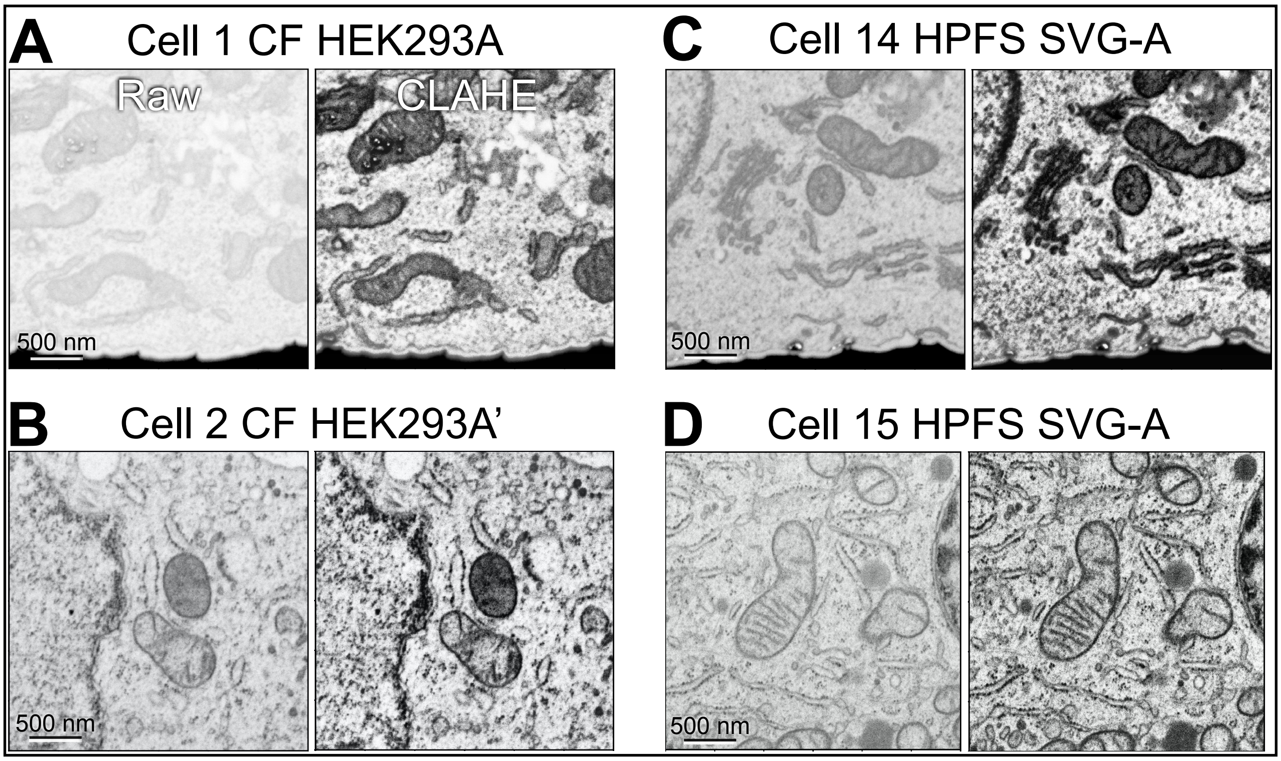
A
Cell 1 CF HEK293A
Raw
CLAHE
500 nm
B
Cell 2 CF HEK293A'
500 nm
C
Cell 14 HPFS SVG-A
500 nm
D
Cell 15 HPFS SVG-A
500 nm

### Figure S4

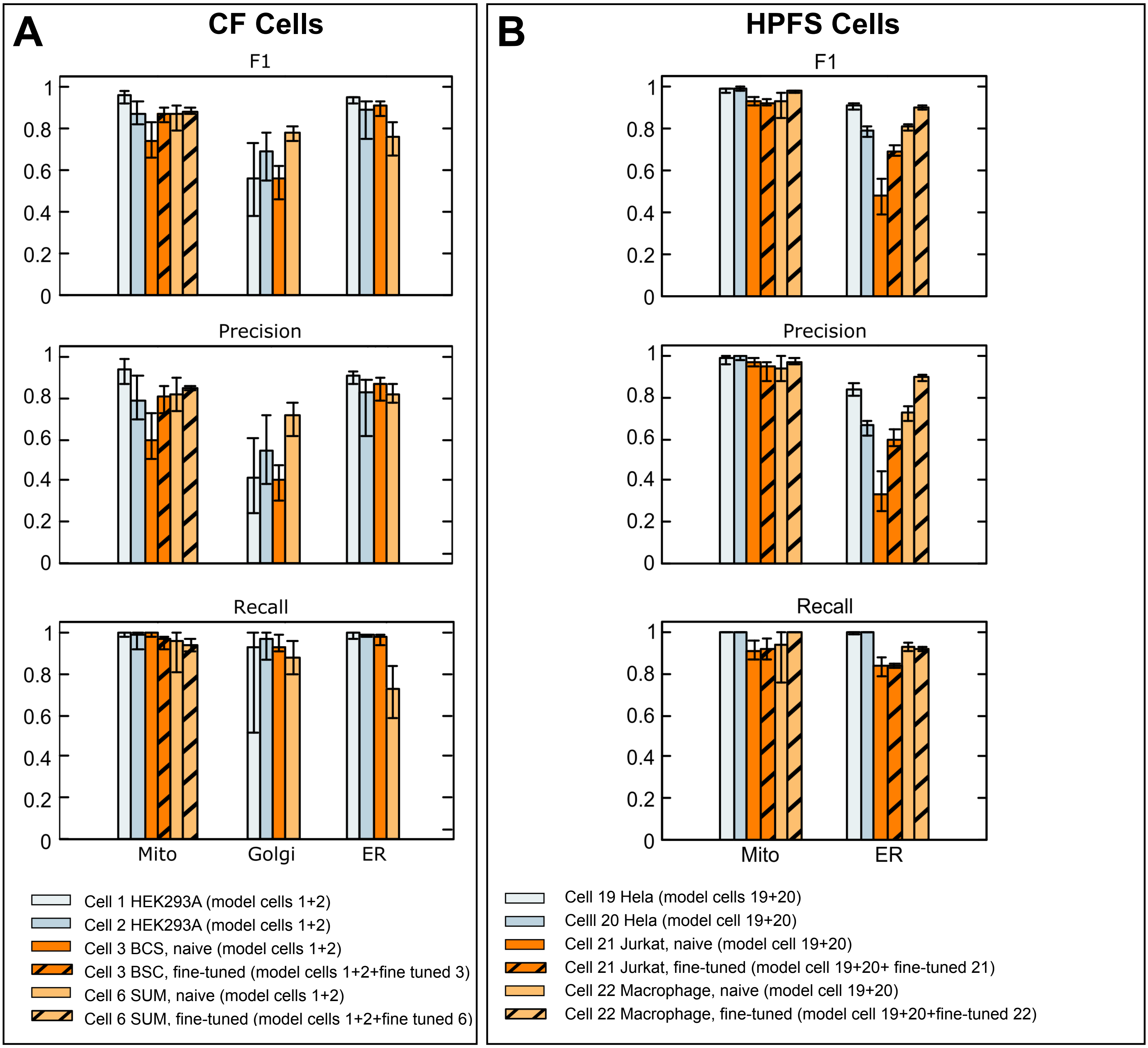
A
CF Cells
F1
Precision
Recall
Mito
Golgi
ER
Cell 1 HEK293A (model cells 1+2)
Cell 2 HEK293A (model cells 1+2)
Cell 3 BCS, naive (model cells 1+2)
Cell 3 BSC, fine-tuned (model cells 1+2+fine tuned 3)
Cell 6 SUM, naive (model cells 1+2)
Cell 6 SUM, fine-tuned (model cells 1+2+fine tuned 6)
B
HPFS Cells
F1
Precision
Recall
Mito
ER
Cell 19 Hela (model cells 19+20)
Celll 20 Hela (model cell 19+20)
Cell 21 Jurkat, naive (model cell 19+20)
Cell 21 Jurkat, fine-tuned (model cell 19+20+ fine-tuned 21)
Cell 22 Macrophage, naive (model cell 19+20)
Cell 22 Macrophage, fine-tuned (model cell 19+20+fine-tuned 22)

\

**Figure S5**

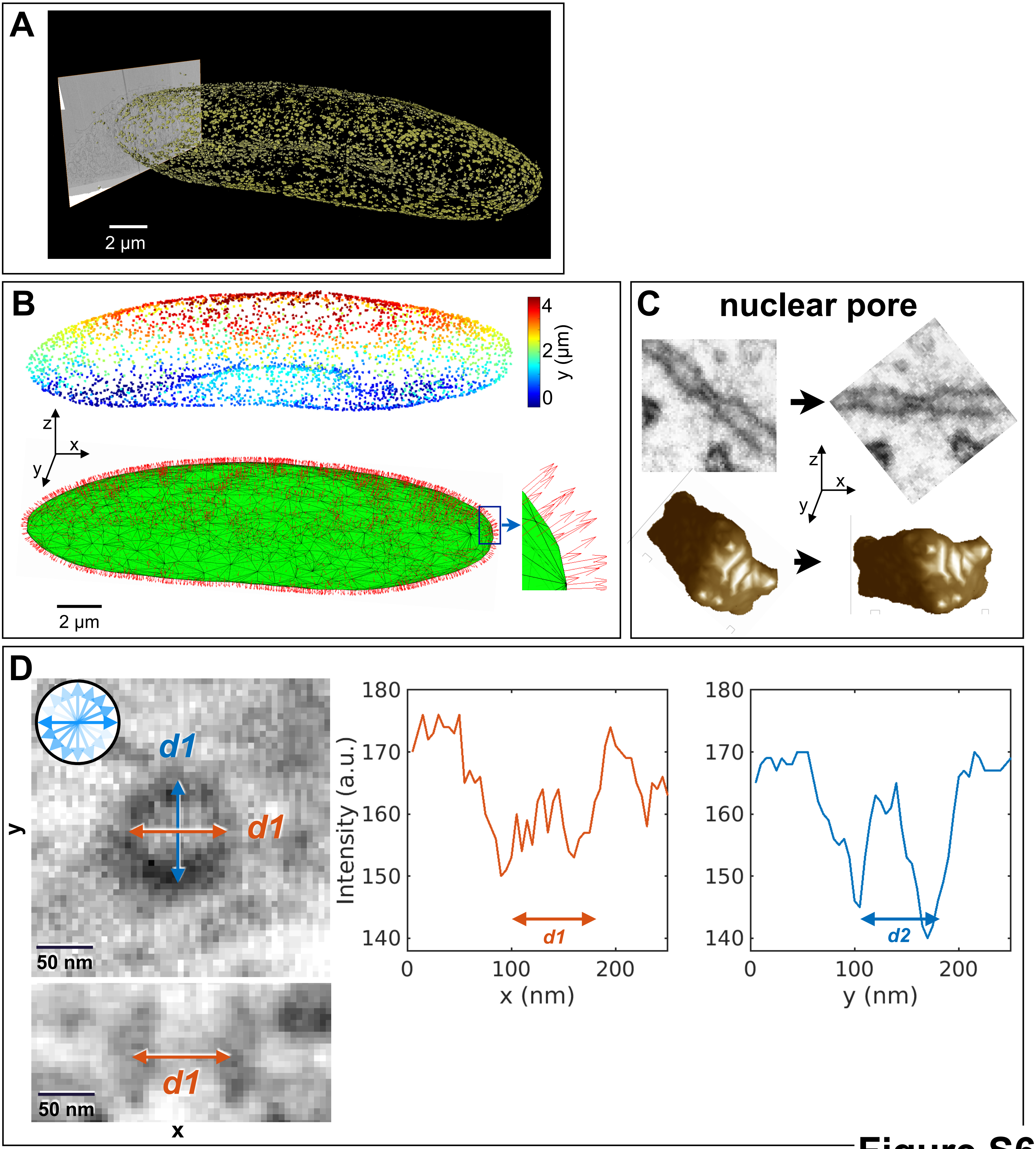
A
2 µm
B
4
2
0
y (µm)
z
x
y
2 µm
C
nuclear pore
z
x
y
D
d1
d1
y
50 nm
d1
50 nm
x
180
170
160
150
140
Intensity (a.u.)
0
100
200
x (nm)
d1
180
170
160
150
140
0
100
200
y (nm)
d2

Figure S6

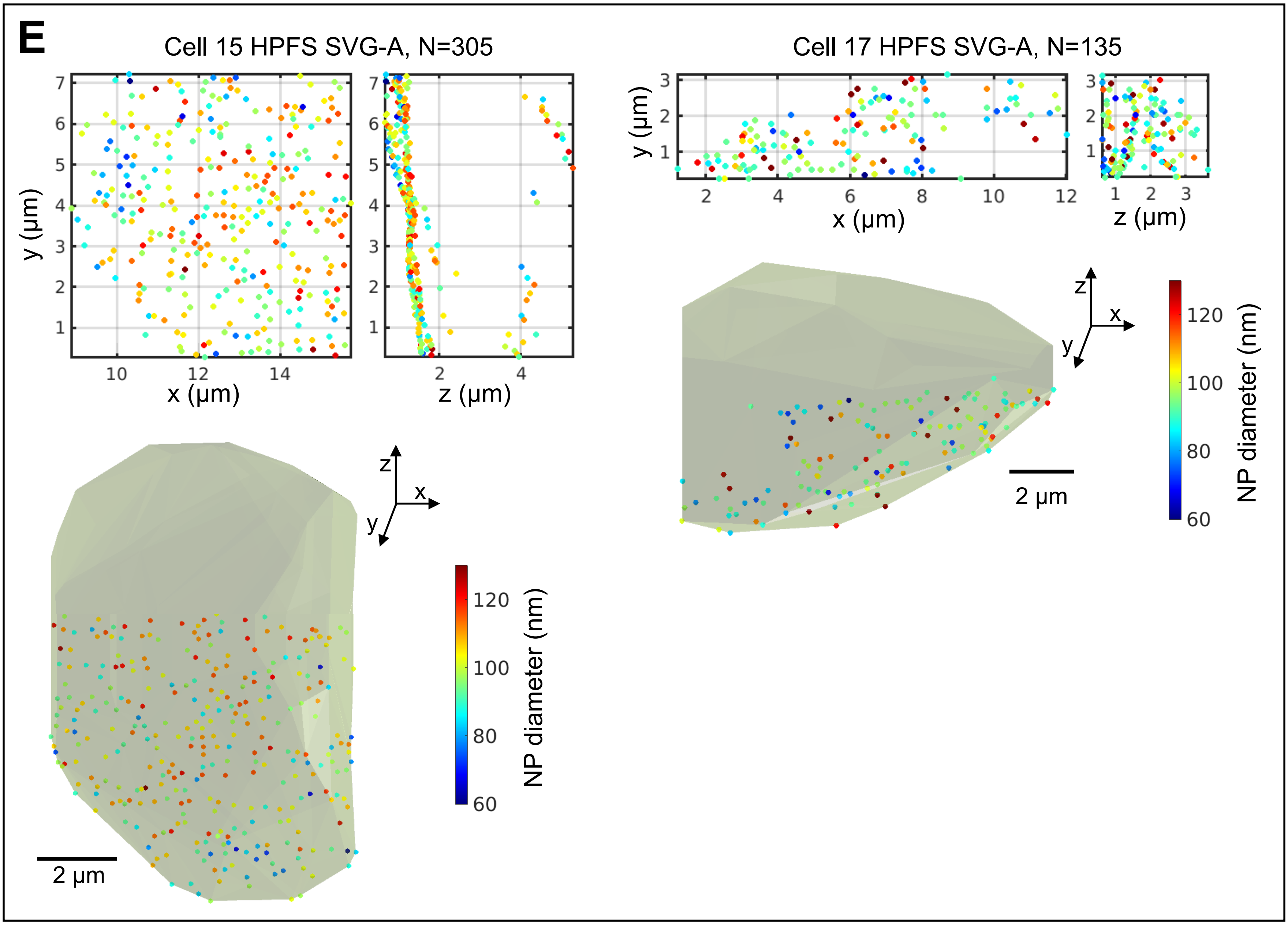
E
Cell 15 HPFS SVG-A, N=305
Cell 17 HPFS SVG-A, N=135
y (µm)
x (µm)
z (µm)
2 µm
NP diameter (nm)
60
80
100
120

**Figure S6**

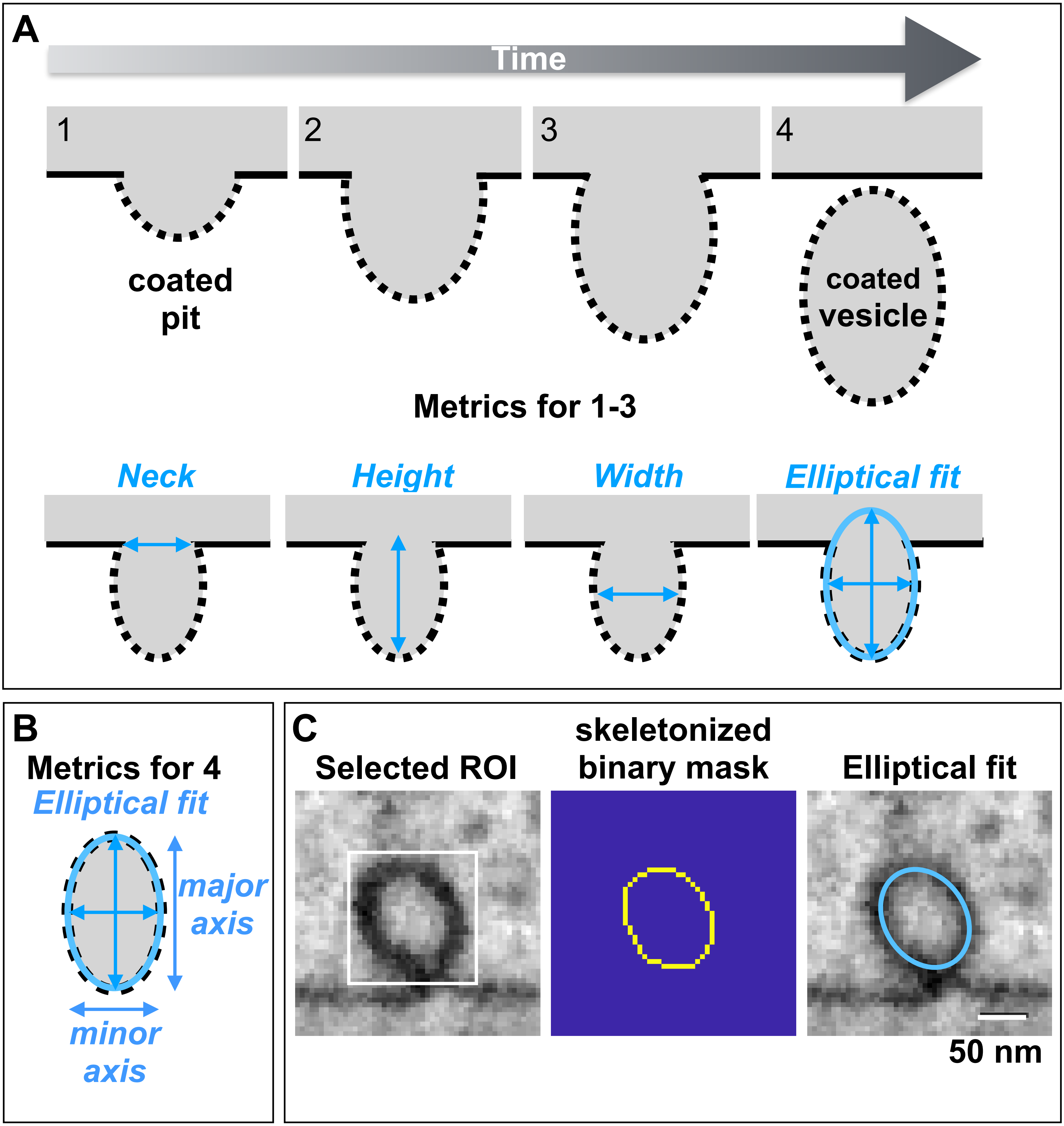
A
Time
1
2
3
4
coated
pit
coated
vesicle
Metrics for 1-3
Neck
Height
Width
Elliptical fit
B
Metrics for 4
Elliptical fit
major
axis
minor
axis
C
Selected ROI
skeletonized
binary mask
Elliptical fit
50 nm

**Figure S7**

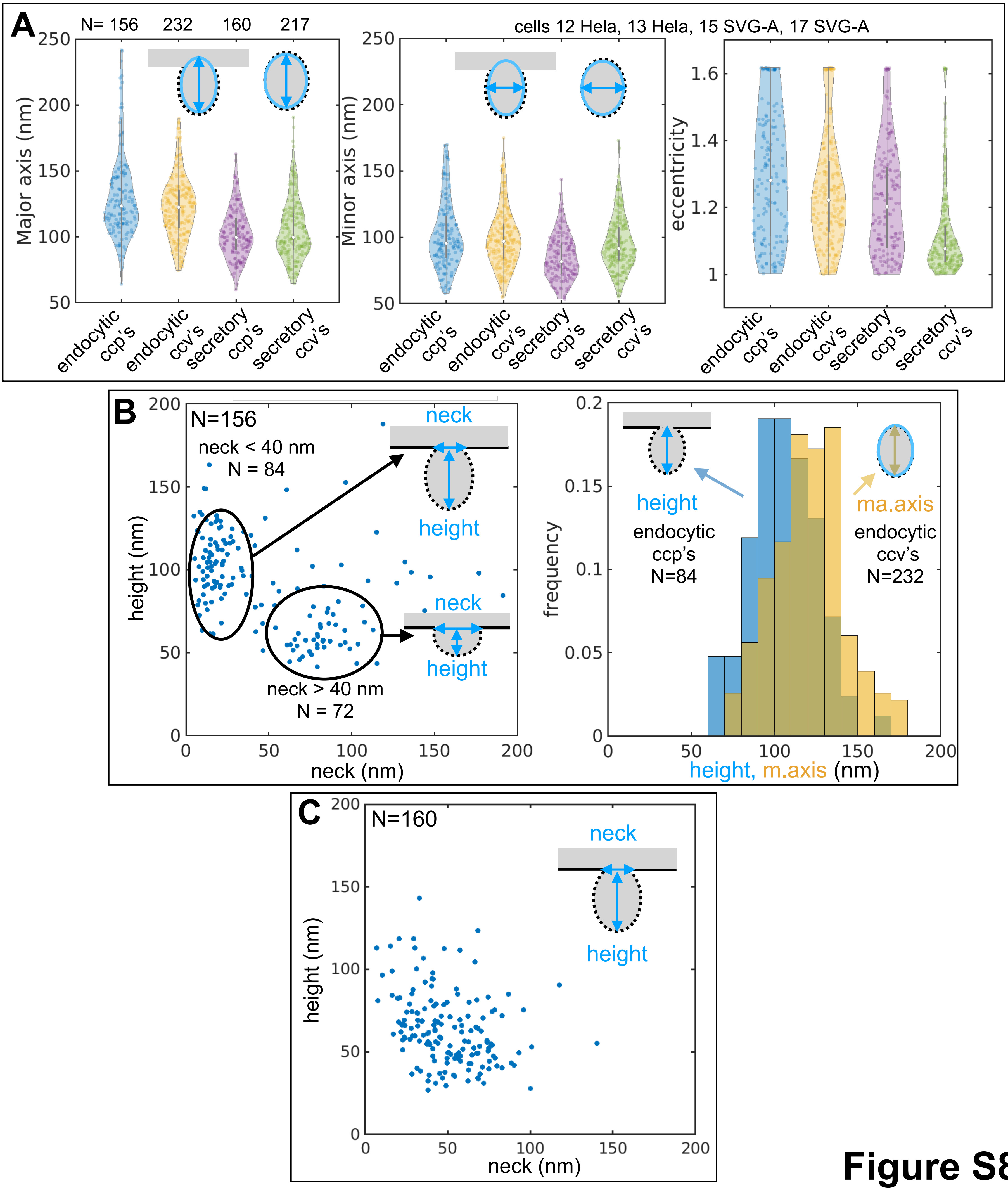
A
N= 156 232 160 217
cells 12 Hela, 13 Hela, 15 SVG-A, 17 SVG-A
Major axis (nm)
Minor axis (nm)
eccentricity
endocytic ccp's
endocytic ccv's
secretory ccp's
secretory ccv's
B
N=156
neck < 40 nm
N = 84
neck > 40 nm
N = 72
neck
height
height (nm)
neck (nm)
frequency
height
endocytic ccp's
N=84
ma.axis
endocytic ccv's
N=232
height, m.axis (nm)
C
N=160
neck
height
height (nm)
neck (nm)

Figure S8
